# Machine learning-based exploration, expansion and definition of the atropopeptide family of ribosomally synthesized and posttranslationally modified peptides

**DOI:** 10.1101/2023.11.03.565440

**Authors:** Friederike Biermann, Bin Tan, Milena Breitenbach, Pakjira Nanudorn, Yuya Kakumu, Yoana Dimitrova, Allison S. Walker, Reiko Ueoka, Eric J. N. Helfrich

**Affiliations:** Institute for Molecular Bio Science Goethe University Frankfurt, Max-von-Laue Strasse 9, 60438 Frankfurt am Main, Germany; LOEWE Center for Translational Biodiversity Genomics (TBG) Senckenberganlage 25, 60325 Frankfurt am Main, Germany; Department of Chemistry Vanderbilt University, Stevenson Center 7330, Nashville, TN 37240, USA; Department of Biological Sciences Vanderbilt University, VU Station B, Box 35-1634, Nashville, TN 37235, USA; School of Marine Biosciences Kitasato University 1-15-1 Kitasato, Minami-ku, Sagamihara, Kanagawa, 252-0373, Japan

## Abstract

Ribosomally synthesized and posttranslationally modified peptides (RiPPs) constitute a diverse class of natural products. Atropopeptides are a recent addition to the fast-growing number of RiPP families. Characterized members of the peptide family feature a particular intricate three-dimensional shape. Here we developed AtropoFinder, a machine learning-based algorithm to chart the biosynthetic landscape of the atropopeptides. AtropoFinder identified more than 650 atropopeptide biosynthetic gene clusters (BGCs). Through bioinformatics and modeling analyses, we pinpointed crucial motifs and residues in leader and core peptide sequences, prompting a refined definition of the atropopeptide RiPP family. Our study revealed that a substantial subset of atropopeptide BGCs harbors multiple tailoring genes, thus suggesting a broader structural diversity than previously anticipated. To verify AtropoFinder, we heterologously expressed four atropopeptide BGCs, which resulted in the identification of novel atropopeptides with varying peptide lengths, number and type of modifications. Most notably, our study resulted in the characterization of an atropopeptide that is more extensively modified than previously identified members, resulting in an even more rigid 3-dimensional shape. Moreover, one characterized atropopeptide BGC encoding a single P450 is involved in the biosynthesis of two peptides with the same sequence but distinct and non-overlapping modification patterns. This work expands the atropopeptide chemical space, advances our understanding of atropopeptide biosynthesis and underscores the potential of machine learning in uncovering the uncharted biosynthetic diversity encoded in RiPP biosynthetic blueprints.

## Introduction

Peptide natural products play a crucial role in drug discovery due to their diverse structures, high potency, selectivity, and strong affinity towards drug targets, as well as favorable pharmacokinetic properties.^1^ Examples of pharmaceutically relevant peptides include vancomycin (antibacterial), cyclosporin (immunosuppressive), bacitracin (antibacterial), and thiostrepton (antibacterial).^2–5^ Historically, complex peptide natural products were believed to be solely biosynthesized by non-ribosomal peptide synthetases (NRPSs).^6,7^ NRPSs are giant multi-enzyme complexes that assemble complex peptides in an assembly line-like fashion independent of the ribosome. However, over the past two decades, numerous complex peptides have been discovered that are biosynthesized by ribosomally synthesized and posttranslationally modified peptide biosynthetic pathways (RiPPs).^8^ The array of characterized posttranslational modifications in RiPP systems is constantly increasing. RiPPs encompass a heterogenous group of natural products, which can be subdivided into more than 40 families.^8^ The corresponding gene clusters can be distinguished by the presence of characteristic tailoring genes (e.g., linear polyazole-containing peptides), conserved precursor genes (e.g., the Nif11-like proteusin precursors) or the intricate structure of the associated peptide (e.g., lassopeptides).^9^ State-of-the-art genome mining tools frequently overlook RiPP BGCs that do not belong to the well-studied RiPP families.^10^ Consequently, RiPP-derived products are betimes initially misannotated as NRPS products.^7,11,12^ Extensive research into RiPPs over the past two decades has expanded the RiPP biosynthetic repertoire and revealed substantial overlap in the diverse array of modifications observed in peptides from RiPP and NRPS pathways (Figure S1).^7^ As a result, the annotation of complex peptides as nonribosomal peptides (NRPs) or RiPPs can often be challenging unless peptide class-defining modifications are present in the molecules.^7^ This challenge can be showcased by tryptorubin A which was initially associated with a NRPS BGC.^11^ The proposed BGC was the sole peptide biosynthetic gene cluster (BGC) detected by state-of-the-art genome mining platforms to be present in all three reported tryptorubin A producers.^11^ Tryptorubin A (**1**) is a hexapeptide that features an unusually complex three-dimensional structure.^13^ The extremely rigid three-dimensional shape arises from a biaryl and a carbon-nitrogen bridge between aromatic amino acid side chains and an additional carbon-nitrogen bridge between an aromatic amino acid side chain and the peptide backbone, respectively.^13^ These modifications result in two possible unusual atropisomeric configurations, only one of which has been found in nature.^13^ Unusual atropisomerism (recently also referred to as ansamerism^14^) is a type of stereoisomerism in which both isomers are theoretically interconvertible by multiple nonphysical bond torsions.^13^ Intrigued by the structural complexity of the hexapeptide, we revisited its biosynthesis. A combination of manual genome mining and heterologous expression studies unveiled that tryptorubin A is the first member of a new RiPP-family that we named atropopeptides (also referred to as atropitides^8^).^15^ The biosynthetic blueprint for the production of characterized atropopeptides resides within small BGCs containing only two genes: one encoding a precursor peptide and another a cytochrome P450 monooxygenase.^15^

The precursor peptide, comprising a leader and core peptide, is ribosomally synthesized. The leader peptide recruits the BGC-encoded cytochrome P450 which introduces the distinctive biaryl and carbon-nitrogen bridges in an atropospecific manner (Figure 1).^15^ In the final step, the modified core peptide is proteolytically released from the leader peptide. It is believed that an ubiquitous protease catalyzes the release of the mature hexapeptide, which can be further modified by the removal of the N-terminal alanine residue by another unknown peptidase that is not encoded in the BGC to yield the pentapeptide variant.^15^

**Figure 1:**
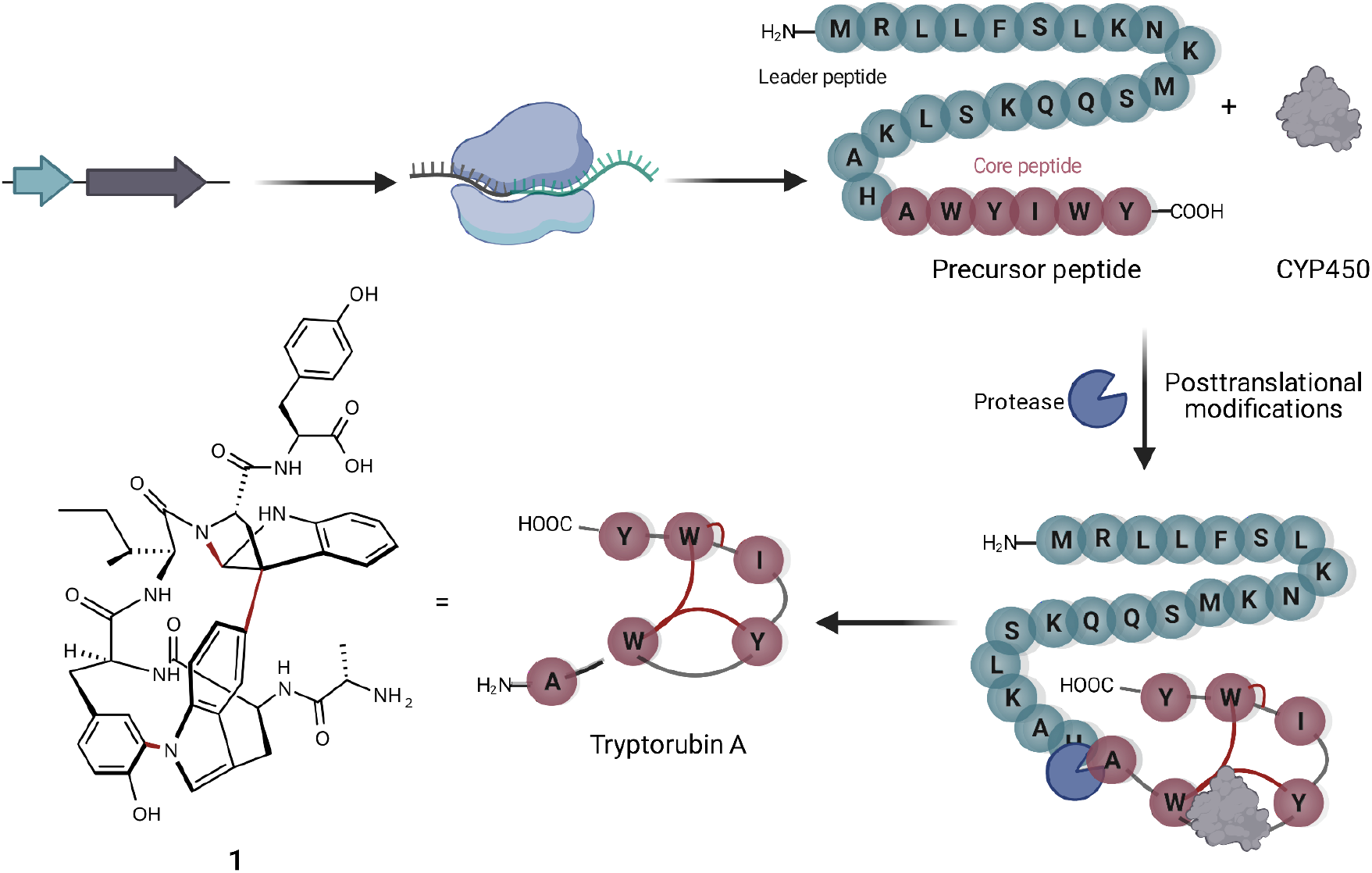
Schematic overview of the proposed model for tryptorubin A (1) biosynthesis. The genes of the *trp* BGC are transcribed and the resulting mRNA ribosomally translated. Subsequently, the precursor peptide, consisting of a core peptide and a leader peptide, is posttranslationally modified by the cytochrome P450 installing the characteristic atropopeptide crosslinks. Finally, an unspecific protease cleaves the leader peptide to release the hexapeptide tryptorubin A (**1**).^15^

In recent genome mining efforts, the atropopeptide-modifying P450 encoded in the *trp* BGC was subjected to BLAST analysis to identify homologs likely involved in atropopeptide maturation. The associated precursor peptide sequence was subsequently discerned through manual analysis of the genomic vicinity of the putative atropopeptide modifying P450s. While this approach has extended the atropopeptide family to encompass the cihunamides^16^, kitasatides^17^ and amyxirubins^15^ (also referred to as clavorubins^18^), it is apparent that this BLAST-based approach has limitations in capturing the full biosynthetic diversity of the atropopeptides. The BLAST algorithm excels at detecting closely related homologs of a query sequence.^10^ However, to identify more distant relatives of tryptorubin A, a more flexible strategy allowing significant deviations from the original sequence is imperative. This flexibility might be achieved through the utilization of supervised machine learning (ML).^19^ Supervised learning entails training algorithms on a labeled dataset, which are subsequently utilized for classifying previously unseen data by the trained algorithm.^20^ Herein we comprehensively explore the biosynthetic potential encoded in atropopeptide BGCs. We introduce AtropoFinder, a machine learning-based genome mining tool for the identification of atropopeptide BGCs. The AtropoFinder algorithm identified 683 putative atropopeptide BGCs across publically accessible genome sequences, thus significantly augmenting the number of atropopeptide BGCs compared to BLAST analysis. The identified atropopeptide BGCs revealed unexpected trends in core peptide composition and length, enabling us to refine the definition of atropopeptides and expand their biosynthetic and structural space. Insights from the identified putative atropopeptide precursor peptides along with modeling studies suggest that atropopeptide precursors share a single conserved residue in the core peptide and are characterized by a conserved motif in the leader sequences, likely facilitating precursor binding and anchoring the core peptide within the P450’s active site. In addition, co-occurrence analysis revealed trends in associated tailoring enzymes encoded within these BGCs. The expansion of the atropopeptide biosynthetic space is furthermore showcased by the AtropoFinder-guided identification of four new atropopeptides featuring different core peptide length, number and types of modifications. Moreover, we report the identification of the most complex atropopeptide reported to date and the unexpected production of two atropopeptides with the same core peptide sequence but distinct non-overlapping modification patterns from a single atropopeptide BGC.

## Results

### P450-guided identification of putative atropopeptide gene clusters (AtropoFinder)

Recent studies conducted by our research group and other laboratories describe the BLAST-based identification of atropopeptide biosynthetic pathways using the atropopeptide family-defining P450 as a query sequence.^15,16^ Manual analysis of the obtained results indicated that the identified core peptide sequences in the vicinity of the identified P450 genes exhibit a moderate level of sequence diversity.^15^ This lack of sequence diversity, suggests that the BLAST approach might not comprehensively capture the entire biosynthetic diversity of atropopeptides. Furthermore, the workflow is time-intensive and requires manual search to locate the associated precursor peptide gene in the proximity of the P450 gene. The challenge of identifying the precursor sequence is exacerbated with precursor peptide sequences that display only limited similarity to known atropopeptide precursors. To address these limitations, we hypothesized that a machine learning-based approach is more suitable to comprehensively map the atropopeptide biosynthetic landscape. In addition, our approach aims to replace the laborious manual precursor peptide identification with a robust algorithm that detects the precursor and core peptides regardless of their level of homology to characterized atropopeptides. Similar to the existing BLAST-based approaches, we decided to use the atropopeptide-defining P450s as bait for the identification of atropopeptide BGCs. After the initial identification, a subsequent step, focused on identifying the ORF coding for the precursor peptide, serves to confirm the identified BGC (CoreFinder). This step also allows for the detailed identification of the primary amino acid sequences of both precursor and core peptides.

To obtain a training data set for the ML classifier, a BLAST search was conducted using the P450 encoded in the *trp* BGC as a query sequence (WP_007820080.1). Through manual curation, we identified 51 sequences putatively encoding tryptorubin A-like precursors. After dereplication with a 95% sequence similarity cutoff, 37 sequences remained. To train the ML-classifier to differentiate atropopeptide-modifying P450s from P450s that modify other substrates, we collected a dataset from the antiSMASH database. This dataset consisted of P450s encoded in BGCs involved in the biosynthesis of products that are unrelated to atropopeptides that were dereplicated at a 95% cutoff, totaling 9,065 entries.^21^ We simplified the amino acid code based on the amino acids’ physico-chemical properties to prevent overfitting (Figures S2-S3). Next, we trained a machine-learning Random Forest classifier on a subset of the training data, which we validated using the remainder of the training dataset. We then applied the classifier to a dataset of 154,364 protein sequences from NCBI that were annotated as “P450” (for a detailed workflow and metrics see SI Detailed description of AtropoFinder, Figure S2-S4, Table S1). In the initial round of P450 classification, the algorithm identified 202 putative atropopeptide-modifying P450s. However, the associated precursors, predicted by CoreFinder (see below), only showed moderate core peptide sequence variations. To address these limitations, we iterated the process, incorporating the 202 newly identified atropopeptide-modifying P450s into the initial training dataset to refine the algorithm. Using the refined classifier, we identified 440 additional putative atropopeptide-modifying P450s from the dataset of 154,364 putative P450 sequences, thus more than doubling the number of putative atropopeptide-modifying P450s in this round.

### Identification of putative atropopeptide precursor- and core peptides (CoreFinder)

More than a dozen highly sophisticated RiPP genome mining tools for the identification of putative RiPP precursor peptides have been developed.^10^ While these tools have led to the identification of countless putative precursor sequences and the targeted discovery of numerous RiPPs, none of the existing tools is capable of identifying atropopeptide precursors and BGCs, respectively (SI Comparison of AtropoFinder to existing bioinformatic tools, Figure S5). The manual search process for atropopeptide precursor genes is not only laborious but may also miss out on sequences that deviate significantly from known atropopeptide precursors. To address this problem, we introduced "CoreFinder" (detailed description can be found in SI Detailed description of CoreFinder). The command-line tool streamlines the screening for open reading frames (ORFs) in the genomic vicinity of a specified protein-encoding gene (here specifically the gene encoding the characteristic atropopeptide-modifying P450) within all GenBank nucleotide records in which the query protein is annotated. We screened the genetic neighborhood of all putative atropopeptide-modifying P450 genes from NCBI records for atropopeptide-like open reading frames. The identification criteria prioritize precursor ORFs that range between 10 to 40 amino acids in length, consisting of a leader and a core peptide, which contains two consecutive aromatic amino acids and is followed by a stop codon. The latter criterion stemmed from the observation that the atropopeptides we had previously characterized all featured at least one pair of consecutive aromatic acids that are crucial for anchoring the characteristic atropopeptide modifications. While we experimented with detecting the presence of a ribosomal binding site (RBS) preceding the ORF of a potential precursor sequence, we noted that a significant number of manually detectable putative atropopeptide precursor ORFs do not seem to be associated with canonical RBSs. CoreFinder detected 803 potential precursor peptide-encoding genes. Out of these, 475 genes from 533 genomes exhibited core characteristics of the tryptorubin A precursor. Based on overall sequence homology, the rest seemed to be false positive hits.

### Analysis of atropopeptide precursor sequences and recalculation of putative atropopeptide precursor sequences

In our endeavor to characterize conserved motifs within atropopeptides, we aligned all identified putative precursor peptide sequences. We used the MEME suite to pinpoint the conserved KSLK motif (or closely related motifs like ’RSLK’, ’ESLK’, ’KSRK’, ’KPLK’, or ’PSLK’) strategically situated between residues 17-20 of the putative precursors (Figure 2; Figure S6). We thus hypothesized that the KSLK motif might be necessary for the recognition of the precursor by the P450. To verify our hypothesis we used AlphaFold2 multimer^22^ to investigate the predicted interaction between the precursor and corresponding P450 associated with tryptorubin A (Figure 2). With a iptm+ptm score of 0.8395, the model seems to be a good representation of the protein interaction (for details on the pLDDT score for each residue see Figure S7). Our analyses underscored the putatively critical role of the KSLK motif for threading the precursor into the P450’s substrate binding tunnel. The KSLK motif occupies a strategic junction, marking the transition from the N-terminal α-helix of the leader peptide —which is predicted to bind to the P450 enzyme’s surface—to a second, shorter α-helix which transitions into the seemingly flexible core peptide. Based on the AlfaFold model, the KSLK motif can be regarded as a biochemical "anchor", threading the precursor peptide into the substrate binding tunnel. This stabilization might allow the core peptide to “swirl” around within the enzyme’s active site. Consequently, this flexibility renders the formation of different bond types at different locations within the core peptide possible. Guided by the lessons learned from the combination of sequence analysis and modeling studies, we refined the classification algorithm of CoreFinder to mandate the inclusion of one of the following motifs within the precursor: ’KSLK’, ’RSLK’, ’ESLK’, ’KSRK’, ’KPLK’, or ’PSLK’. The recomputation unveiled 685 putative precursor peptides dispersed across 683 nucleotide sequences. Among these, 684 exhibited archetypal features of atropopeptide precursors. One sequence is thought to be a putative false positive. Although it possesses the "KSLK" motif, it significantly diverges from canonical leader peptide sequences and appeared to adopt this motif at an aberrant locus. Such deviation could represent a fusion of a atropopeptide BGC with another RiPP family. Two putative atropopeptide BGCs harbor two instead of one putative precursor gene (Figure 3). A BiG-SCAPE^23^ analysis comparing all obtained gene clusters with BGCs from MIBiG^23^ shows that atropopeptide BGCs cluster separately from all BGCs deposited to MIBiG^24^ (Figure S8). In a sequence similarity network of all identified putative precursor peptides (Figure S9), the precursor sequences group based on their core peptide sequence and phylogenetic origin. Surprisingly, the sequences associated with the tryptorubin A core peptide group into multiple clusters, suggesting that they might have evolved towards this sequence multiple times. A similar evolutionary pattern is observed in the phylogenesis of atropopeptide-modifying cytochrome P450s (Figure 3).

**Figure 2:**
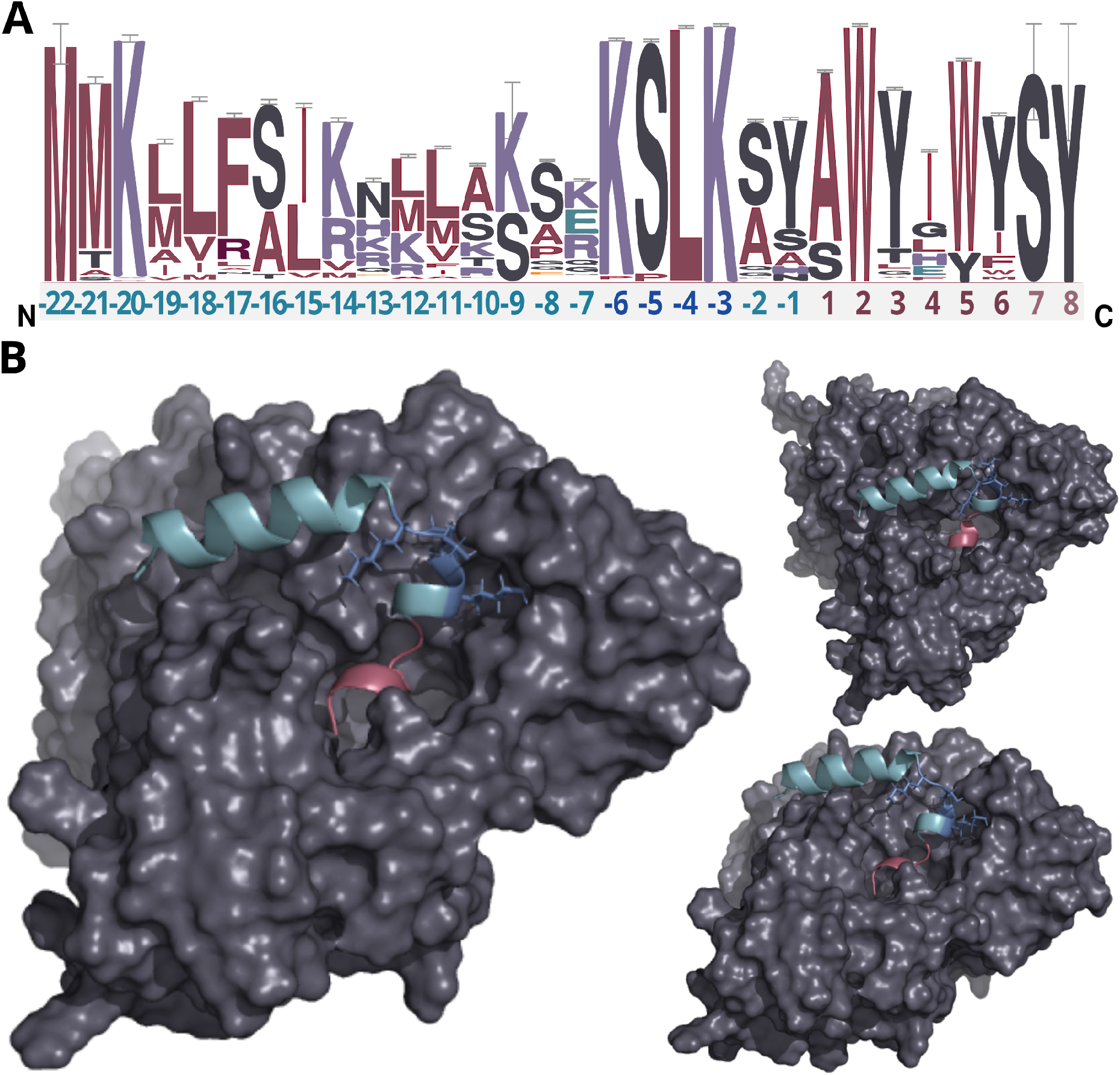
Analysis of putative precursor peptides and modeling of protein interactions of the precursor peptide and P450 encoded in the *trp* BGC. A) Sequence logo of putative precursor peptide sequences. The amino acids of the leader peptide, KSLK motif and core peptide are depicted in green, blue, and red, respectively. Residues only present in very few peptides are depicted in light red (residue 7-8). The height of each letter in the sequence logo corresponds to the frequency of the respective amino acid at that position, with taller letters indicating higher frequency and conservation. B) AlphaFold2 multimer protein model of WP_007820080.1 and the tryptorubin A precursor peptide from different angles (a video showcasing intricate details of the protein model can be found in Supplementary Data File S1, metrics of the modeling can be found in Figure S7). The leader peptide, KSLK motif, core peptide, and cytochrome P450 are depicted in green, blue, red, and gray, respectively. The leader peptide forms an α-helix that bends at the KSLK motif. The KSLK motif acts as an anchor of the precursor to the P450 that allows the core peptide to move freely within the active site.

**Fig 3:**
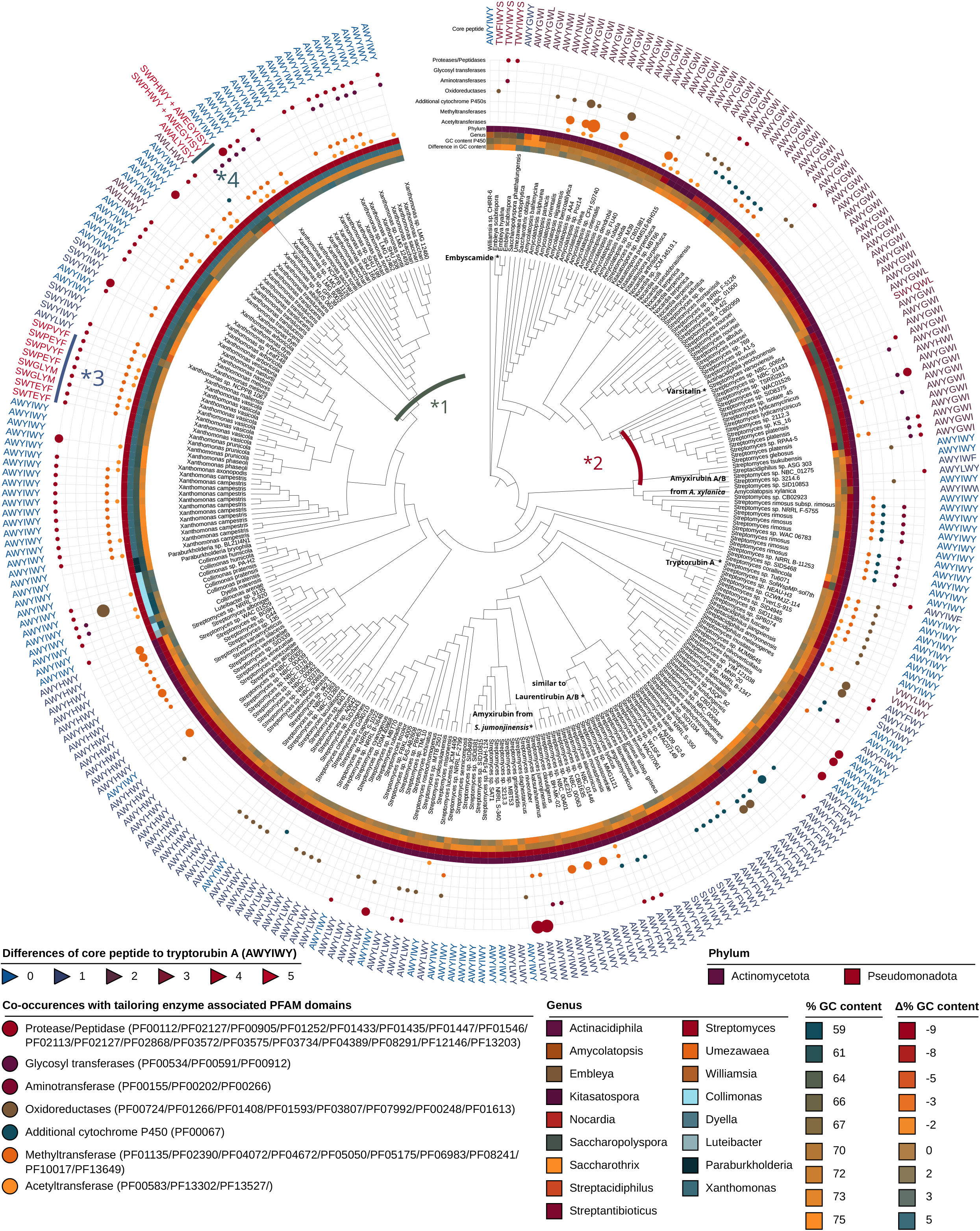
Phylogenetic tree of characteristic atropopeptide-modifying P450s encoded in putative atropopeptide BGCs, putative core peptide sequences as determined by CoreFinder and tailoring enzymes encoded in the associated BGCs. Depicted is the species, genus and phylum of the producer, their co-occurrence with tailoring enzyme-associated PFAM domains, the associated putative core peptide and its similarity to the tryptorubin A sequence. Additionally, the GC content of the P450 (absolute and relative to the GC content of the producer) is shown. The clade marked with *1 contains clusters with significantly higher GC content than the producer genome whereas the clade marked with *2 contains clusters with a significantly lower GC content than the genome of the producer, indicating relatively recent horizontal gene transfer events between the two producer phyla. The clade marked with *3 is associated with core peptides not containing two tryptophans and the clade marked with *4 is associated with two precursor peptides for each BGC.

### Analysis of atropopeptide core sequences

The analysis of all identified putative core peptides revealed that the majority are likely hexapeptides. Nonetheless, there are deviations from this norm. 14 are discerned as octapeptides, while three are classified as heptapeptides. Intriguingly, the octapeptides appear to incorporate additional amino acids interspersed amidst the known modified amino acids (W2, Y3, W6, Y7). In contrast, the heptapeptides exhibit an appended serine at their C-terminus (Figure 3). Amidst the core peptide sequences (Figure 2.A), the only conserved amino acid residue is W2. We hypothesize that the W2 plays a key role in one of two ways: it is either vital for forming the atropopeptide’s 3D shape, especially considering all characterized atropopeptides have at least one if not multiple crosslinks anchored to this tryptophan residue. Alternatively, W2 might be crucial for binding to the P450, although our modeling studies (Figure 2.B) do not corroborate this hypothesis. A striking observation is that not all putative core peptides encompass two successive aromatic amino acids, potentially implying varied modification motifs. Moreover, in comparison to a recent report^16^, not all core peptides exhibit a pair of tryptophan residues. Several core peptides originating from *Xanthomonas* species display an xWxxYx pattern, deviating from the conventional xWxxWx pattern reported before. This observation suggests that the recently coined bitryptides (xWxxW) refer to a subfamily of atropopeptides. Moreover, our in-silico studies indicate that atropopeptides can be more adequately classified by the presence of conserved motifs and residue in the precursor peptide than by their complex 3-dimensional shape.^16^

### Phylogenetic distribution of atropopeptides

The phylogenetic distribution of putative atropopeptide BGCs revealed their abundance mainly in *Streptomyces*, *Xanthomonas,* and related genera in the phyla actinomycetota and pseudomonadota (Figure 3, Figure S10). This abundance is surprising because these phyla are not closely related within the bacterial branch of life. To delve deeper into the evolutionary history of P450s encoded in putative atropopeptide BGCs, we constructed a maximum likelihood phylogenetic tree utilizing all putative atropopeptide-modifying P450 amino acid sequences (Figure 3). Notably, the sequences group based on the phylogeny of their producer organisms and the associated putative core peptide sequences.

Upon contrasting the GC content of the P450s (serving as a proxy for the GC content of the BGC) with the mean GC content of each species’ genome, we found a close correlation between the two for most BGCs. The observed pattern suggests that these BGCs have an ancient evolutionary origin and have adapted to the GC content of the producer’s genomes. However, two distinct clades of P450s presented significant disparities in their GC content relative to their producer’s genomes GC content. Within the clade designated as *1, the P450’s GC content exceeds that of the associated Pseudomonadota genomes, mirroring instead the GC profile characteristic for Actinomycetota. In contrast, the clade labeled as *2 reveals a GC content of the P450 that is markedly diminished relative to its corresponding Actinomycetota producer strains, but aligns more closely with GC contents characteristic for Pseudomonadota. Such observations might allude to recent horizontal gene transfer events between the two phyla.

### Analysis of the genomic vicinity of minimal atropopeptide BGCs

For an in-depth analysis of the tailoring enzymes encoded in the putative atropopeptide BGCs, we employed RODEO’s functionality to track concomitant Pfam domains encoded by a specified gene.^25^ Our findings underscore a frequent association between proteases and atropopeptide P450s within the genus of *Xanthomonas*. This is particularly noteworthy, as previously characterized atropopeptides do not harbor proteases encoded in their associated BGC. This association potentially facilitates the cleavage in instances where an external protease is absent for leader peptide excision or the proteases/peptidases are responsible for a secondary cleavage of the core peptide, reminiscent of the model for tryptorubin B biosynthesis. Other tailoring enzymes that are frequently encoded in atropopeptide BGCs are acetyltransferases, methyltransferases, oxidoreductases, additional P450s, and glycosyltransferases. It is tempting to speculate that the biosynthetic space of atropopeptides includes molecules that are more extensively modified by the identified tailoring enzymes than the previously characterized atropopeptides.

### Atropofinder-based expansion of the atropopeptide chemical space

To validate the AtropoFinder algorithm and expand the biosynthetic and chemical space of characterized atropopeptides, we selected four putative atropopeptide BGCs for characterization through heterologous expression in *Streptomyces albus* J1074.

We first selected the atropopeptide BGC from *Streptomyces jumonjinensis* DSM 747 (further referred to as *jum*) which was predicted to harbor the same core peptide sequence as the one reported for amyxirubin A and B. Since the homology of the P450s encoded in the *jum* and *amyxirubin* BGCs was only 59% and the P450 phylogeny suggested a distant relationship to the P450s involved in tryptorubin and amyxirubin biosynthesis (Figure 3), we speculated that the resulting atropopeptide might show a different bridging pattern when compared to both reported atropopeptides. Heterologous expression of the *jum* BGC led to the isolation of the associated atropopeptide (**2**) from the culture supernatant (Figure S11). The comparison of ^1^H NMR chemical shifts and high-resolution electrospray ionization mass spectrometry (HR-ESI-MS) data of **2** and an authentic standard of amyxirubin B (Figures S12-S13), however, revealed that the structure of **2** is identical to amyxirubin B, suggesting that the 3-dimensional shape characteristic for the amyxirubins and tryptorubins is widely distributed among the atropopeptide RiPP family.

We next set out to characterize a putative atropopeptide BGC from *Streptomyces varsoviensis* DSM 40346 (further referred to as *sva*). The BGC is unconventional in that it harbors one (2nd generation CoreFinder) or two (1st generation CoreFinder) precursor genes, dependent on which rules were used for the identification of the precursor and a single P450 gene. The predicted core peptide sequence of the first putative precursor peptide SvaA1 is CYSWYI, while the predicted core peptide sequence of the second precursor peptide SvaA2 is SWYQWL (Figure 4). The *sva* BGC SvarCore1 containing both putative precursor genes *svaA1*, *svaA2* and the P450 gene *svaB* or the *sva* BGC SvarCore2 containing only the second precursor gene *svaA2* and P450 gene *svaB* were heterologously expressed. Extraction of HP20 resin supplemented to the culture supernatant and analysis of the crude extract by HR-ESI MS revealed the presence of two compounds (**3** and **4**) detected at *m/z* 791.3540 [M + H]^+^ (calcd. for *m/z* 791.3511, C_42_H_47_N_8_O_8_^+^, Δ 3.67 ppm) and *m/z* 793.3655 [M + H]^+^ (calcd. for *m/z* 793.3668, C_42_H_49_N_8_O_8_^+^, Δ 1.64 ppm) (Figures S14 and S16) from both cultures. The calculated formulae of **3** and **4** are in good agreement with molecular formulae of the predicted pentapeptide variant of the second core peptide (WYQWL) that had lost four and two protons, respectively. The proton loss indicated at least one or two modifications, respectively. To increase the yield for subsequent purification, we inserted a ribosomal binding site in front of the gene encoding the P450, which resulted in a 100-fold increase in yield of the putative pentapeptides (Figure S15). One of the associated pentapeptides (**3**), named varsitalin B1, was purified from a 16 L culture using a combination of open column chromatography and semi-preparative HPLC. Analysis of the ^1^H and ^13^C NMR spectra, with the aid of 2D NMR correlations (Figures S17-S22 and Table S2), confirmed the amino acid sequence WYQWL. Compound **3** is a pyrroloindoline-containing bicyclic macrolactam that features a C–C bond bridging two Trp residues, along with two C–N bonds between Trp-1 and Tyr-1, and between Trp-2 and the amide nitrogen. These linkages result in an overall bridging pattern as described for tryptorubin B and amyxirubin B, albeit with a different amino acid sequence. Moreover, NOESY correlations of H-11(Trp-2)/H-38(Trp-1) and H-9(Trp-2)/H-40,41(Trp-1) suggest that **3** features the “bridge-above” configuration of the bicyclic macrolactam ring as previously reported for all complex atropopeptides (Figure S23). Based on the recently proposed terminology for the conformational isomerism of bicyclic peptides,^14^ we assigned the configuration of the bridge-above macrolactam for **3** as *P*_ansa_. The detection of **3** and **4**, that are both associated with the second putative precursor peptide SvaA2 suggests that the first putative precursor peptide is not associated with atropopeptide biosynthesis. Modeling studies carried out with AlphaFold2 multimer support this hypothesis, as the precursor peptide SvaA1 adopts a different, rather unstructured fold that indicates a more loose superficial interaction with the P450 hat prevents the positioning of the core peptide in the active site of the P450 (Figure S24). The AlphaFold2 multimer model of SwaA2 and the P450, however, suggests that the interaction of both partners resembles the interaction between TrpA and TrpB.

**Figure 4.**
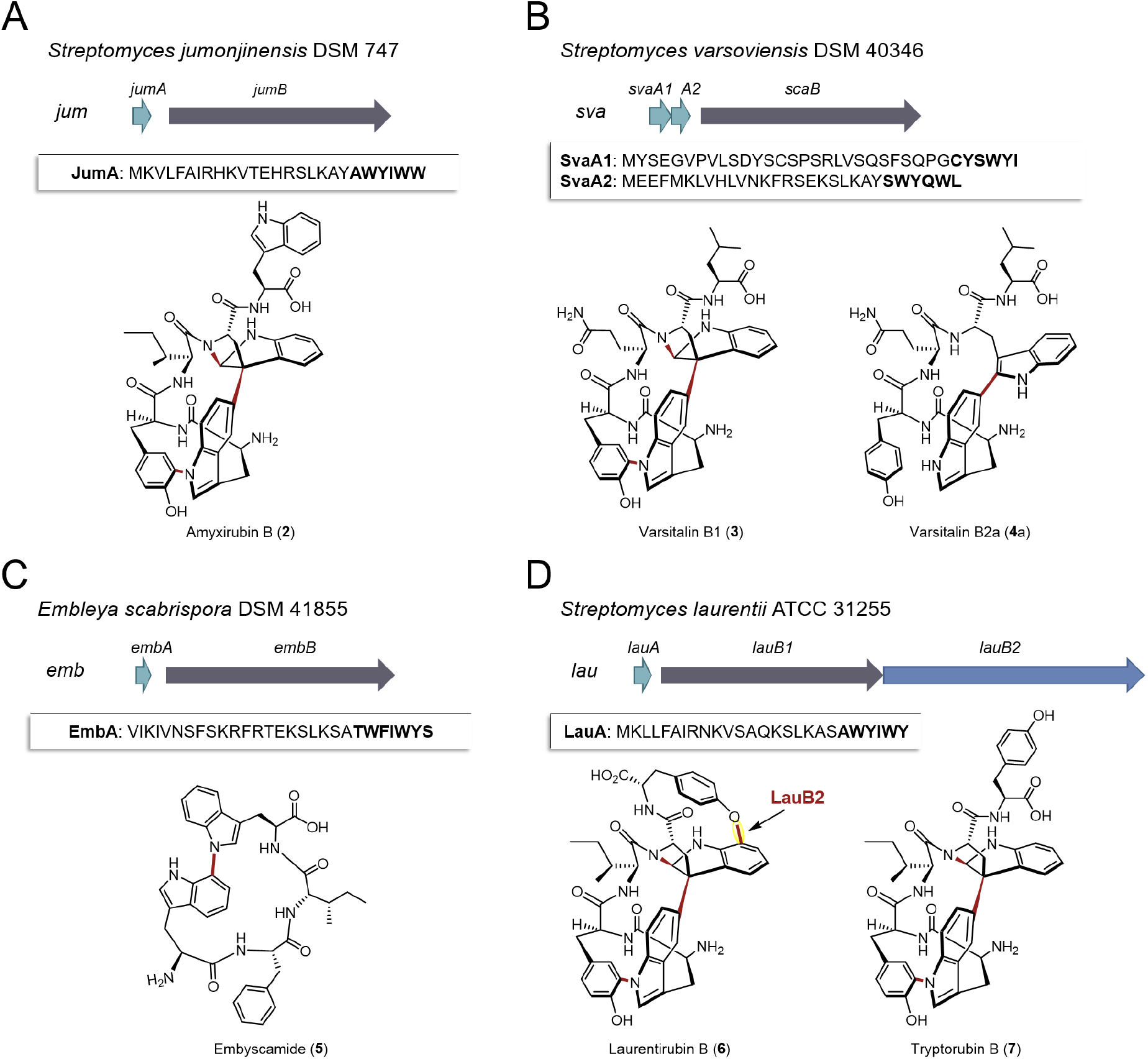
Characterization of four putative atropopeptide BGCs. The precursor genes and genes encoding the atropopeptide family-defining P450s are depicted in green and gray, respectively. The additional P450 is depicted in blue. The precursor peptide sequences are shown and the putative core peptides are highlighted in bold. A) *jum* BGC and structure of purified amyxirubin B (**2**) associated with the *jum BGC*. B) *sva* BGC and the structure of varsitalin B1 (**3**) and varsitalin B2a (**4a**) obtained by heterologous expression of the second precursor gene *svaA2* and the P450 encoding gene *svaB* in front of which a RBS was inserted. C) *emb* BGC and structure of purified embyscamide (**5**) obtained by heterologous expression of the *emb BGC*. D) *lau* BGC and structure of purified laurentirubin B (**6**) obtained from the heterologous expression of the full *lau* BGC and tryptorubin B (**7**), the product obtained from coexpressing *lauA* and *lauB1*. For a more detailed overview of the *lau* BGC, see Figure S56.

Careful analysis of HPLC-HR-ESI-MS data revealed that the second peak contained **4** and an additional compound with the same molecular formula as **4** (Figures S25-S26). Further chromatographic separation resulted in the isolation of **4a**, which we named varsitalin B2a, as well as its isomer **4b**. The structure of the isomer **4b** was not unambiguously elucidated by NMR analysis because of the low yield. Meanwhile, 1D and 2D NMR spectra of **4a** indicated that the molecule no longer possesses the complex three-dimensional shape that is characteristic for **3** (Figures S27-S34 and Table S3). Instead, only a single modification (C–C bond between C-39 of Trp-1 and C-11 of Trp-2) was present in **4a**. Interestingly, the position of this C–C bond at Trp-2 differs from the C–C bond in varsitalin B1 which is located at C-10 of Trp-2. **3** and **4a** share the same amino acid sequence but are characterized by a non-overlapping bridging pattern that is installed by the same P450.

Subsequently, we selected the putative atropopeptide BGC from *Embleya scabrispora* DSM 41855, (further referred to as *emb)* for characterization. The precursor EmbA harbors the putative heptapeptide core peptide sequence TWFIWYS (Figure 4). Culture extracts of *S. albus* harboring the *emb* BGC were screened for the presence of modified heptapeptides associated with the BGC. To our surprise, however, comparative HPLC-HR-ESI-MS analyses of extracts from *S. albus* cultures harboring the *emb* plasmid with an empty plasmid control revealed the presence of a new peak with an *m/z* 649.3132 [M + H]^+^ (calcd. for *m*/*z* 649.3133, C_37_H_41_N_6_O_5_ ^+^, Δ 0.15 ppm) (Figures S35-S36) that was significantly smaller than expected. The putative atropopeptide (**5**) was subsequently extracted and purified from a 10 L culture. Analysis of ^1^H and ^13^C NMR spectra, with the aid of HSQC, HMBC, and NOESY correlations (Figures S37-S43 and Table S4), revealed the presence of a tetrapeptide with the sequence WFIW featuring a C–N bond between C-34 of Trp-1 and N-5 of Trp-2 (Figure S44). We named the cyclic tetrapeptide embyscamide (**5**). The characterization of embyscamide with the sequence WFIW from the putative heptapeptide core TWFIWYS suggests that peptidases in the heterologous host cleave both the N-terminal and the C-terminal amino acids of the core peptide.

We next set out to characterize an atropopeptide BGC harboring an additional gene encoding another P450 to investigate possible additional tailoring reactions. AtropoFinder identified the putative BGC in *Streptomyces* sp. MMG1121, which was not available in any strain collection. We therefore conducted an NCBI BLAST^26^ homology search on the P450 to find a similar BGC in a commercially available strain. To our surprise, we identified a BGC from *Streptomyces laurentii* ATCC 31255 (further referred to as *lau* BGC) that harbors the characteristic atropopeptide-modifying P450 alongside three additional annotated P450 gene fragments that are significantly shorter than the first P450 encoded in the *lau* BGC (Figure S45). This BGC was not originally identified by AtropoFinder because we ran AtropoFinder solely on RefSeq P450s which excluded the *S. laurentii* P450. We initially hypothesized that the three P450 fragments might act together to form one functional P450. Alternatively, we hypothesized that the three P450 fragments are a sequencing artifact and that the three P450 gene fragments indeed encode one contiguous P450 such as in *Streptomyces* sp. MMG1121. To verify the latter hypothesis, we subjected the *lau* BGC to resequencing. Analysis of the sequencing data revealed that the three P450 gene fragments are indeed a sequencing artifact. As a consequence, the *lau* BGC is only predicted to harbor two genes encoding P450s, the atropopeptide-family defining P450 and a second P450 with low identity to TryB (28%) (Figure 4). To characterize the *lau* BGC, we cloned and heterologously expressed the BGC in *S. albus*.

HPLC-HR-ESIMS analysis of the crude extract obtained from the *S. albus* culture harboring the *lau* BGC revealed the presence of two candidate atropopeptides detected at *m/z* 895.3760 [M + H]^+^ (calcd. sum formula C_49_H_51_N_8_O_9_^+^, Δ 1.6 ppm) and 824.3401 [M + H]^+^ (calc. sum formula C_46_H_46_N_7_O_8_^+^, Δ 0.1 ppm) to be associated with the *lau* BGC (Figures S46-S47). The mass differences between the predicted core peptide sequence and the detected compounds are indicative of the candidate hexa- and pentapeptides with a loss of six protons which suggests the presence of at least three modifications. To pinpoint the localization of the modifications, the main product, a pentapeptide (laurentirubin B (**6**)) was isolated from a six liter culture. Based on the ^1^H NMR spectrum and 2D NMR correlations (Figures S48-S55 and Table S5), the structure of **6** was determined. **6** is characterized by a highly strained tricyclic macrolactam ring system that is constructed by two C–N bonds (Trp-1–Tyr-1 and Trp-2–peptide backbone), a C–C bond (Trp-1–Trp-2), and a C–O bond (Trp-2–Tyr-2) (Figure 4). The localization of the unprecedented aryl ether linkage between C-17 of Trp-2 and C-7 of Tyr-2 is supported by density functional theory (DFT) calculations (Table S6). Moreover, the atropisomeric configuration of *P*_ansa_ for **6** was determined by NOE correlations and ^13^C chemical shift calculation by the DFT method (Table S6). We hypothesized that the three common modifications are installed by the characteristic P450 homolog that is conserved in all atropopeptide BGCs and that the aryl ether bridge is formed via the second P450. To verify our hypothesis, we coexpressed both the precursor LauA with the first P450 LauB1 and LauA with the second P450 LauB2 in *S. albus* and analyzed the extracts for biosynthetic intermediates. The results showed that coexpressing *lauA* and *lauB2* resulted in the complete abolishment of atropopeptide production. While coexpressing *lauA* and *lauB1*, on the other hand, resulted in the accumulation of tryptorubin B (**7**) (Figure S56), a biosynthetic intermediate with three modifications, indicating that the second P450 is responsible for introducing the 4th modification. Laurentirubins are the most complex atropopeptides reported to date. Moreover, the *lau* BGC is the first characterized atropopeptide BGC harboring an additional gene that encodes a P450 that modifies the peptide beyond the characterized atropopeptide-specific modifications. Our gene coexpression studies furthermore indicate that the introduction of the ether bridge takes place after the formation of the archetypical atropopeptide modifications.

## Discussion

Atropopeptides are one of the recently described RiPP families that feature characteristic P450-catalyzed post-translational modifications that result in crosslinking of amino acid side chains (i.e. biaryl, aryl-nitrogen and aryl-ether linkages).^8,15,17^ Biarylitides are characterized by either a P450-catalyzed biaryl or a carbon-nitrogen bridge between histidine and tyrosine residues.^27^ Similarly, cittilins harbor both a biaryl and an aryl-ether bridge between tyrosine side chains, while atropopeptides are notable for featuring biaryl, carbon-nitrogen, and aryl-ether bridges, rendering them the most intricate RiPP family modified by P450s.^13,15^ Recently, a RiPP superfamily, named cyptides, that includes all RiPP families that feature biaryl linkages, has been proposed.^28^ Within this superfamily, the atropopeptides (also referred to as group 1 cyptides and atropitides) are the largest family with over 270 previously identified members.^28^ Our efforts to systematically map the biosynthetic space of the atropopeptides resulted in the significant expansion of the atropopeptide family over previous studies that employed BLAST-based approaches, which are inherently biased towards limited sequence diversity, and provided several key insights.

Our machine learning-based approach that uses the cytochrome P450 enzymes as a bait for the detection of atropopeptide BGCs, resulted in the identification of 684 putative atropopeptide BGCs. Insights from the comparison of all putative atropopeptide leader peptide sequences revealed a conserved KSLK motif in the leader that is characteristic for all atropopeptide leaders and that can be used to differentiate atropopeptides from other RiPP families that feature biaryl-linkages. Using AlphaFold2 multimer^22^ modeling, we were able to propose a putative role of the conserved motif in interacting with the P450, thus offering insights into its functional relevance as an anchor for the core peptide within the active site of the cytochrome P450. Moreover, all identified atropopeptide cores are characterized by the presence of a conserved Trp residue in the second position of the core peptide as the only conserved residue in the core. As a result, the recently proposed bitryptides^16^ that feature a xWxxWx core and the KSLK leader motif are a large subfamily of the atropopeptides. In contrast, atropopeptide precursors are characterized by the KSLK leader motif in combination with the xWxxxx core motif.

Our study and recent reports from other labs^16,17^ suggest that not all atropopeptides are characterized by the complex 3-dimensional shape recently reported for tryptorubin A which results in two possible non-canonical atropisomeric configurations. While most of the characterized atropopeptides (amyxirubin B, varsitalin B1 and laurentirubin B in this study) feature this unusual type of stereoisomerism, the family members that do not (embyscamide and varsitalin B2a in this study), at least, require canonical atropisomeric assignments.^16^ As a result, we propose to keep the name atropopeptides for the RiPP family. As the complex 3-dimensional shape is not a common characteristic of all atropopeptides, we propose to redefine the atropopeptide family based on their conserved biosynthetic features that include the presence of an atropopeptide-class defining P450, the conserved KSLK leader motif and a W residue at position two of the core peptide.

Validation of the AtropoFinder algorithm resulted in the characterization of four atropopeptide BGCs and the structure elucidation of their associated compounds. Embyscamide (**5**) features a single P450-catalyzed carbon-nitrogen bond between two tryptophan residues of the tetrapeptide core; the s*va* BGC produces varsitalins (**3** and **4a**) with three and one modifications, respectively, and laurentirubin B (**6**), the most complex atropopeptide reported to date, harbors a total of four modifications including an unprecedented ether bond between Trp-2 and Tyr-2. The presence of three different products with the same core peptide sequence and length (varsitalins) is remarkable as previous reports indicate the full conversion of the core peptide sequence into a single product. Moreover, the *lau* BGC is the only characterized atropopeptide BGC that does not harbor the minimal BGC architecture made up of precursor and P450 genes. The formation of the ether bond after the atropospecific formation of the bridge-above configuration of a tryptorubin-like intermediate by the second P450 encoded in the BGC significantly impacts the 3-dimensional shape of the molecule.

The three conserved bridges that are present in most characterized atropopeptides result in the formation of a highly strained bismacrocyclic ring that shows non-canonical atropisomerism. In accordance with the *P*_ansa_/*M*_ansa_ nomenclature for the stereochemical description of the conformationally diastereomeric bismacrocyclic peptides which was recently defined by Süssmuth and co-workers,^14^ we determined the configuration of the bicyclic macrolactam rings for all naturally occurring atropopeptides as *P*_ansa_. This nomenclature is applicable for the configurational description of atropopeptides in case they feature a bicyclic ring system. The only amino acid not involved in the formation of the bicyclic ring system in atropopeptides is the relatively flexible sixth amino acid of the core (e.g., Tyr6) which is not locked into place by the three characteristic modifications. In the case of laurentirubin B, the flexibility of Tyr6 gets significantly restrained by the fourth modification, making the 3D structure significantly more rigid.

The number of modifications installed into atropopeptides reported here ranges from one to four and the number of bond types from one (embyscamide), to two (e.g., varsitalins) and three (laurentirubin) that are installed by a single multi-functional P450 or alternatively four through the joint action of two P450s in the case of laurentirubin.

In conclusion, our research expands the atropopeptide family by more than 50%, provides a comprehensive understanding of atropopeptide biosynthesis and timing of maturation. The systematic investigation of atropopeptide biosynthetic space lays the foundation for future investigations into this peptide family and showcases the potential of machine learning-based genome mining algorithms for the identification of non-canonical biosynthetic pathways that elude unrecognized by existing genome mining tools.

## Supporting information

Video of the AlphaFold2 multimer protein model of WP_007820080.1

## Acknowledgements

EJNH gratefully acknowledges funding by the LOEWE Center for Translational Biodiversity Genomics (LOEWE TBG), the German Research Foundation (Emmy Noether Program), and the Funds of the Chemical Industry Germany. FB is supported by a Kekulé Fellowship of the Funds of the Chemical Industry Germany. We thank Bertholt Gust for providing strains and plasmids. We gratefully thank Gabriele Sentis for her technical assistance with NMR measurements. Open Access funding enabled and organized by Projekt DEAL. Figure 1 was created in BioRender.

## Data Availability Statement

All code and raw data pertaining to the machine learning analyses presented in this study can be accessed at the following GitHub repository: https://github.com/FriederikeBiermann/AtropoFinder. Furthermore, all biosynthetic gene clusters (BGCs) and associated compounds derived from this research have been submitted to the MiBIG (Minimum Information about a Biosynthetic Gene Cluster) database for public access and reference.

