## Supplementaries for "Machine learning-based exploration, expansion and definition of the atropopeptide family of ribosomally synthesized and posttranslationally modified peptides"

### Table of contents

| | | |
|---|---|---|
| **Detailed description of AtropoFinder** | | S5 |
| **Detailed description of CoreFinder** | | S5 |
| **Comparison of AtropoFinder to existing bioinformatic tools** | | S6 |
| **Figure S1** | Overview over characterized peptide modifications in non-ribosomal peptides and ribosomally synthesized and posttranslationally modified peptides. | S7 |
| **Figure S2** | Featurisation of the amino acid sequences of P450s to feed into the machine learning classifier. | S8 |
| **Figure S3** | Simplified amino acid code that reflects the physicochemical properties of the proteinogenic amino acids. | S9 |
| **Figure S4** | Precision-recall curve of atropopeptide-modifying P450s for conditional random forest classifier utilized for the prediction of atropopeptide-modifying P450s. | S10 |
| **Figure S5** | Comparison of BGC predictions of different state-of-the-art genome mining tools on a nucleotide sequence containing the atropopeptide BGCs identified by AtropoFinder. | S11 |
| **Figure S6** | Meme conserved motif search. | S12 |
| **Figure S7** | Metrics of the AlphaFold predicted model of precursor peptide and P450 encoded in the *trp* BGC. | S13 |

| | | |
|---|---|---|
| **Figure S8** | BiG-SCAPE analysis of all putative atropopeptide-containing gene clusters (red) and all characterized BGC deposited to the MiBIG database (blue) at threshold of 0.5. | S14 |
| **Figure S9** | Sequence similarity network of all putative atropopeptide precursors and their corresponding core peptide sequences. | S15 |
| **Figure S10** | Phylogenetic distribution of putative atropopeptide clusters. | S16 |
| **Figure S11** | Extracted ion chromatograms of amyxirubin B and jumorubin (**2**). | S16 |
| **Figure S12** | HPLC-ESI-QTOF-HRMS analysis of jumorubin (**2**). | S17 |
| **Figure S13** | $^{1}$H NMR spectrum (600 MHz) of jumorubin (**2**) in DMSO-$d_6$. | S17 |
| **Figure S14** | Extracted ion chromatogram of varsitalin B1 (**3**) and varsitalin B2 (**4**). | S18 |
| **Figure S15** | Extracted ion chromatogram of varsitalin B1 (**3**) and varsitalin B2 (**4**). | S19 |
| **Figure S16** | HPLC-ESI-QTOF-HRMS analysis of varsitalin B1 (**3**). | S20 |
| **Figure S17** | $^{1}$H NMR spectrum (600 MHz) of varsitalin B1 (**3**) in DMSO-$d_6$. | S20 |
| **Figure S18** | $^{13}$C NMR spectrum (150 MHz) of varsitalin B1 (**3**) in DMSO-$d_6$. | S21 |
| **Figure S19** | HSQC spectrum (600 MHz) of varsitalin B1 (**3**) in DMSO-$d_6$. | S21 |
| **Figure S20** | COSY spectrum (600 MHz) of varsitalin B1 (**3**) in DMSO-$d_6$. | S22 |
| **Figure S21** | HMBC spectrum (600 MHz) of varsitalin B1 (**3**) in DMSO-$d_6$. | S22 |
| **Figure S22** | NOESY spectrum (600 MHz) of varsitalin B1 (**3**) in DMSO-$d_6$. | S23 |
| **Figure S23** | Structure elucidation of varsitalin B1 (**3**). | S24 |
| **Figure S24** | AlphaFold2 model of SvaB P450 with two putative precursor peptides SvaA1 and SvaA2. | S25 |
| **Figure S25** | Extracted ion chromatograms of (A) varsitalin B2b (**4b**), (B) varsitalin B2a (**4a**) and (C) varsitalin B1 (**3**). | S25 |
| **Figure S26** | HPLC-ESI-QTOF-HRMS analysis of varsitalin B1 (**3**). | S26 |
| **Figure S27** | $^{1}$H NMR spectrum (600 MHz) of varsitalin B2a (**4a**) in DMSO-$d_6$. | S26 |
| **Figure S28** | HSQC spectrum (600 MHz) of varsitalin B2a (**4a**) in DMSO-$d_6$. | S27 |
| **Figure S29** | COSY spectrum (600 MHz) of varsitalin B2a (**4a**) in DMSO-$d_6$. | S27 |

| | | |
|---|---|---|
| **Figure S30** | Structure elucidation of varsitalin B2a (**4a**). | S28 |
| **Figure S31** | HMBC spectrum (600 MHz) of varsitalin B2a (**4a**) in DMSO-*d*$_{6.}$ | S29 |
| **Figure S32** | TOCSY spectrum (600 MHz) of varsitalin B2a (**4a**) in DMSO-*d*$_6$. | S29 |
| **Figure S33** | NOESY spectrum (600 MHz) of varsitalin B2a (**4a**) in DMSO-*d*$_6$. | S30 |
| **Figure S34** | Key NOESY correlations of varsitalin B2a (**4a**) in DMSO-*d*$_6$. | S30 |
| **Figure S35** | Extracted ion chromatogram of embyscamide (**5**). | S31 |
| **Figure S36** | HPLC-ESI-QTOF-HRMS analysis of embyscamide (**5**). | S32 |
| **Figure S37** | $^1$H-NMR spectrum (500 MHz) of embyscamide (**5**) in DMSO-*d*$_6$. | S33 |
| **Figure S38** | $^{13}$C-NMR spectrum (125 MHz) of embyscamide (**5**) in DMSO-*d*$_6$. | S34 |
| **Figure S39** | DEPT135 spectrum (125 MHz) of embyscamide (**5**) in DMSO-*d*$_6$. | S35 |
| **Figure S40** | HSQC spectrum (500 MHz) of embyscamide (**5**) in DMSO-*d*$_6$. | S36 |
| **Figure S41** | COSY spectrum (500 MHz) of embyscamide (**5**) in DMSO-*d*$_6$. | S37 |
| **Figure S42** | HMBC spectrum (500 MHz) of embyscamide (**5**) in DMSO-*d*$_6$. | S38 |
| **Figure S43** | NOESY spectrum (500 MHz) of embyscamide (**5**) in DMSO-*d*$_6$. | S39 |
| **Figure S44** | Structure elucidation of embyscamide (**5**). | S40 |
| **Figure S45** | Resequencing of *lau* BGC. | S41 |
| **Figure S46** | Extracted ion chromatogram of pentapeptide laurentirubin B (**6**) and hexapeptide | S41 |
| **Figure S47** | HPLC-ESI-QTOF-HRMS analysis of laurentirubin B (**6**). | S42 |
| **Figure S48** | $^1$H NMR spectrum(600 MHz) of laurentirubin B (**6**) in DMSO-*d*$_6$. | S42 |
| **Figure S49** | HSQC spectrum (600 MHz) of laurentirubin B (**6**) in DMSO-*d*$_6$. | S43 |
| **Figure S50** | COSY spectrum (600 MHz) of laurentirubin B (**6**) in DMSO-*d*$_6$. | S43 |
| **Figure S51** | TOCSY spectrum (600 MHz) of laurentirubin B (**6**) in DMSO-*d*$_6$. | S44 |
| **Figure S52** | Structure of laurentirubin B (**6**). | S44 |
| **Figure S53** | HMBC spectrum (600 MHz) of laurentirubin B (**6**) in DMSO-*d*$_6$. | S45 |
| **Figure S54** | NOESY spectrum (600 MHz) of laurentirubin B (**6**) in DMSO-*d*$_6$. | S45 |

| | | |
|---|---|---|
| **Figure S55** | Key NOESY of laurentirubin B (**6**) in DMSO-$d_6$. | S46 |
| **Figure S56** | *In vivo* functional characterization of P450s in *lau* BGC. | S46 |
| **Figure S57** | The lowest energy conformer of $P_{ansa}$-**6a**, $M_{ansa}$-**6**, and $P_{ansa}$-**6** calculated at B3LYP/6-31G(d,p) level of theory. The values show distance (Å) of protons which showed NOESY correlations in **6**. Protons not involved in one of the NOESY correlations are hidden. | S47 |
| **Table S1** | Detailed metrics of final classifiers for differentiating atropopeptide-modifying P450s (class 1) from P450s modifying any other substrate (class 0). | S48 |
| **Table S2** | $^{1}$H and $^{13}$C NMR Data for varsitalin B1 (**2**) in DMSO-$d_6$. | S49 |
| **Table S3** | $^{1}$H and $^{13}$C NMR Data for varsitalin B2a (**4a**) in DMSO-$d_6$. | S50 |
| **Table S4** | $^{1}$H and $^{13}$C NMR Data for embyscamide (**5**) in DMSO-$d_6$. | S51 |
| **Table S5** | $^{1}$H and $^{13}$C NMR Data for laurentirubin (**6**) in DMSO-$d_6$. | S52 |
| **Table S6** | Comparison of experimental and calculated $^{13}$C NMR chemical shifts for **6** | S53 |
| **Table S7** | Strains and plasmids used in this study | S54 |
| **Table S8** | Primers used in this study | S55 |
| **Supplementary methods** | | S56 |
| **References** | | S70 |

| | | |
|---|---|---|
| **Supplementary data file S1** | Video of the AlphaFold2 multimer protein model of WP_007820080.1 and the tryptorubin A precursor peptide visualized from different angles. The leader peptide, KSLK motif, core peptide, and cytochrome P450 are depicted in green, blue, red, and gray, respectively. | P450_precursor_movie.mp4 |

### Supplementary Figures and tables

#### Detailed description of AtropoFinder

**AtropoFinder** is a Jupyter notebook-based script designed to handle the data preprocessing and machine learning to identify P450s involved in atropopeptide modification.

The primary input for AtropoFinder consists of amino acid sequence alignments of query P450s in fasta format. The first objective is to featurize these sequences to transform the raw amino acid sequences into numerical features that can be inputted into the machine learning algorithm (Figure S2). To map the specific features to functional regions of the P450, the alignments were prepared by aligning the query P450 sequences against a reference P450 for which functional regions have been annotated. The alignments are separated into functionally annotated regions based on insights from the annotations of the reference sequence. This method results in fragments of each P450 sequence associated with a specific functional region of the P450 (Methods, Data preprocessing). To prevent overfitting and focus less on sequence homology and more on binding properties of the amino acids, sequences are transformed into a simplified amino acid code that groups amino acids with similar physio-chemical properties under one character (Figure S3, Methods, Data preprocessing).

The next phase is the featurization, wherein overlapping occurrences of k-mer motifs with a length of four are counted within the fragmented sequences. To reduce the number of features and target atropopeptide-specific features, only k-mer motifs are considered that occur in at least half of the P450s of the atropopeptide training dataset (Figure S2 and Methods, Training of the classifier and hyperparameter optimization).

The machine learning algorithm utilizes a Random Forest classifier trained in advance on the atropopeptide training data set (Methods, Training data set assembly and Training of the classifier and hyperparameter optimization) to classify the sequences based on the features calculated in the step before. This classifier was selected based on its performance metrics, particularly its f1 score for atropopeptide P450s and is implemented in scikit-learn (Methods, Training of the classifier and hyperparameter optimization).

Atropofinder outputs a CSV table of all results, as well as a fasta file containing all P450s predicted to be atropopeptide-modifying enzymes.

### Detailed description of CoreFinder

**CoreFinder** functions as a command-line tool designed to process fasta files of peptides, utilizing their accession numbers in the header for identification. CoreFinder's main functionality is to search for occurrences of genes coding these non-redundant proteins in the NCBI database using NCBI Entrez and subsequently analyze the genomic regions around these genes.

During its operation, CoreFinder identifies potential genes encoding precursor and core peptides. Open reading frames (ORFs) ranging between 10 to 40 amino acids that possess 'KSLK', 'RSLK', 'ESLK', 'KSRK', 'KPLK', or 'PSLK' in the precursor peptide are flagged. These ORFs should not coincide with existing coding sequences (CDS), with the exception of those categorized as "tryptorubin family RiPP precursor CDS". Overlapping precursor peptides ORFs are filtered to remove duplicates, keeping the shortest possible ORF. It is observed that the annotation of the exact start, and consequently, the length of precursor peptides encoding ORF can be difficult if multiple putative start codons are present. The output of the tool is a genbank file, capturing the 3kb region both upstream and downstream of the gene encoding the P450 of interest including the annotated precursor peptides. A more detailed description of the exact criteria can be found in the Methods Corefinder.

To aid its functionality, CoreFinder has several command-line options available to the user, including specifying input files, setting a genomic boundary for search, enabling dynamic core detection (instead of using the last 6 amino acids, it automatically uses the amino acid in front of W2 as the start of the core protein), and setting up an email used for the queries at NCBI Entrez. The tool also allows users to specify the desired output directory.

### Comparison of AtropoFinder to existing bioinformatic tools

To compare the AtropoFinder results with existing state-of-the-art genome mining tools, all putative AtropoFinder BGCs were combined into one fasta file and used as input for different state-of-the-art genome mining tools. Using the antiSMASH 7 beta webserver and the antismash 6.1.1 webserver, 9 BGC regions were detected, all annotated as "indole", found based on the indole_PTase. The atropopeptide cytochrome p450 gene was labeled as "biosynthetic additional", whereas the precursor gene was labeled as "other gene". Based on the close inspection of the antiSMASH results, it can be assumed that antiSMASH did not detect the atropopeptide cluster, but rather a closeby indole BGC and thus annotated the atropopeptide BGC by chance. DeepBGC, which uses a neural network to distinguish BGC regions, annotated 108 putative gene clusters, usually labeled as "Polyketide-Terpene". These putative BGCs are much larger than the BGCs annotated by AtropoFinder, often containing multiple BGCs predicted by AtropoFinder. The tool GECCO labeled 38 regions as putative gene clusters most of which were not attributed to any natural product class (with the exception of one labeled "terpene"). The results of this query are displayed in Figure S5. A query with PRISM 4 showed no results, and a rodeo search with the tryptorubin A cytochrome P450 gene as a query only resulted in one result, the protein WP_007820080 of *Streptomyces* sp. SID8380 with no annotated precursors. In conclusion, all identified entities were seemingly discovered by chance and none of them were identified as a RiPP BGC.

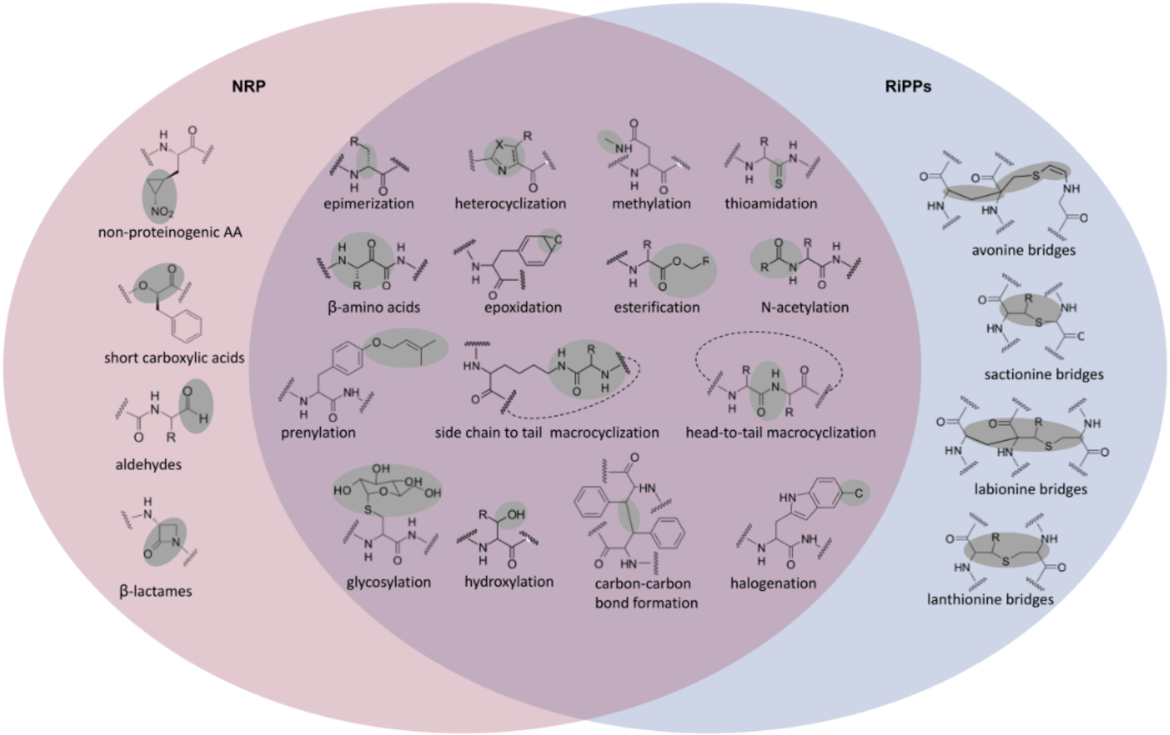


**Figure S1**. Overview over characterized peptide modifications in non-ribosomal peptides and ribosomally synthesized and posttranslationally modified peptides. The overview is intended to illustrate each type of modification with one example, rather than to catalog all characterized modifications. The overview does not aim to be exhaustive.

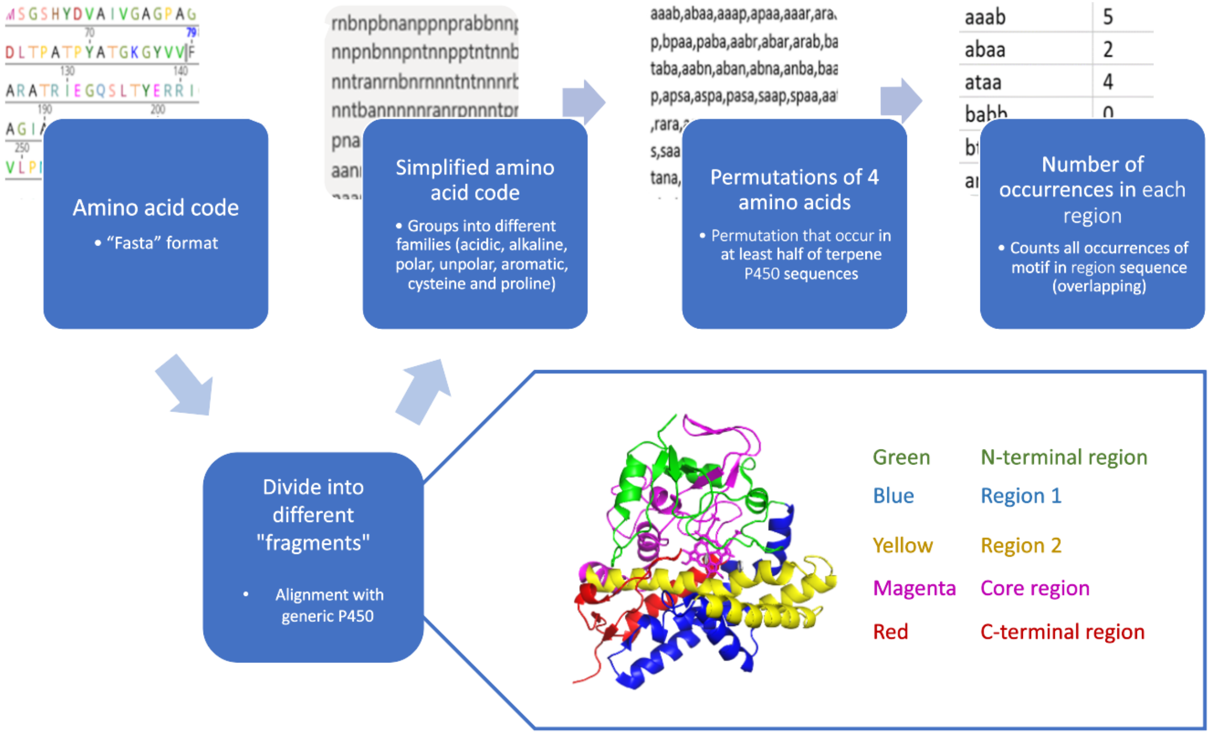


**Figure S2**. Featurisation of the amino acid sequences of P450s to feed into the machine learning classifier: First, the amino acid sequences are aligned against a reference P450 and split into fragments that resemble functional regions within the reference P450. Then, those fragmented amino acid sequences are translated into a simplified amino acid code (Figure S3). Subsequently, the number of occurrences of 4 amino acid long motifs is counted in each fragment and used as a feature.

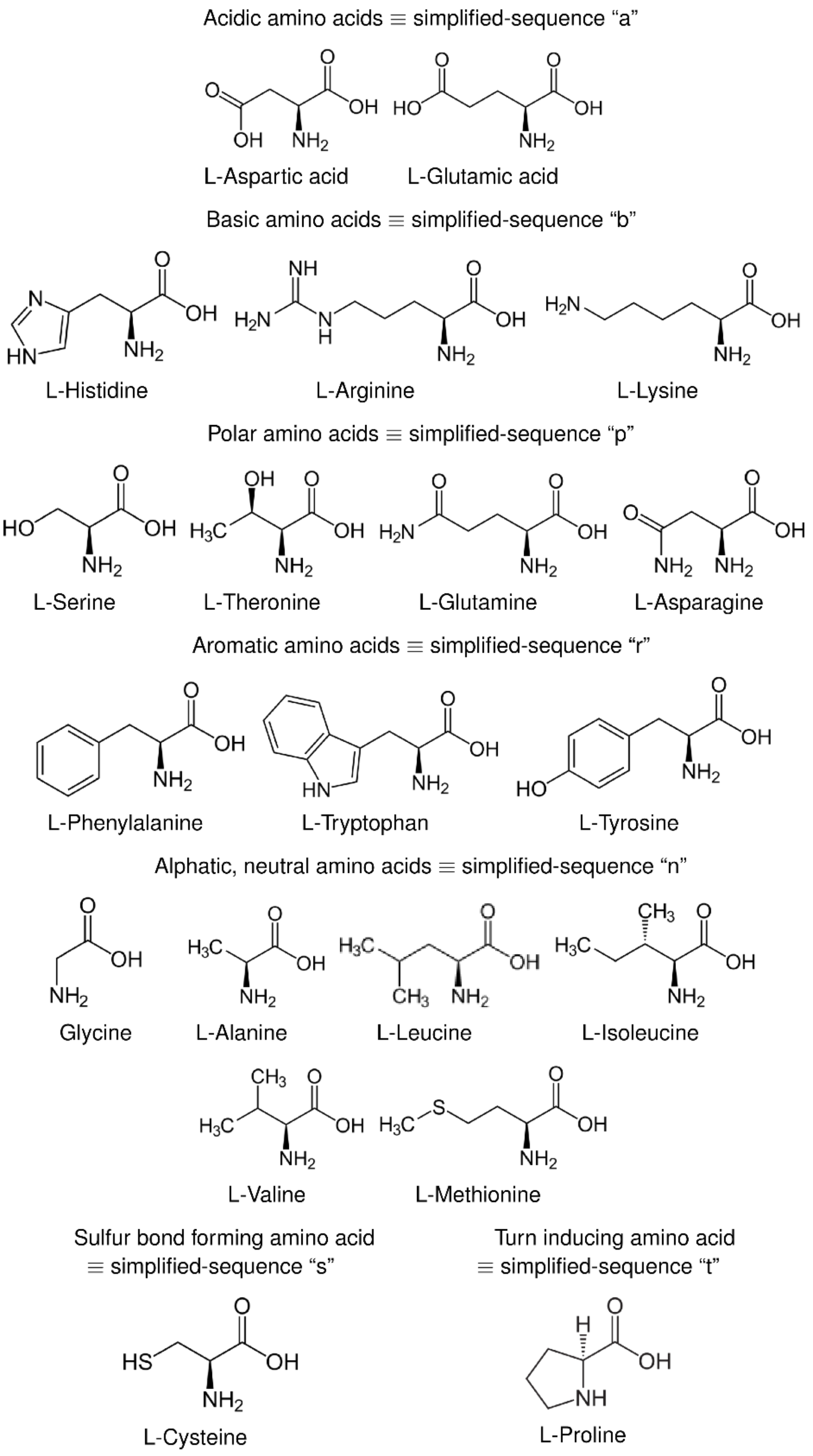


**Figure S3**. Simplified amino acid code that reflects the physicochemical properties of the proteinogenic amino acids.

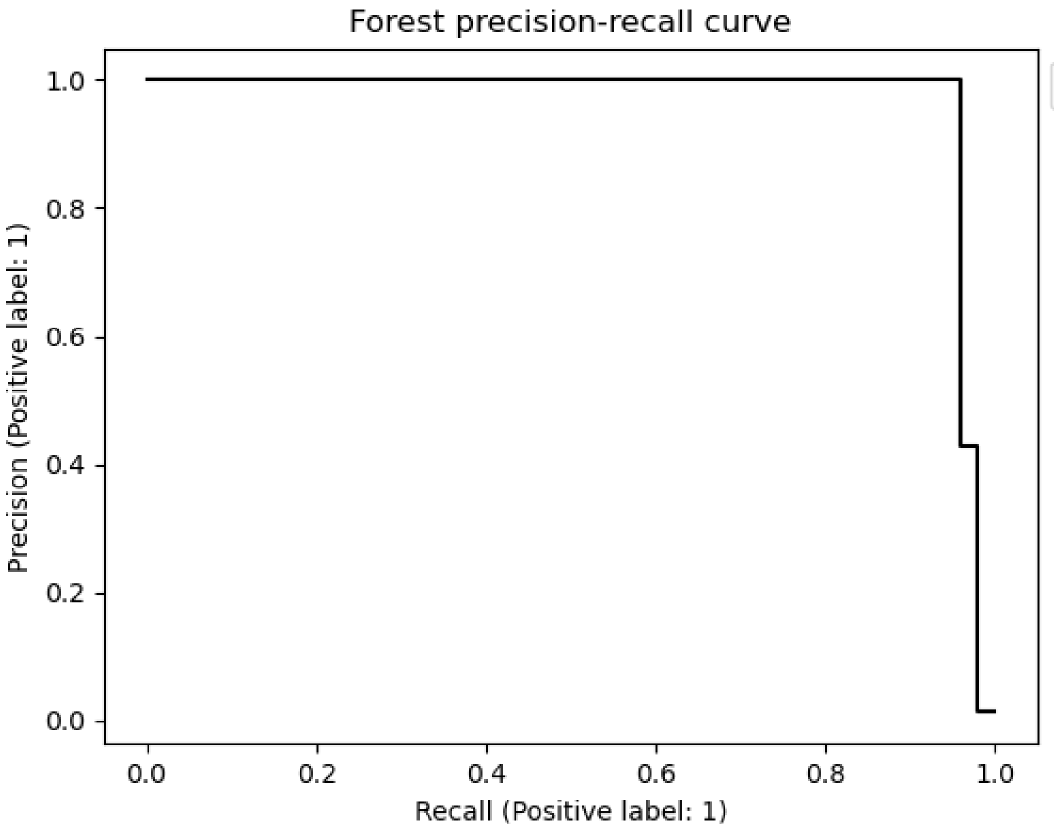


**Figure S4**. Precision-recall curve of atropopeptide-modifying P450s for conditional random forest classifier utilized for the prediction of atropopeptide-modifying P450s.

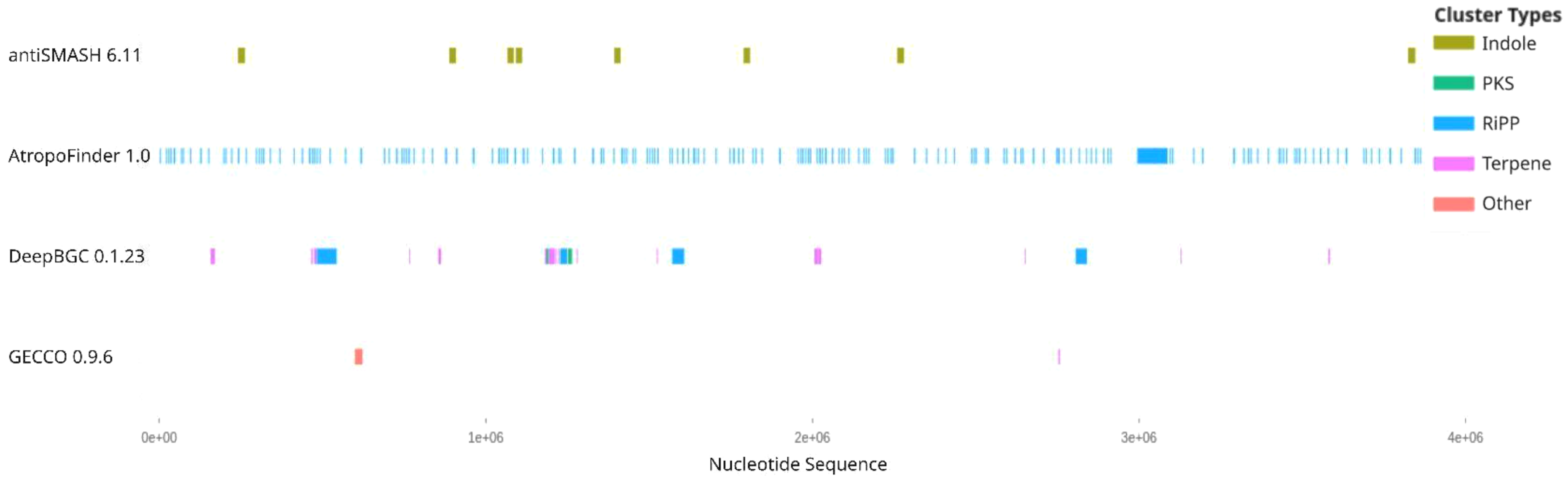


**Figure S5**. Comparison of BGC predictions of different state-of-the-art genome mining tools on a nucleotide sequence containing the atropopeptide BGCs identified by AtropoFinder.

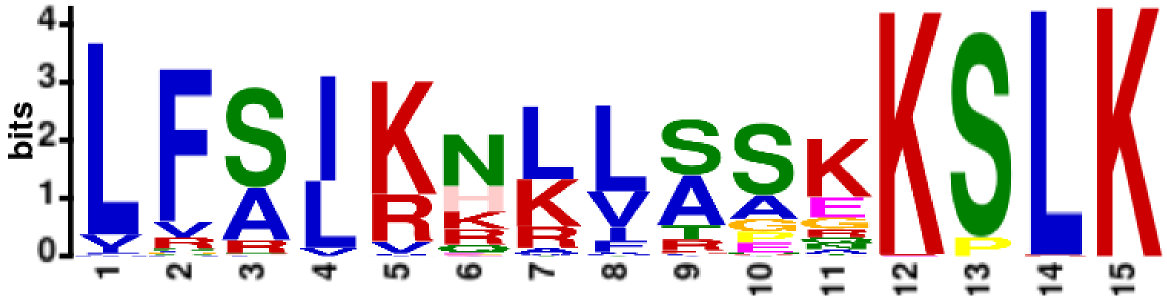

**Figure S6**. Meme conserved motif search. The sequence logo represents a significant motif (E-value: 1.0e-2100) observed 338 times, spanning a width of 15 amino acids. The motif predominantly features the conserved “KSLK” sequence at the C-terminus, preceded by 9 variable or unspecified residues, as well as a conserved “LF” motif at the N-terminus. The height of each letter in the sequence logo corresponds to the frequency of the respective amino acid at that position, with taller letters indicating higher frequency and conservation.

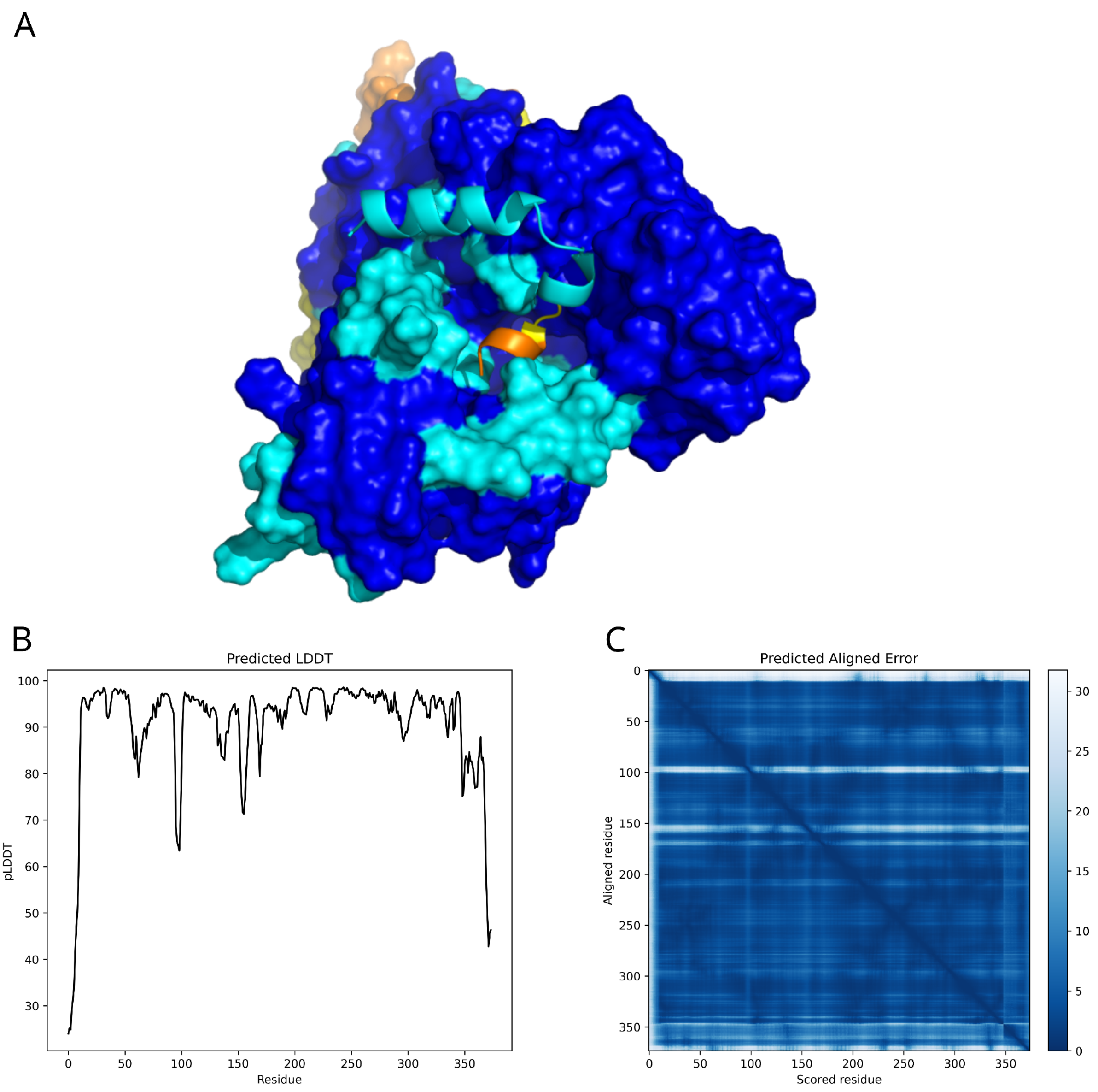


**Figure S7.** Metrics of the AlphaFold predicted model of precursor peptide and P450 encoded in the *trp* BGC. A) 3D structure representation: The model was predicted using AlphaFold. The structure is color-coded based on the predicted local distance difference test (pLDDT) scores. High-confidence regions with pLDDT scores above 90 are colored in blue, while regions with scores between 70 and 90 are colored in cyan. Moderate-confidence regions with scores between 50 and 70 are represented in yellow, and low-confidence regions with scores below 50 are displayed in orange. B) pLDDT Distribution: Line plot of predicted pLDDT scores against protein sequence, illustrating regions of modeling certainty and uncertainty. C) Predicted aligned error heatmap: A heatmap displaying the predicted aligned error for each pair of residues. The color intensity, ranging from dark (low error) to light blue (high error), signifies the magnitude of the aligned error, with the scale provided by a colorbar. This heatmap representation provides a comprehensive view of the alignment's reliability and potential areas where the predicted structure might deviate from a potential experimentally determined structure. In both B) and C), the P450 is represented by residues 1-449, while the precursor peptide is represented by residues 350-376.

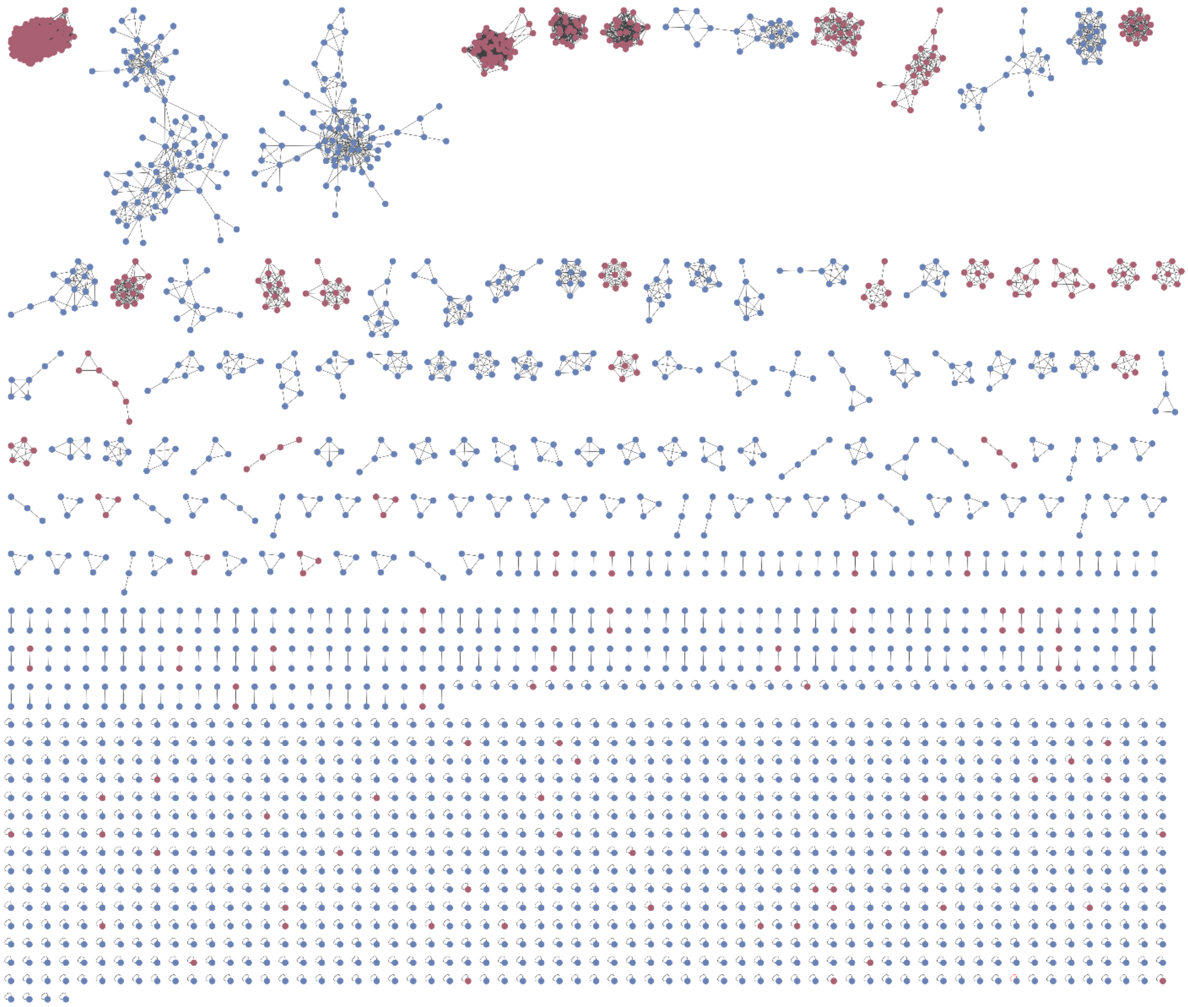

**Figure S8**. BiG-SCAPE analysis of all putative atropopeptide-containing gene clusters (red) and all characterized BGC deposited to the MiBIG database (blue) at threshold of 0.5. The analysis shows that atropopeptide BGCs cluster separately from all BGCs deposited to the MiBIG database.

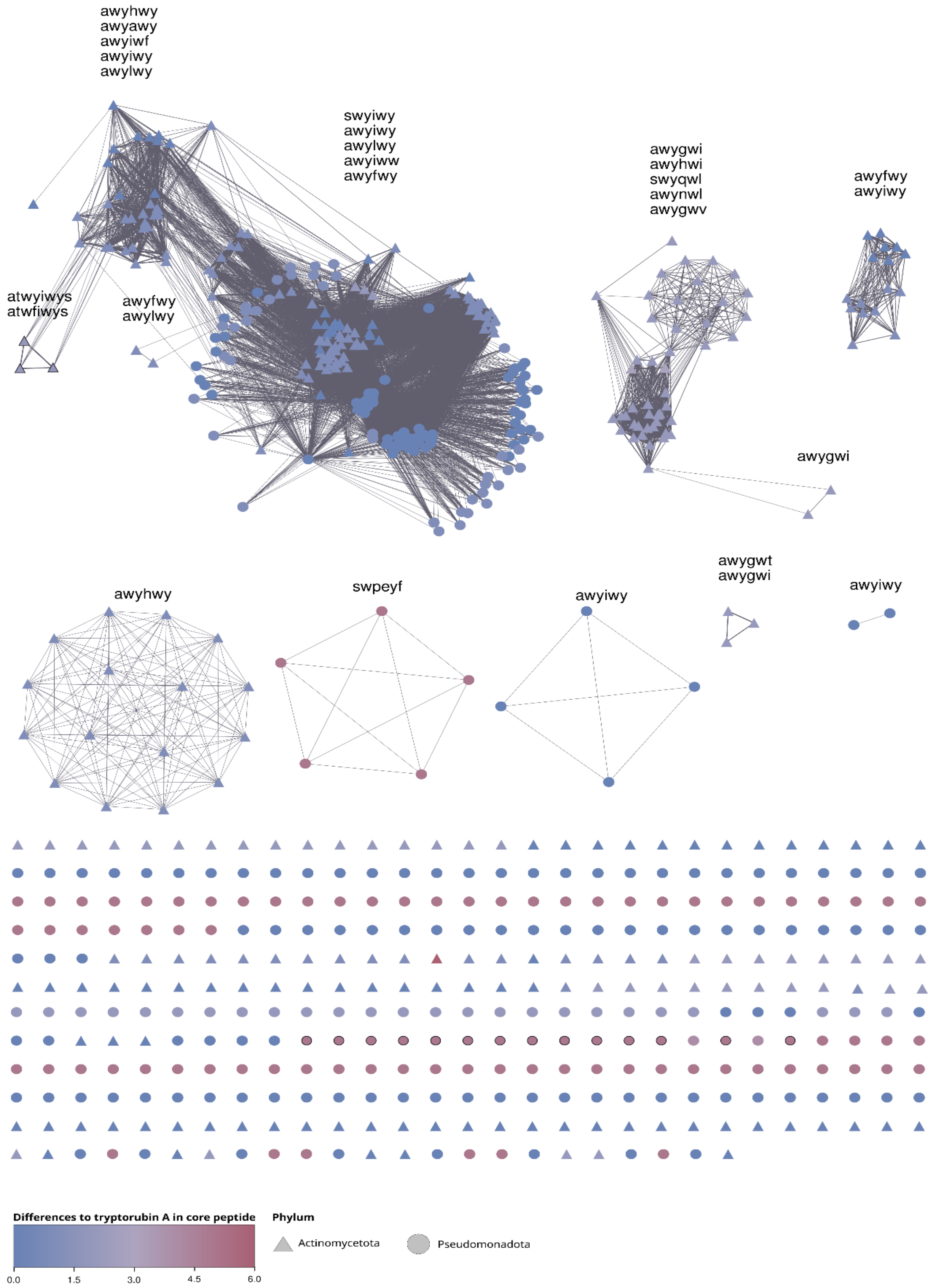


**Figure S9**. Sequence similarity network of all putative atropopeptide precursors and their corresponding core peptide sequences. The edge line width indicates the sequence similarity. Nodes with core peptides predicted to be longer than 6 amino acids are highlighted with a bold outline.

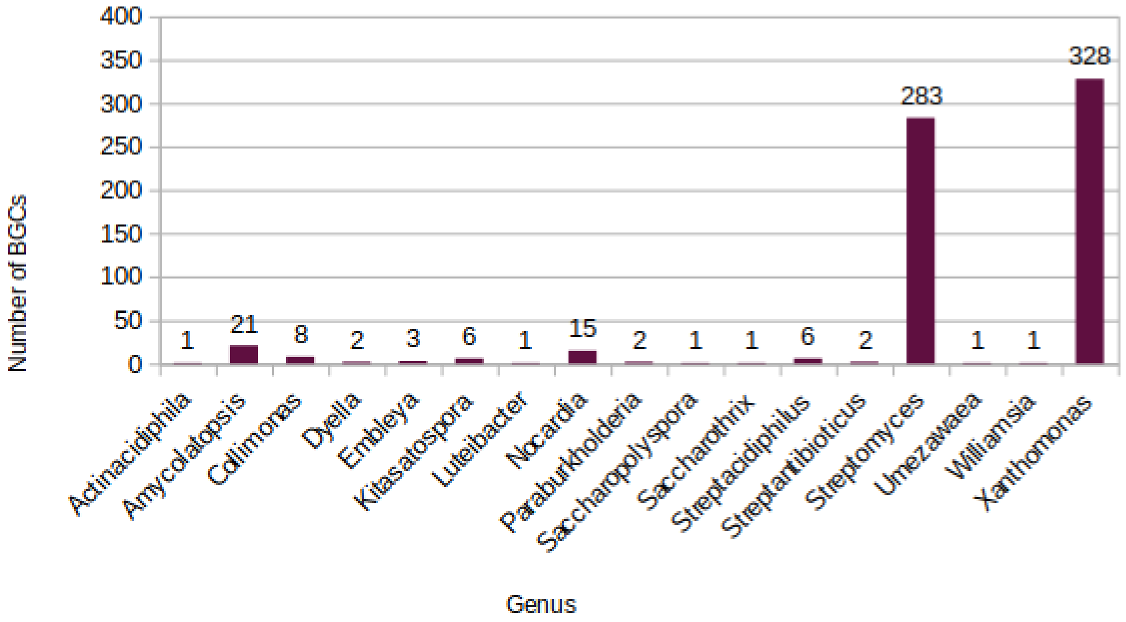


**Figure S10**. Phylogenetic distribution of putative atropopeptide clusters.

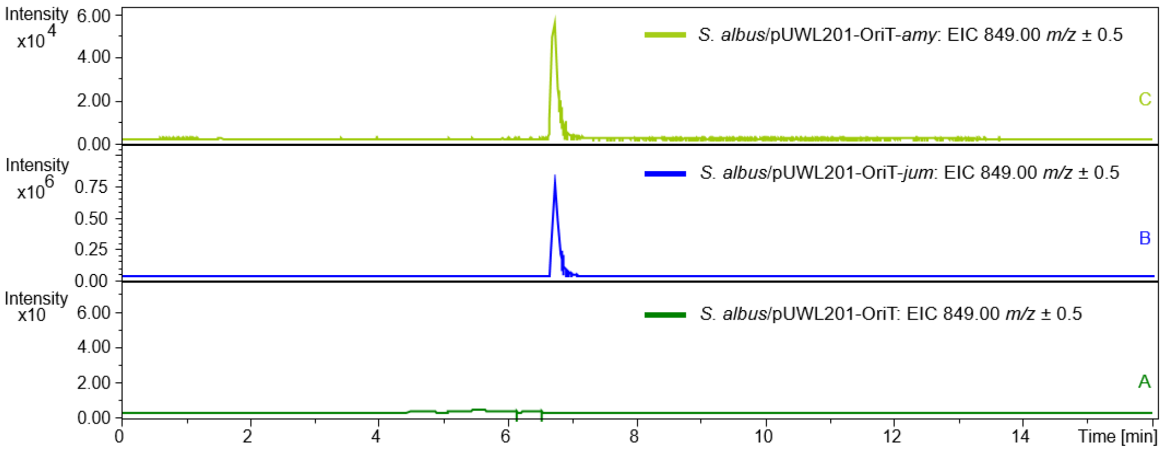


**Figure S11**. Extracted ion chromatograms of amyxirubin B and jumorubin (**2**). (A) *S. albus* harboring the empty pUWL201-oriT. (B) jumorubin produced by heterologous expression of the jumorubin BGC in *S. albus* J1074. (C) amyxirubin B produced by heterologous expression of the amyxirubin BGC in *S. albus* J1074.

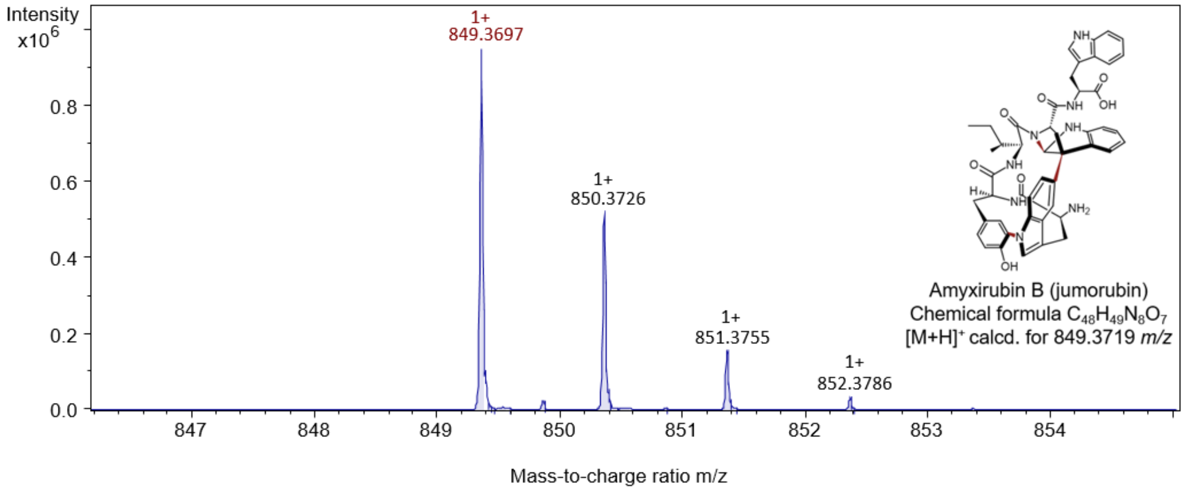


**Figure S12**. HPLC-ESI-QTOF-HRMS analysis of jumorubin (**2**).

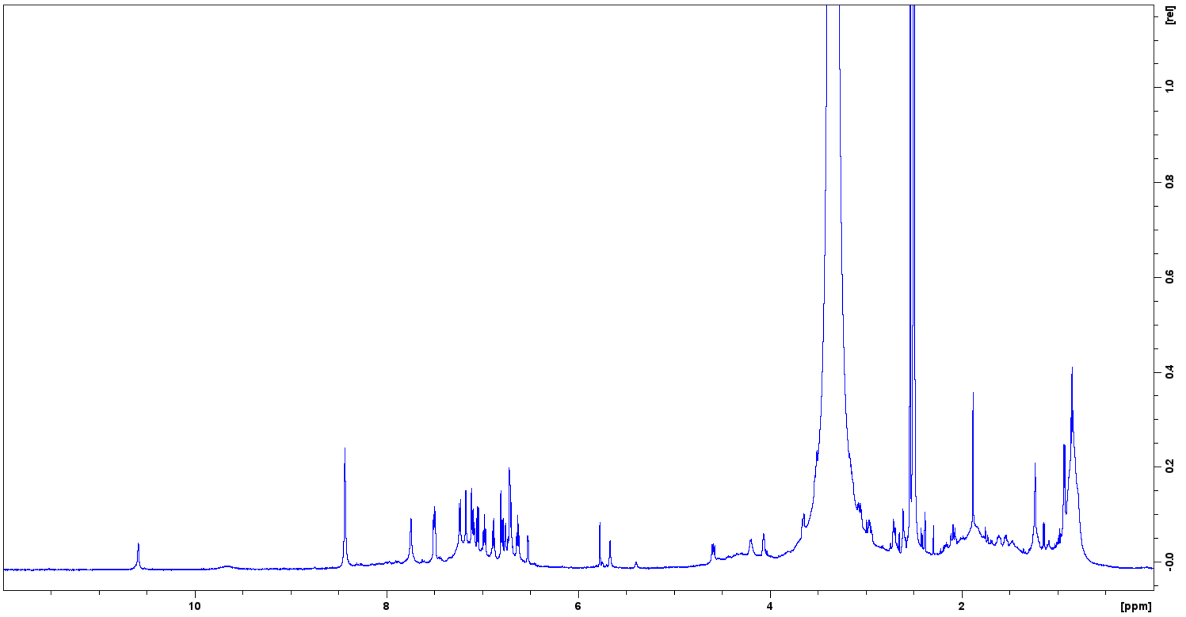


**Figure S13**. $^{1}H$ NMR spectrum (600 MHz) of jumorubin (**2**) in DMSO-$d_6$.

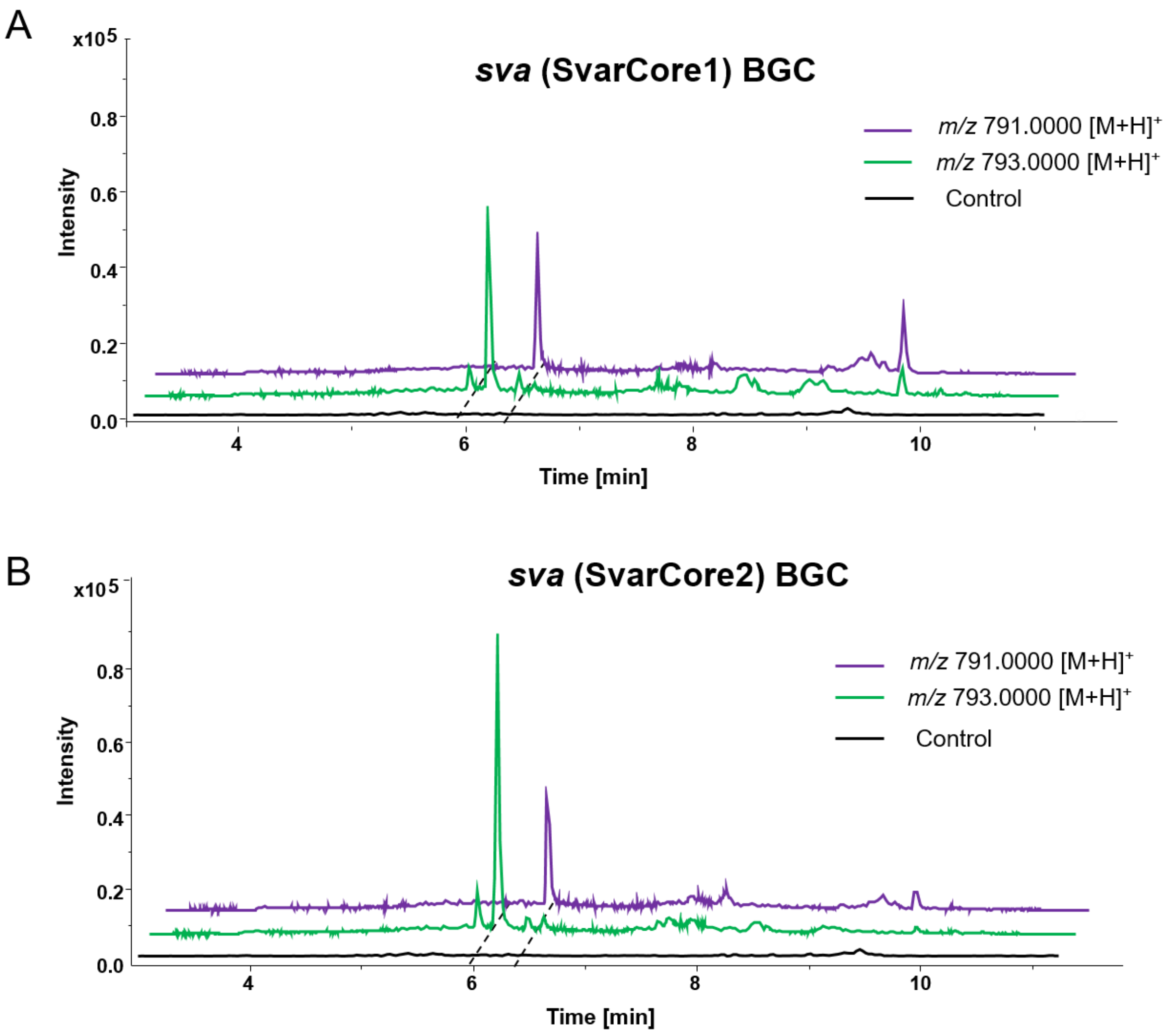


**Figure S14.** Extracted ion chromatogram (EIC) of varsitalin B1 (**3**) and varsitalin B2 (**4**) from different sources. (A) EIC of **3** and **4** from the control *S. albus* harboring the empty pUWL201-oriT and *S. albus* harboring pUWL201-oriT-SvarCore1. (B) EIC of **3** and **4** from the control *S. albus* harboring pUWL201-oriT and *S. albus* harboring pUWL201-oriT-SvarCore2.

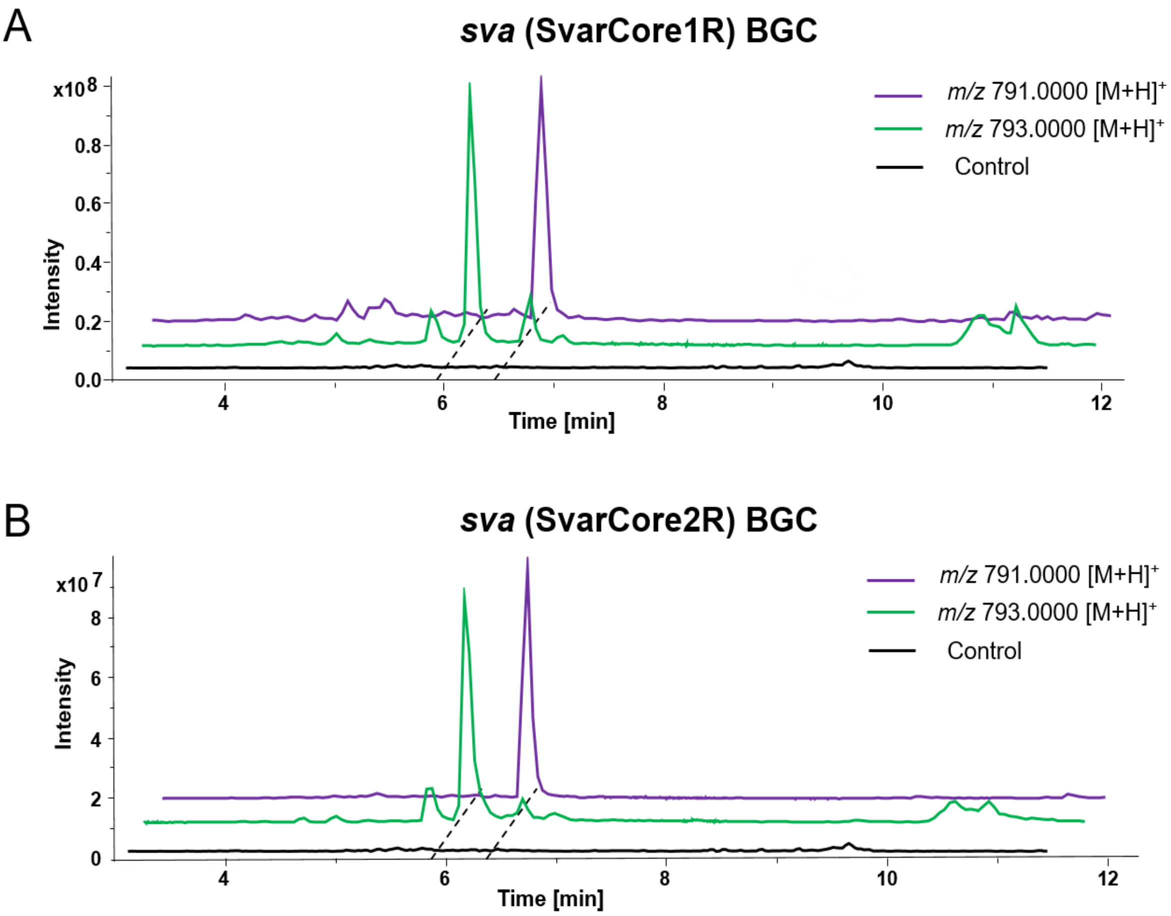


**Figure S15**. Extracted ion chromatogram (EIC) of varsitalin B1 (**3**) and varsitalin B2 (**4**)from different sources. (A) EIC of **3** and **4** from the control *S. albus* harboring pUWL201-oriT and *S. albus* harboring pUWL201-oriT-SvarCore1R. (B) EIC of **3** and **4** from the control *S. albus* harboring pUWL201-oriT and *S. albus* harboring pUWL201-oriT-SvarCore2R.

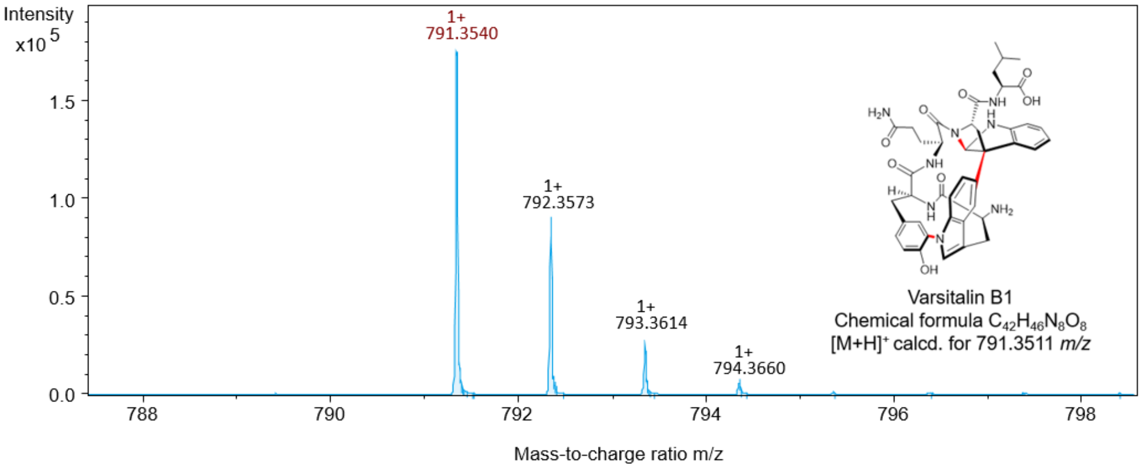


**Figure S16.** HPLC-ESI-QTOF-HRMS analysis of varsitalin B1 (**3**).

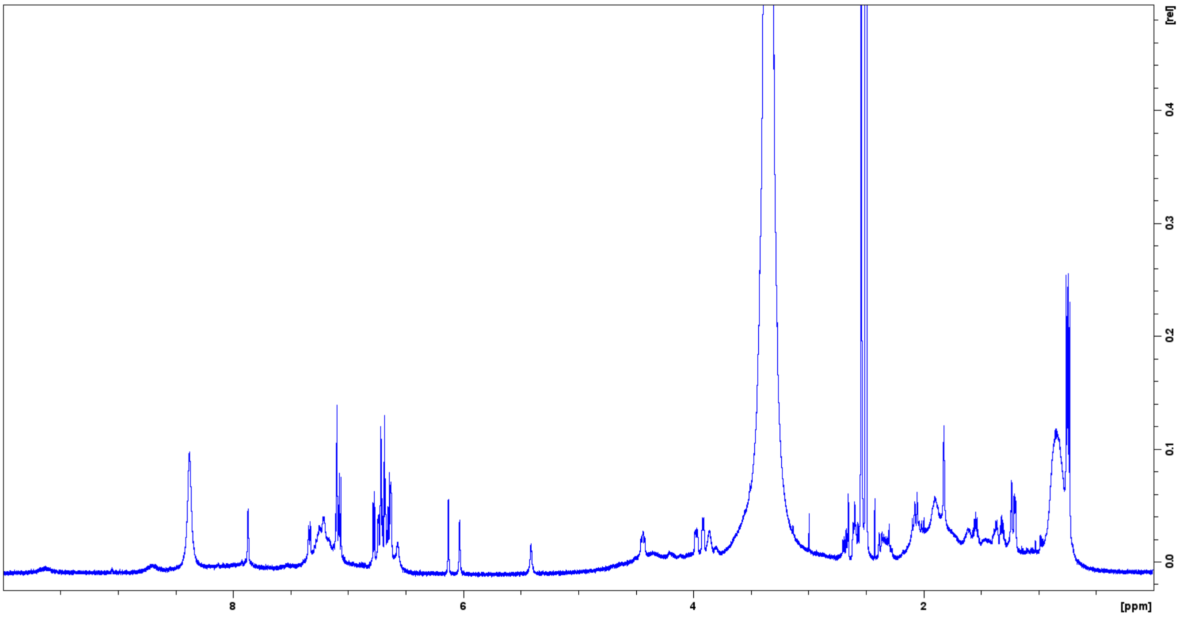


**Figure S17.** $^1$H NMR spectrum (600 MHz) of varsitalin B1 (**3**) in DMSO-$d_6$.

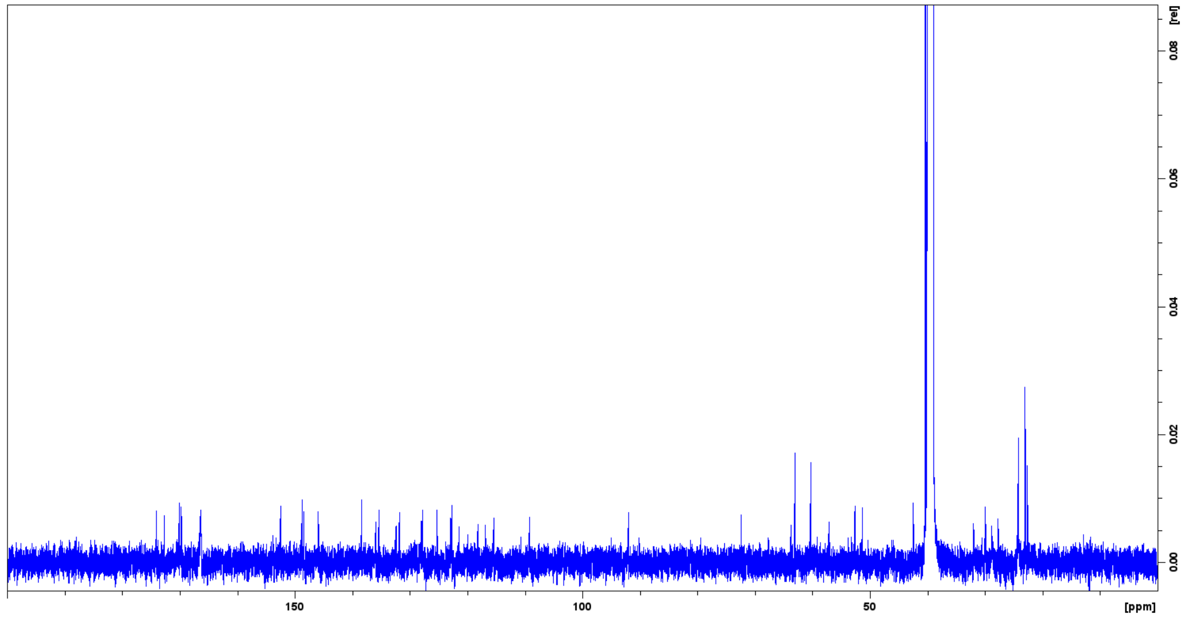


**Figure S18.** $^{13}C$ NMR spectrum (150 MHz) of varsitalin B1 (**3**) in DMSO-$d_6$.

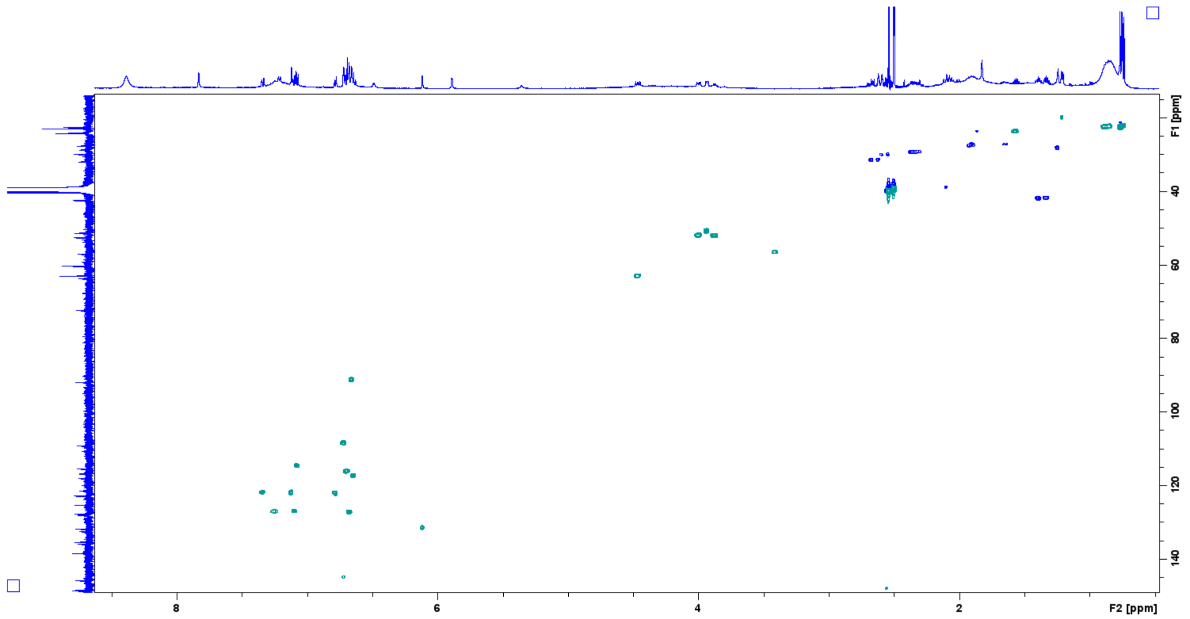


**Figure S19.** HSQC spectrum (600 MHz) of varsitalin B1 (**3**) in DMSO-$d_6$.

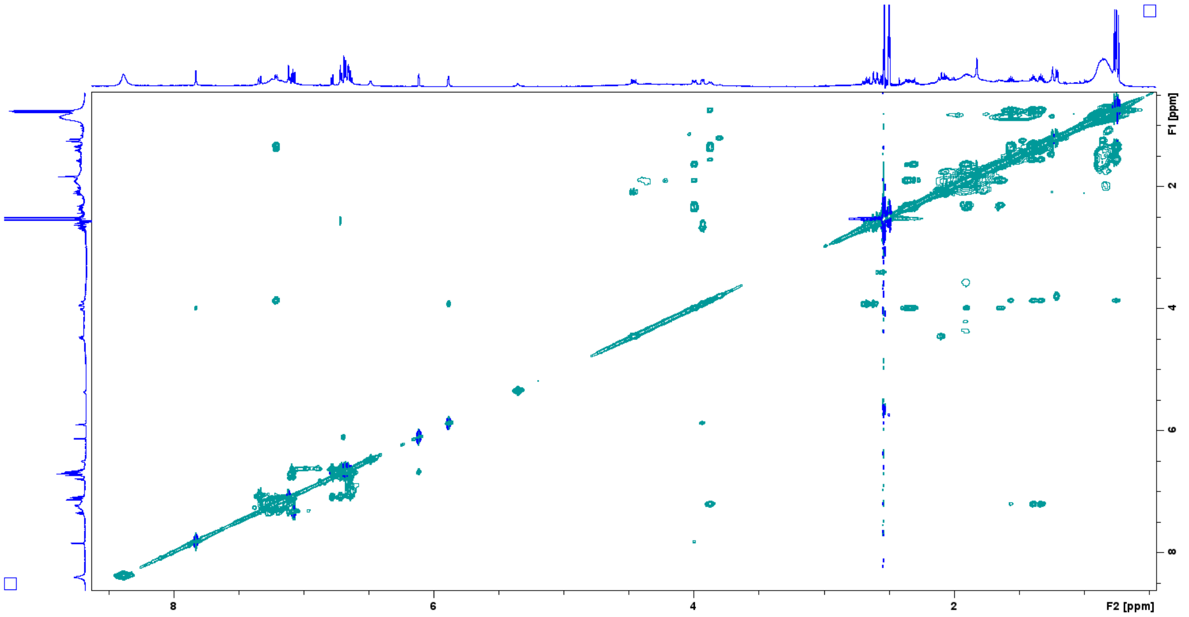


**Figure S20.** COSY spectrum (600 MHz) of varsitalin B1 (**3**) in DMSO-$d_6$.

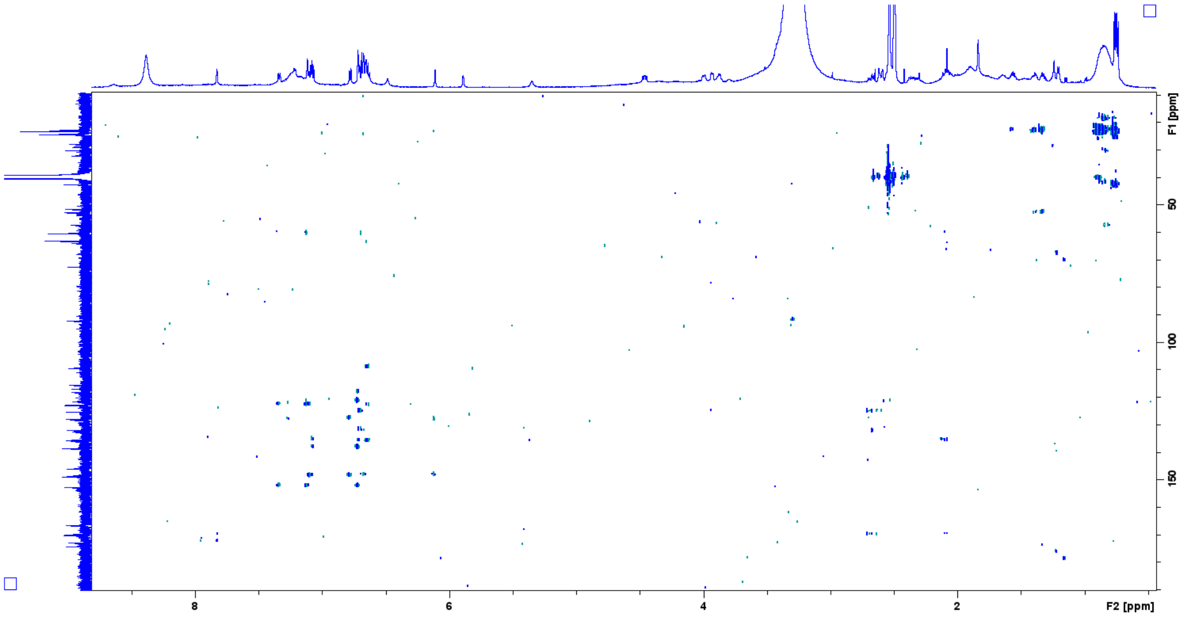


**Figure S21.** HMBC spectrum (600 MHz) of varsitalin B1 (**3**) in DMSO-$d_6$.

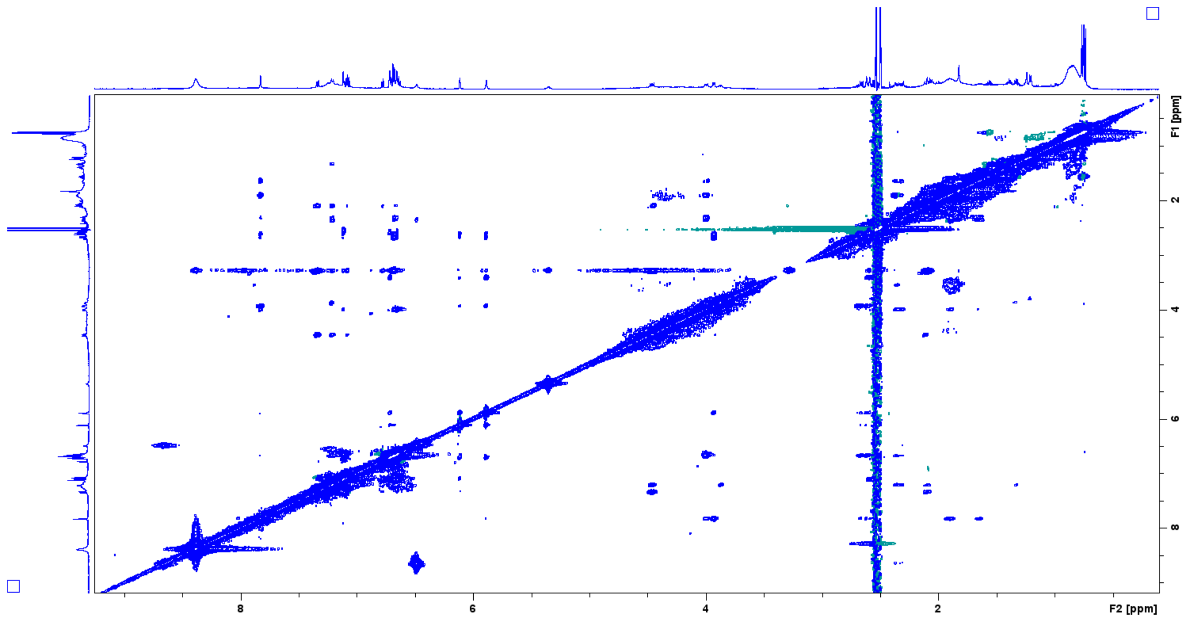


**Figure S22.** NOESY spectrum (600 MHz) of varsitalin B1 (**3**) in DMSO-$d_6$.

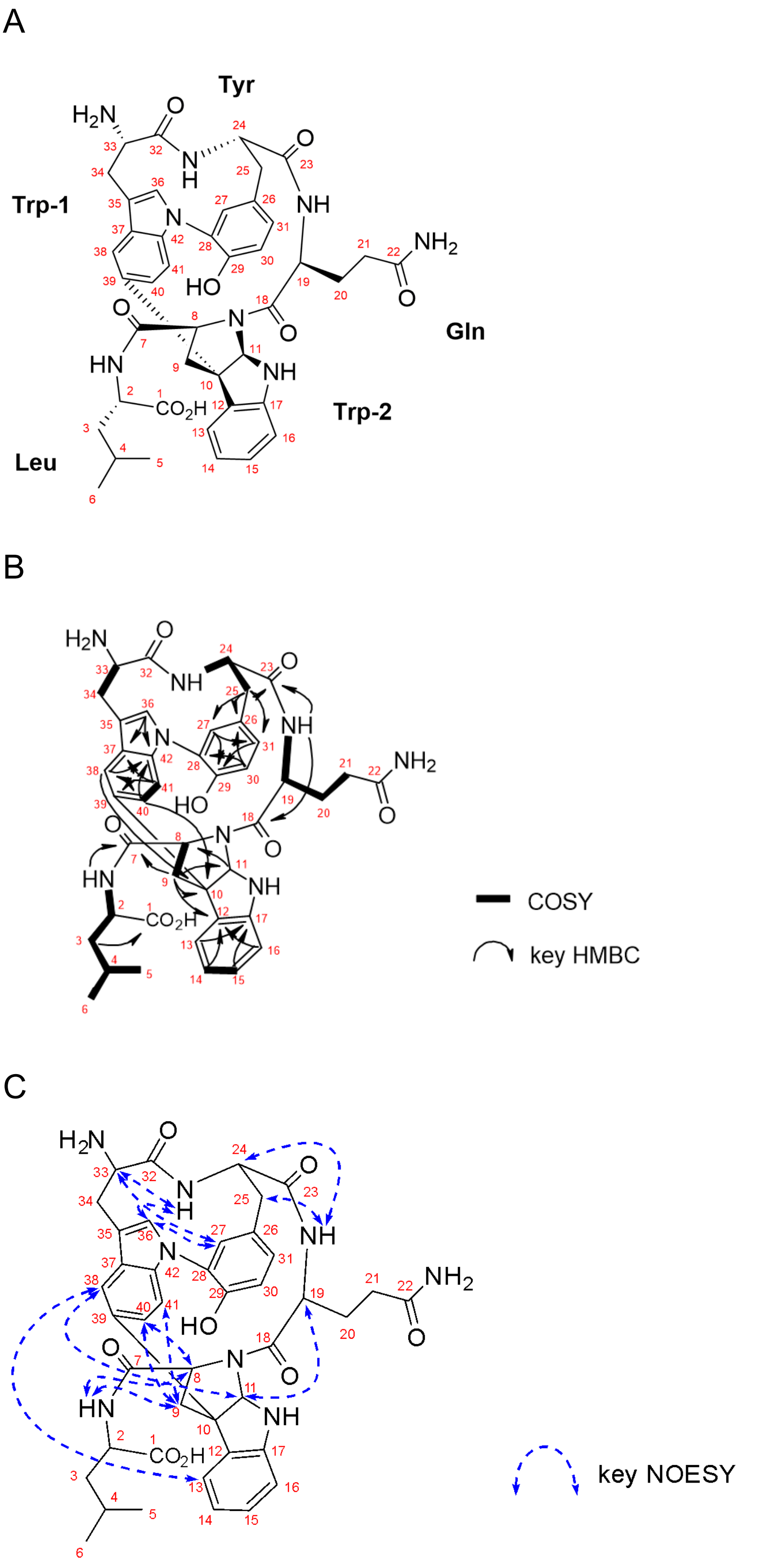


**Figure S23.** Structure elucidation of varsitalin B1 (**3**). (A) Structure of varsitalin B1 (**3**) and atom numbering used throughout this study. (B) COSY and key HMBC correlations in DMSO-$d_6$. (C) Key NOESY correlations of varsitalin B1 (**3**) in DMSO-$d_6$.

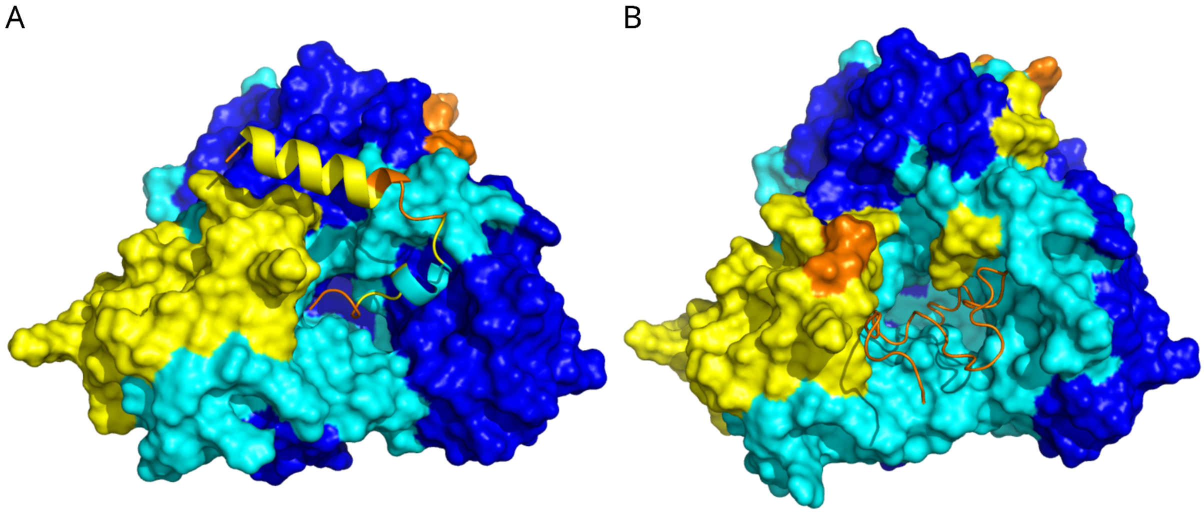


**Figure S24:** AlphaFold2 models of SvaB P450 with two putative precursor peptides SvaA1 and SvaA2. A) AlphaFold model of the P450 interacting with precursor peptide SvaA2. Contrary to the interaction with SvaA1, this model indicates a more favorable interaction mode in line with the system producing varsitalin B. This supports the hypothesis that only SvaA2 acts as the functional atropopeptide precursor peptide in the system. B) AlphaFold model of the svaB P450 interacting with precursor peptide SvaA1. The model predicts that SvaA1 interacts in such a way that its core peptide is not positioned within the P450's active site. The protein structures are color-coded based on pLDDT confidence scores: blue (>90), cyan (70-90), yellow (50-70), and orange (<50).

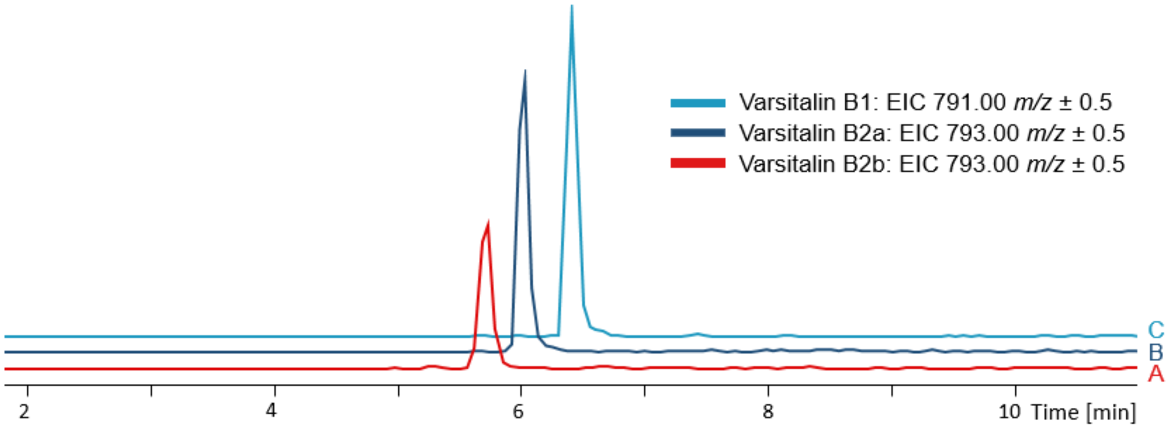


**Figure S25.** Extracted ion chromatograms of (A) varsitalin B2b (**4b**), (B) varsitalin B2a (**4a**) and (C) varsitalin B1 (**3**) from *S. albus* harboring pUWL201-oriT-SvarCore1R.

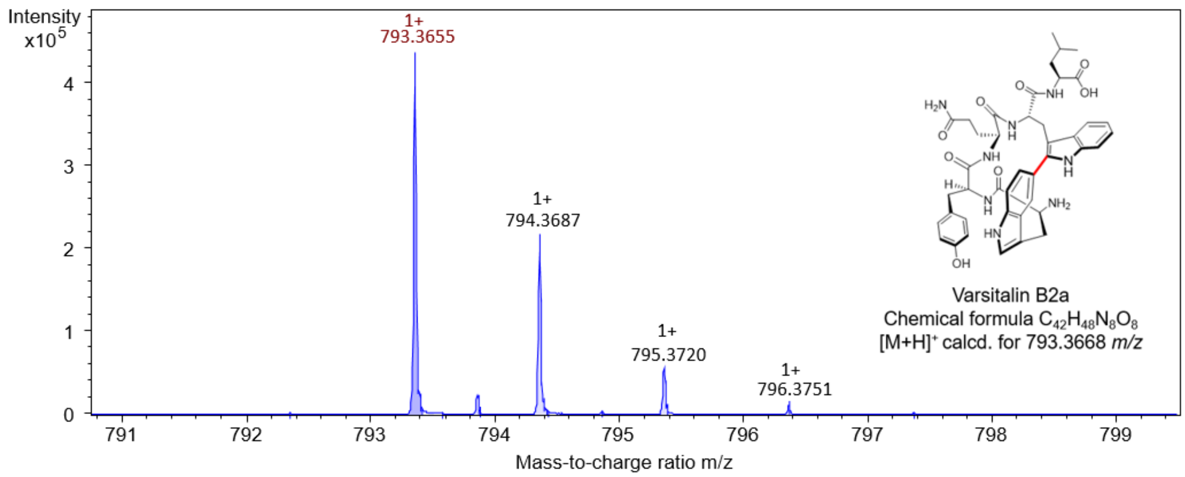


**Figure S26.** HPLC-ESI-QTOF-HRMS analysis of varsitalin B2a (**4a**).

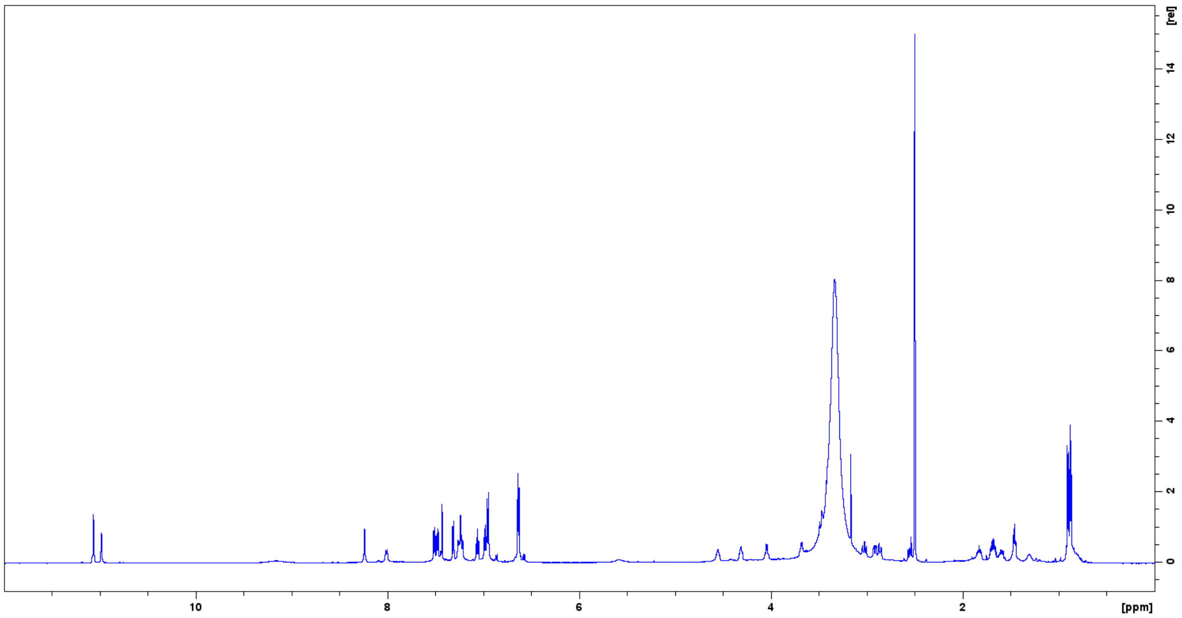


**Figure S27.** $^{1}H$ NMR spectrum (600 MHz) of varsitalin B2a (**4a**) in DMSO-$d_6$.

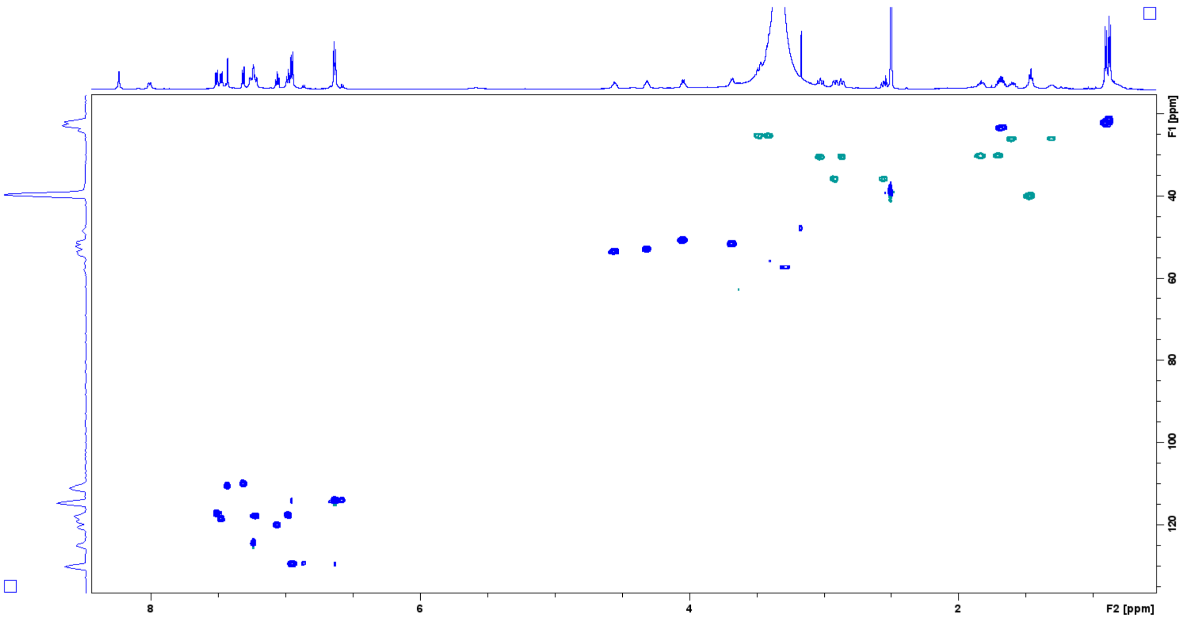


**Figure S28.** HSQC spectrum (600 MHz) of varsitalin B2a (**4a**) in DMSO-*d*$_6$.

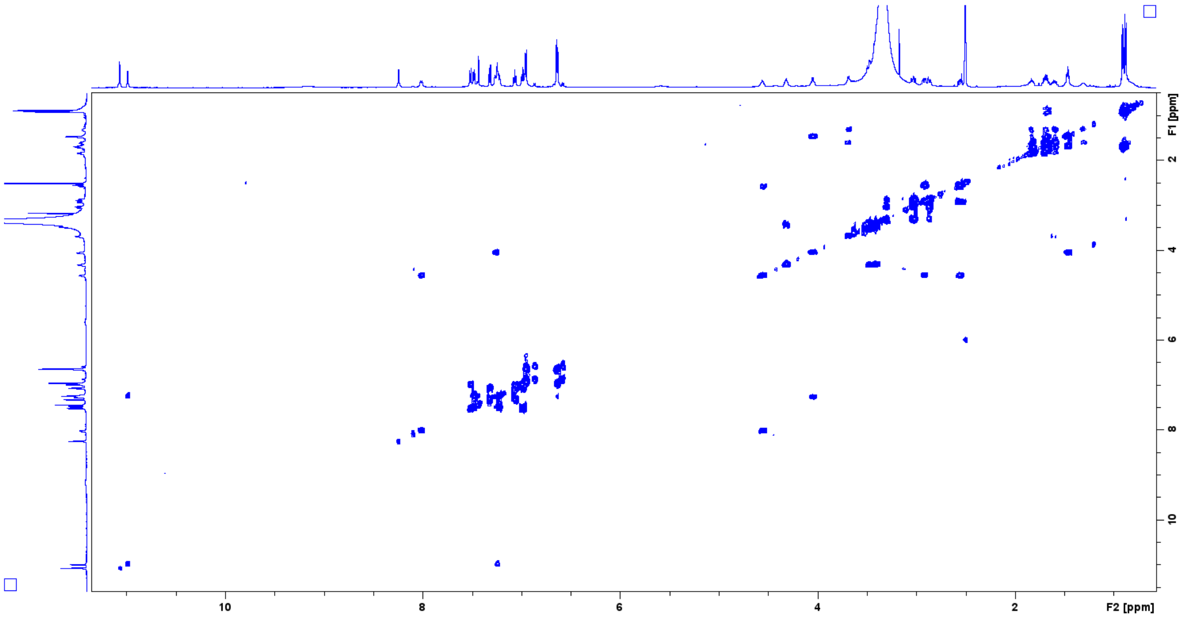


**Figure S29.** COSY spectrum (600 MHz) of varsitalin B2a (**4a**) in DMSO-*d*$_6$.

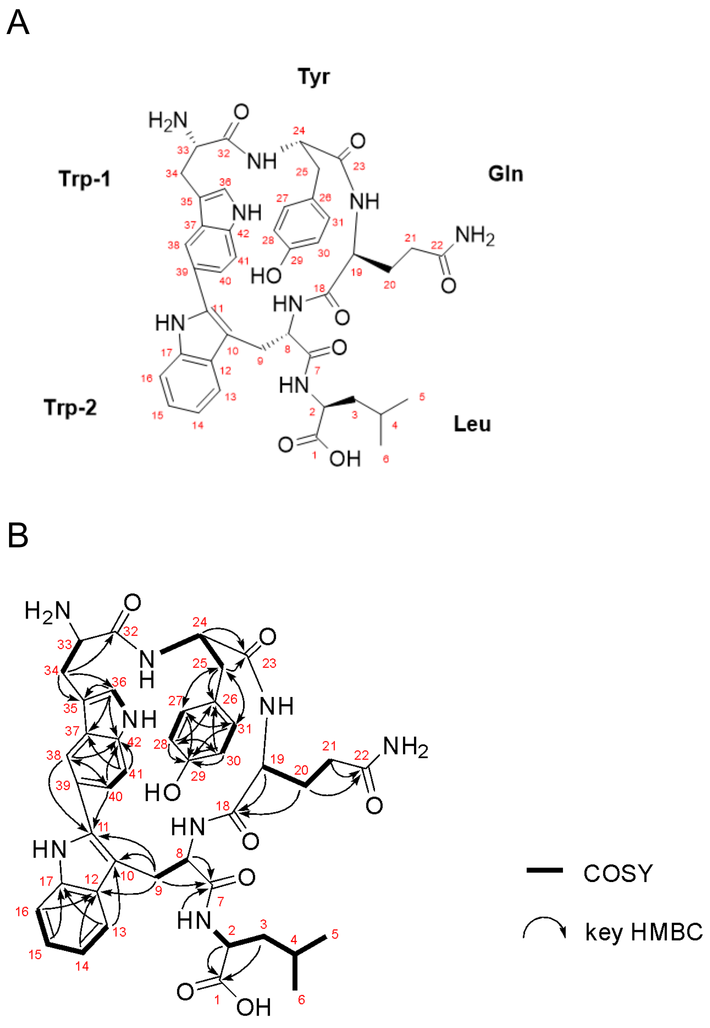


**Figure S30.** Structure elucidation of varsitalin B2a (**4a**). (A) Structure of varsitalin B2a (**4a**) and atom numbering used throughout this study. (B) COSY and key HMBC correlations in DMSO-$d_6$.

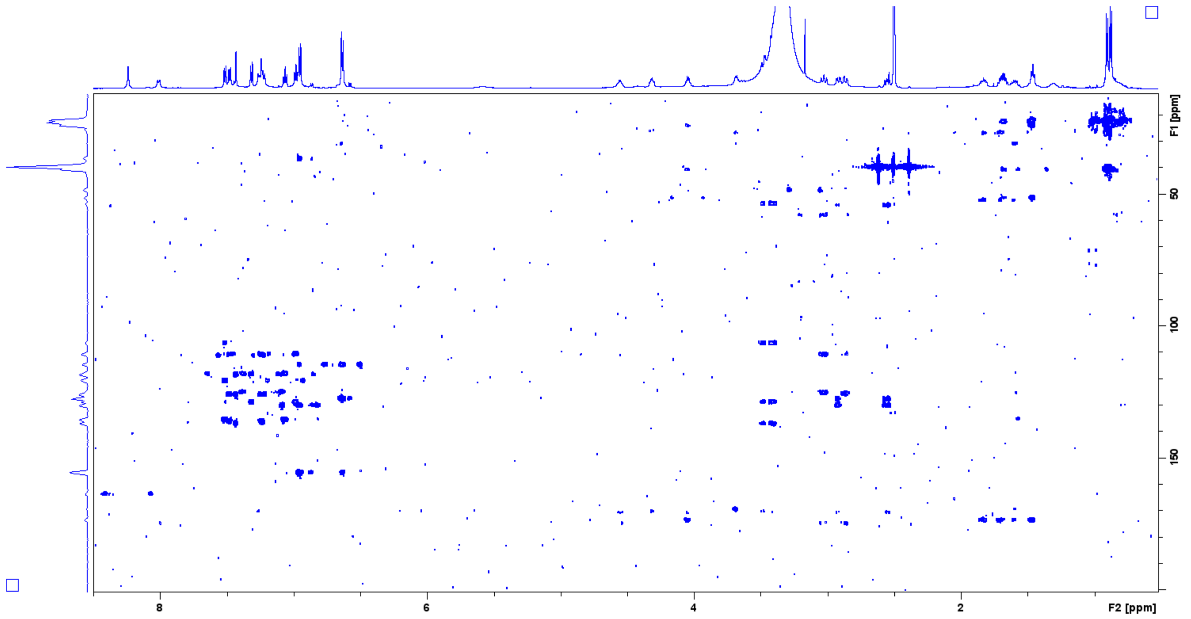


**Figure S31.** HMBC spectrum (600 MHz) of varsitalin B2a (**4a**) in DMSO-$d_6$.

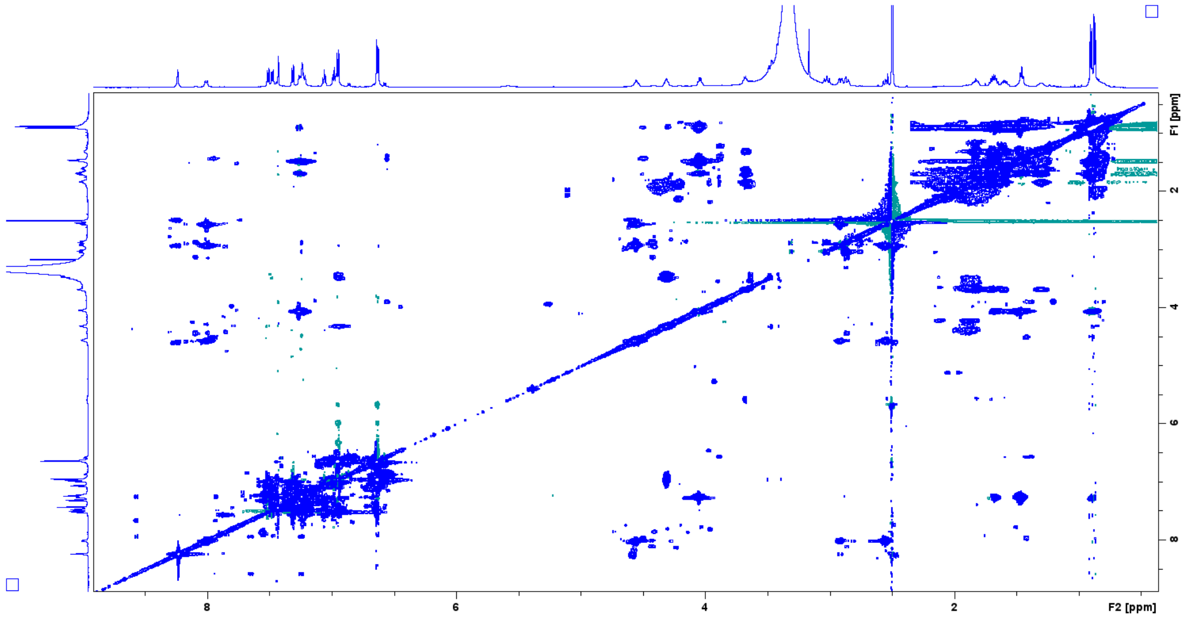


**Figure S32.** TOCSY spectrum (600 MHz) of varsitalin B2a (**4a**) in DMSO-$d_6$.

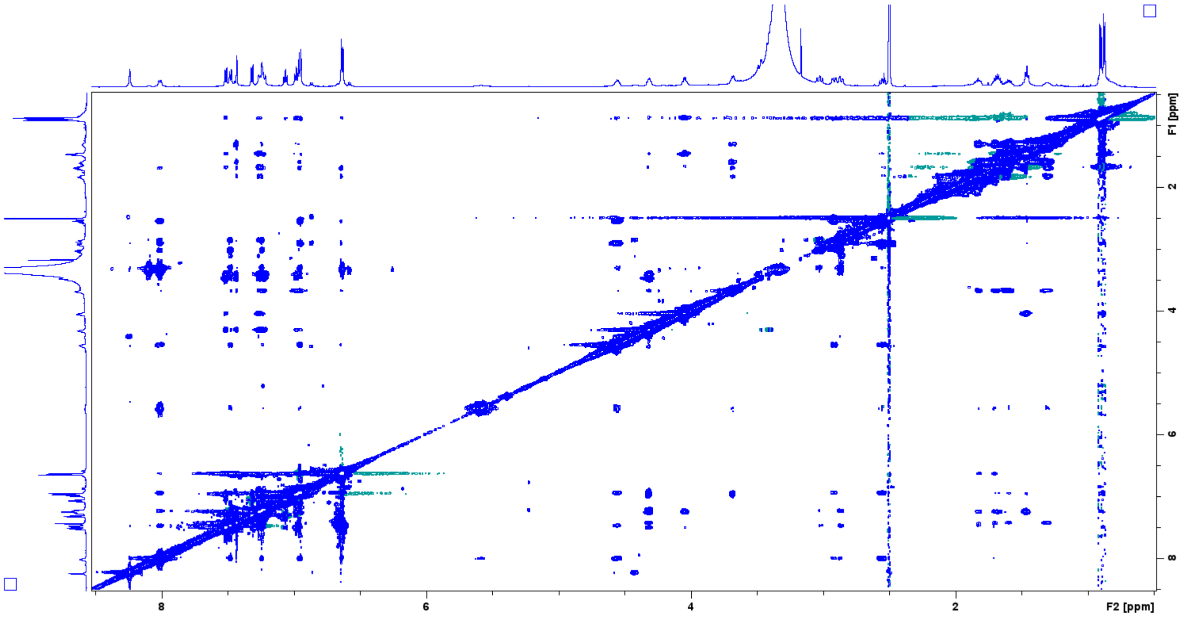


**Figure S33.** NOESY spectrum (600 MHz) of varsitalin B2a (**4a**) in DMSO-$d_6$.

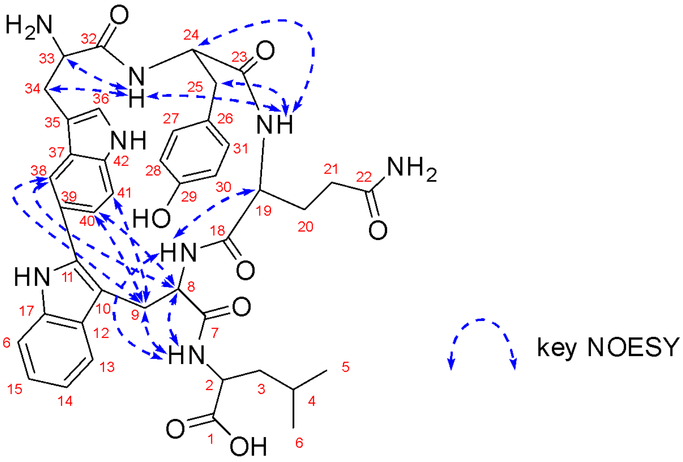


**Figure S34.** Key NOESY correlations of varsitalin B2a (**4a**) in DMSO-$d_6$.

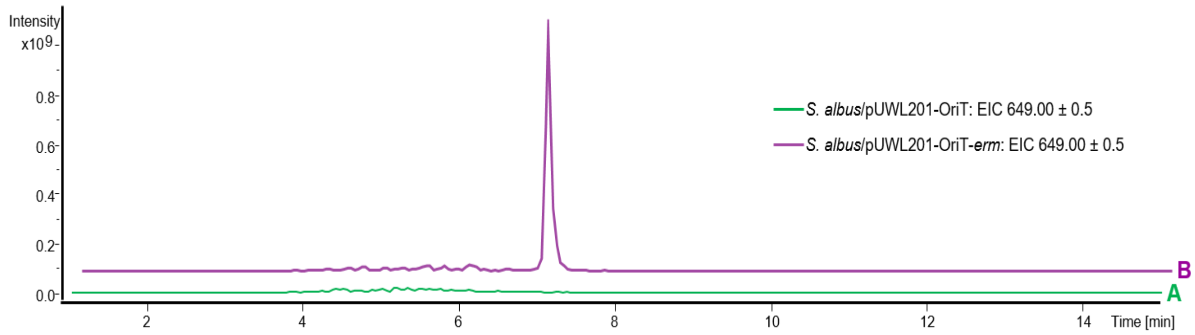


**Figure S35.** Extracted ion chromatogram of embyscamide (**5**) from *S. albus* harboring pUWL201-oriT (A) and *S.albus* harboring pUWL201-OriT-*erm* (B).

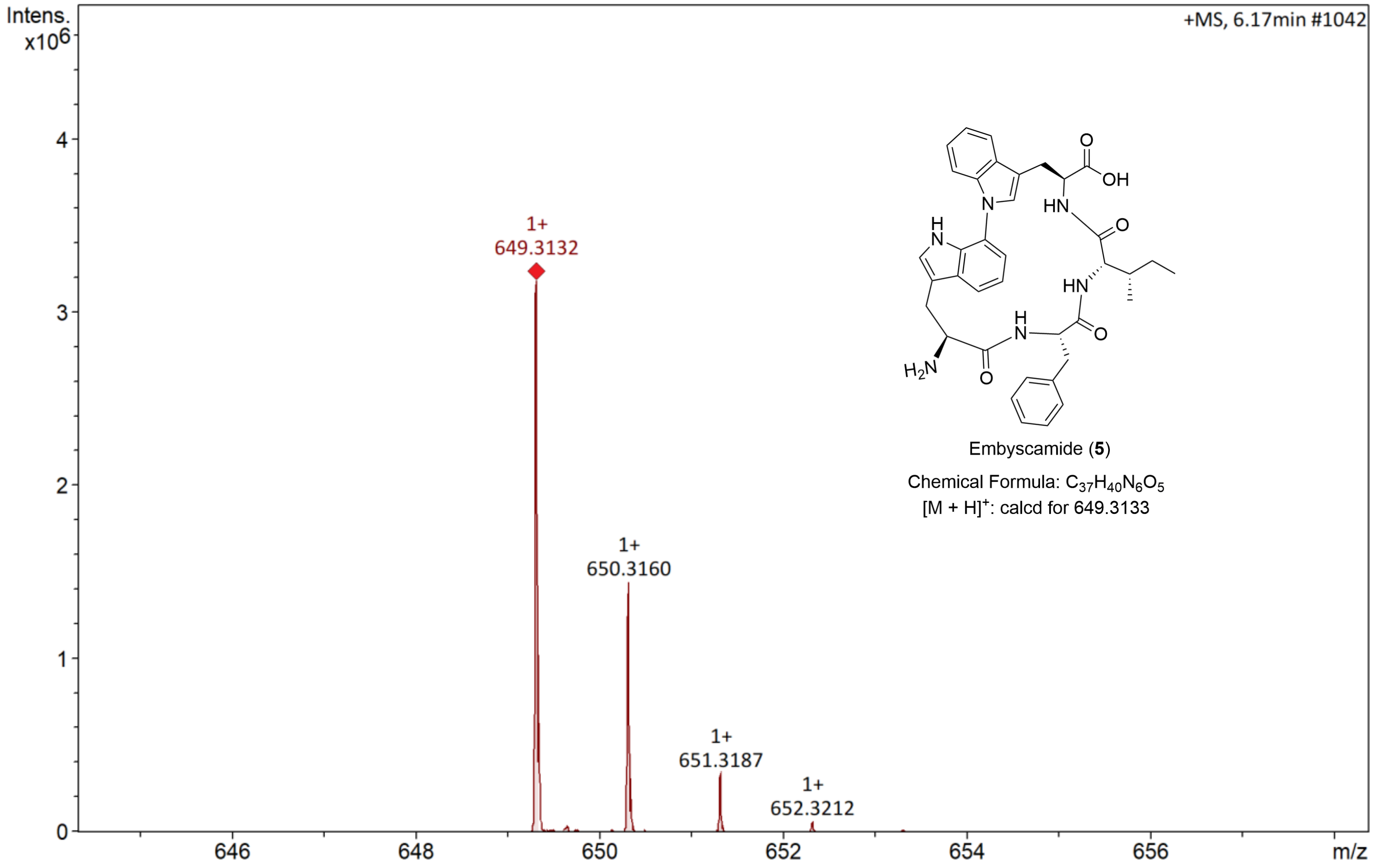


**Figure S36.** HPLC-ESI-QTOF-HRMS analysis of embyscamide (**5**).

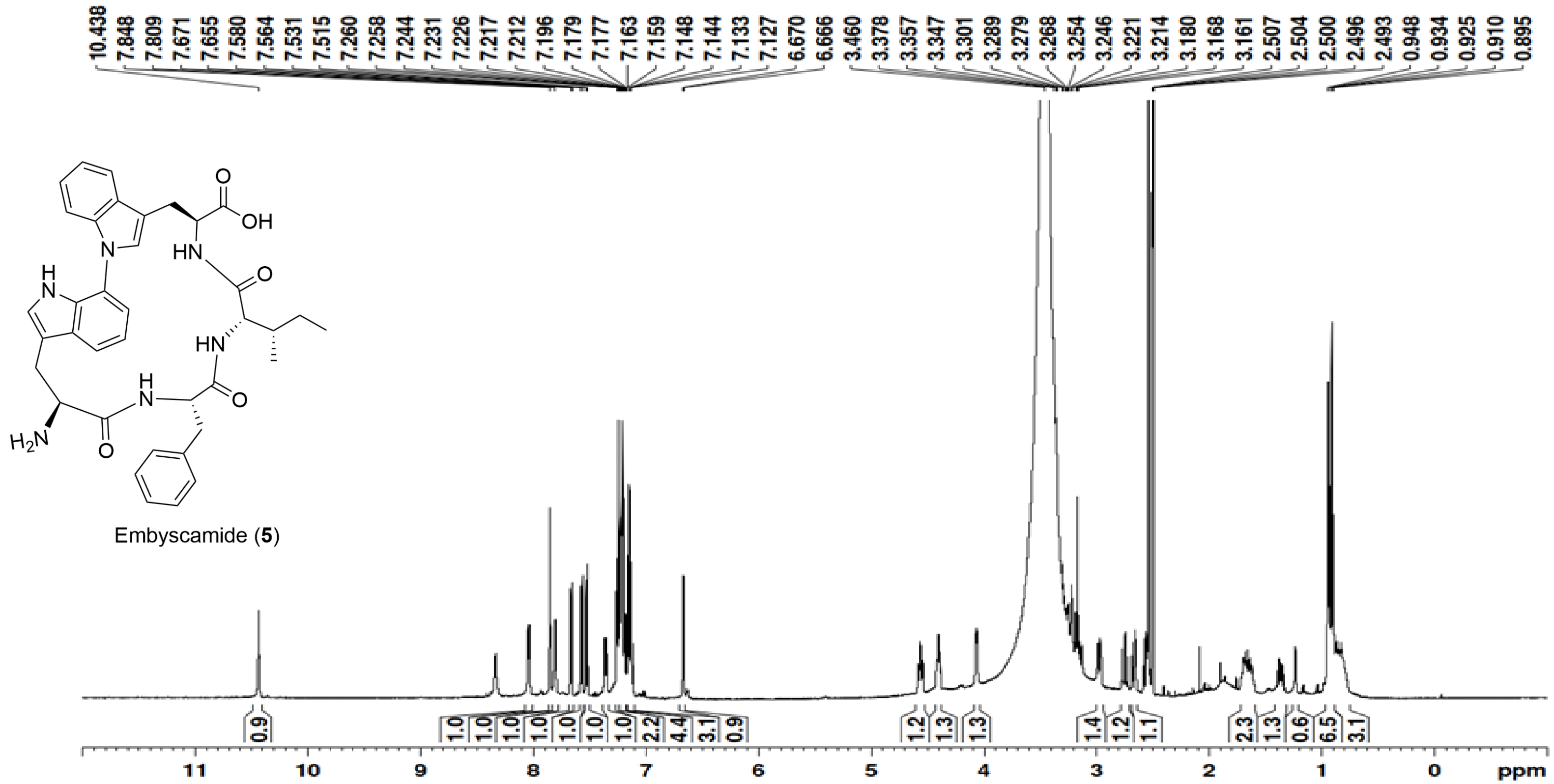


**Figure S37.** $^1$H-NMR spectrum (500 MHz) of embyscamide (**5**) in DMSO-$d_6$.

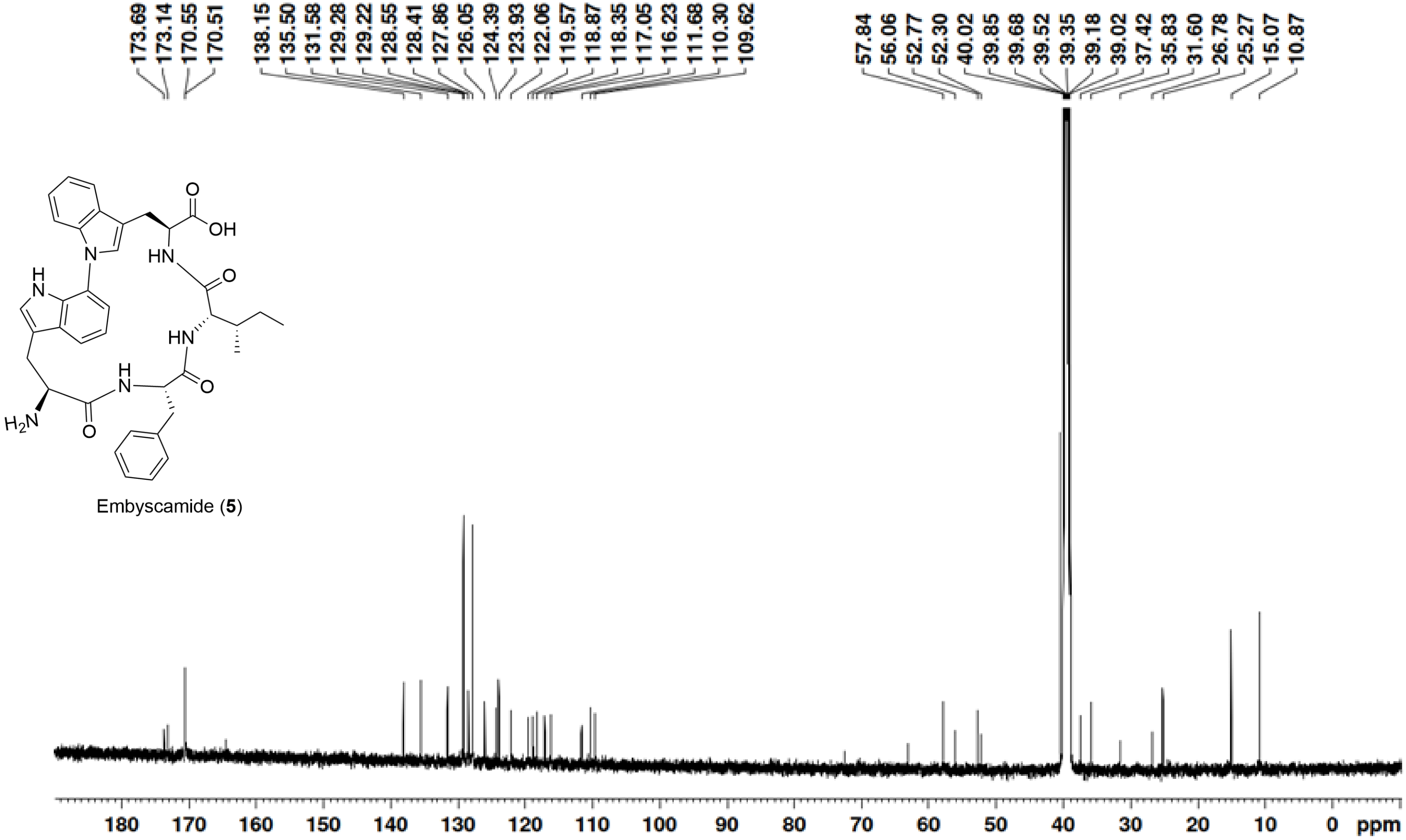


**Figure S38.** $^{13}$C-NMR spectrum (125 MHz) of embyscamide (**5**) in DMSO-$d_6$.

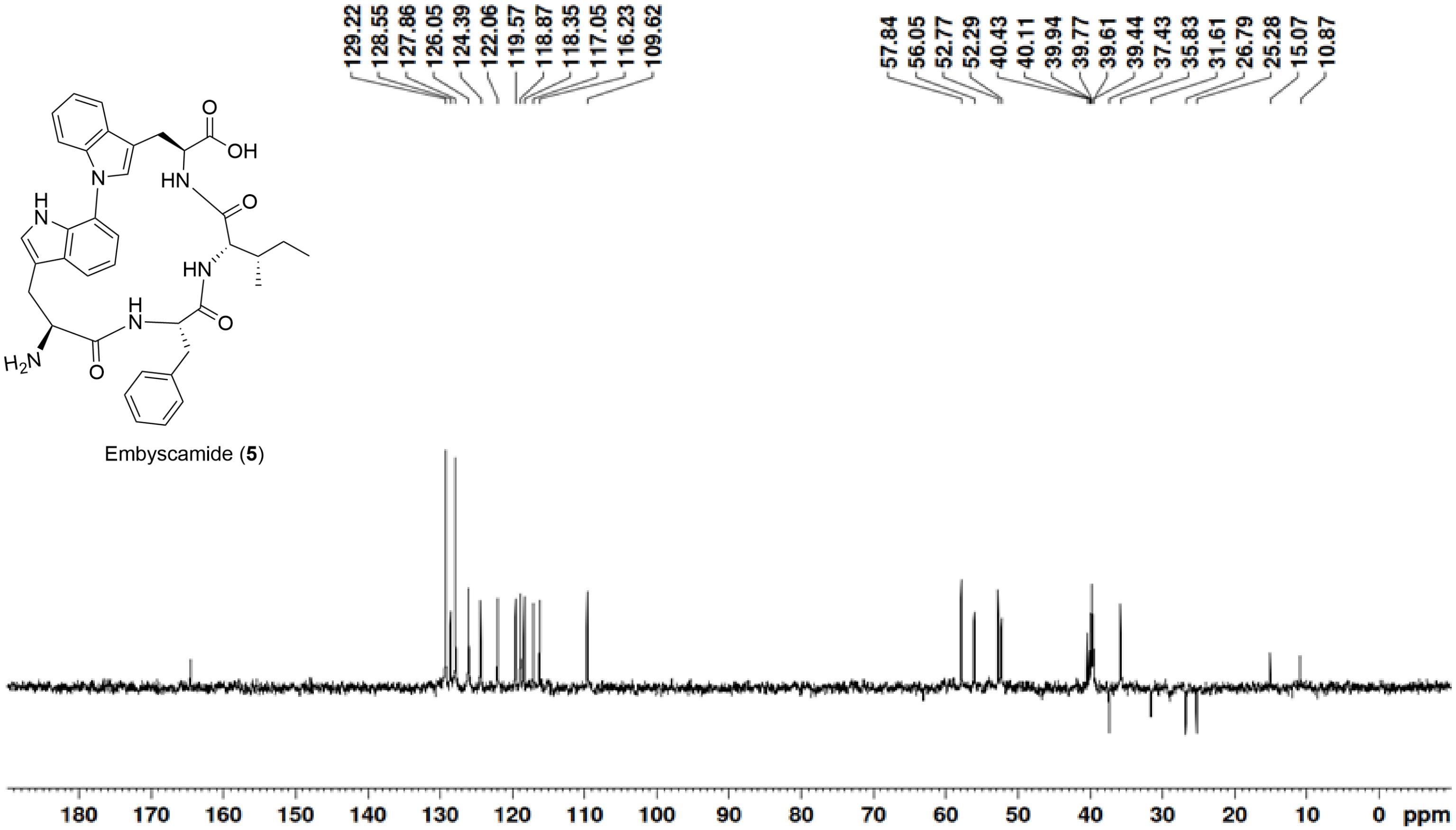


**Figure S39.** DEPT135 spectrum (125 MHz) of embyscamide (**5**) in DMSO-$d_6$.

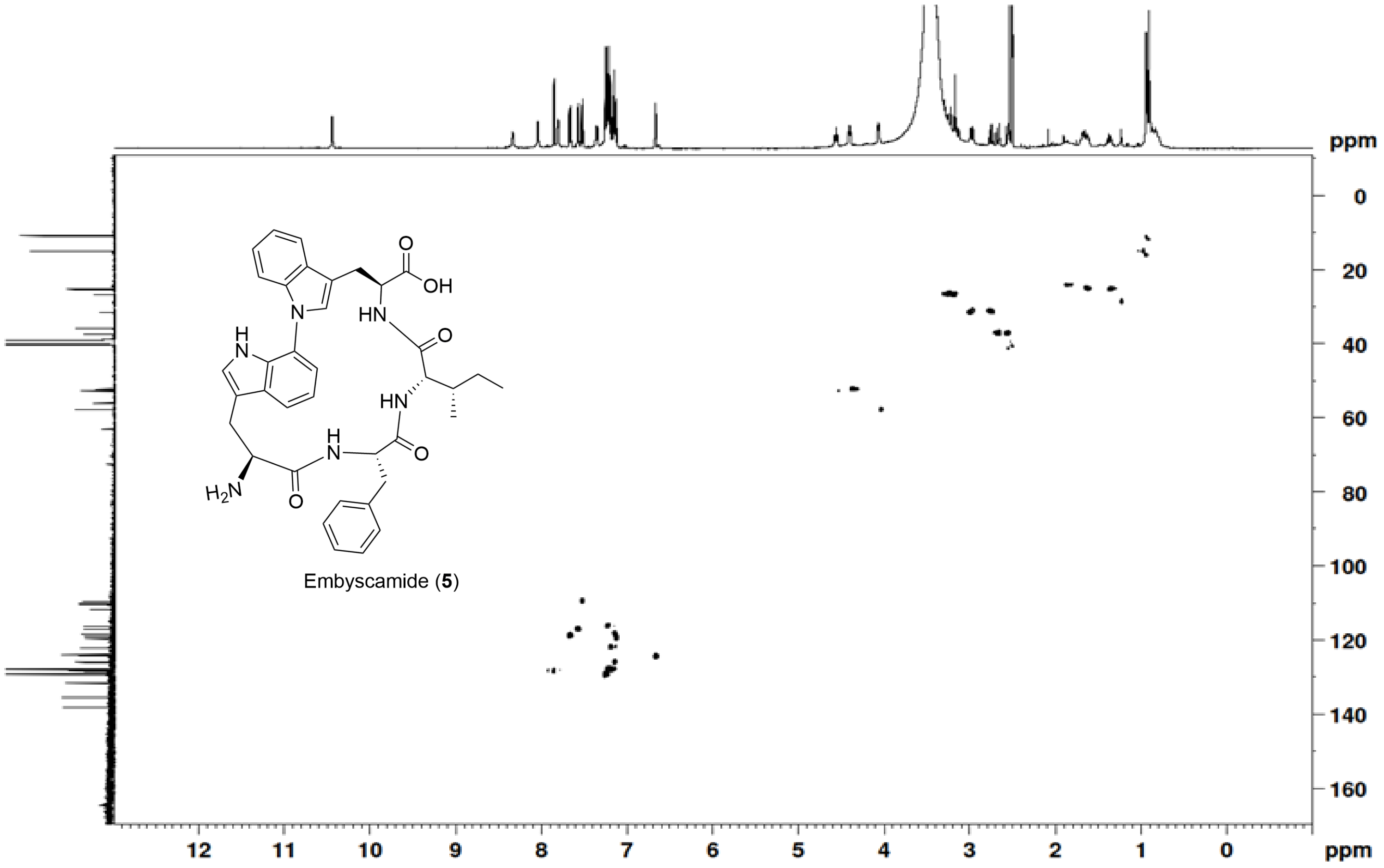


**Figure S40.** HSQC spectrum (500 MHz) of embyscamide (**5**) in DMSO-$d_6$.

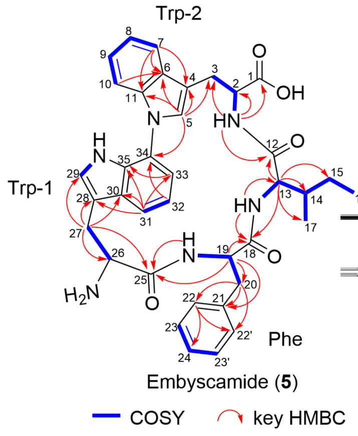


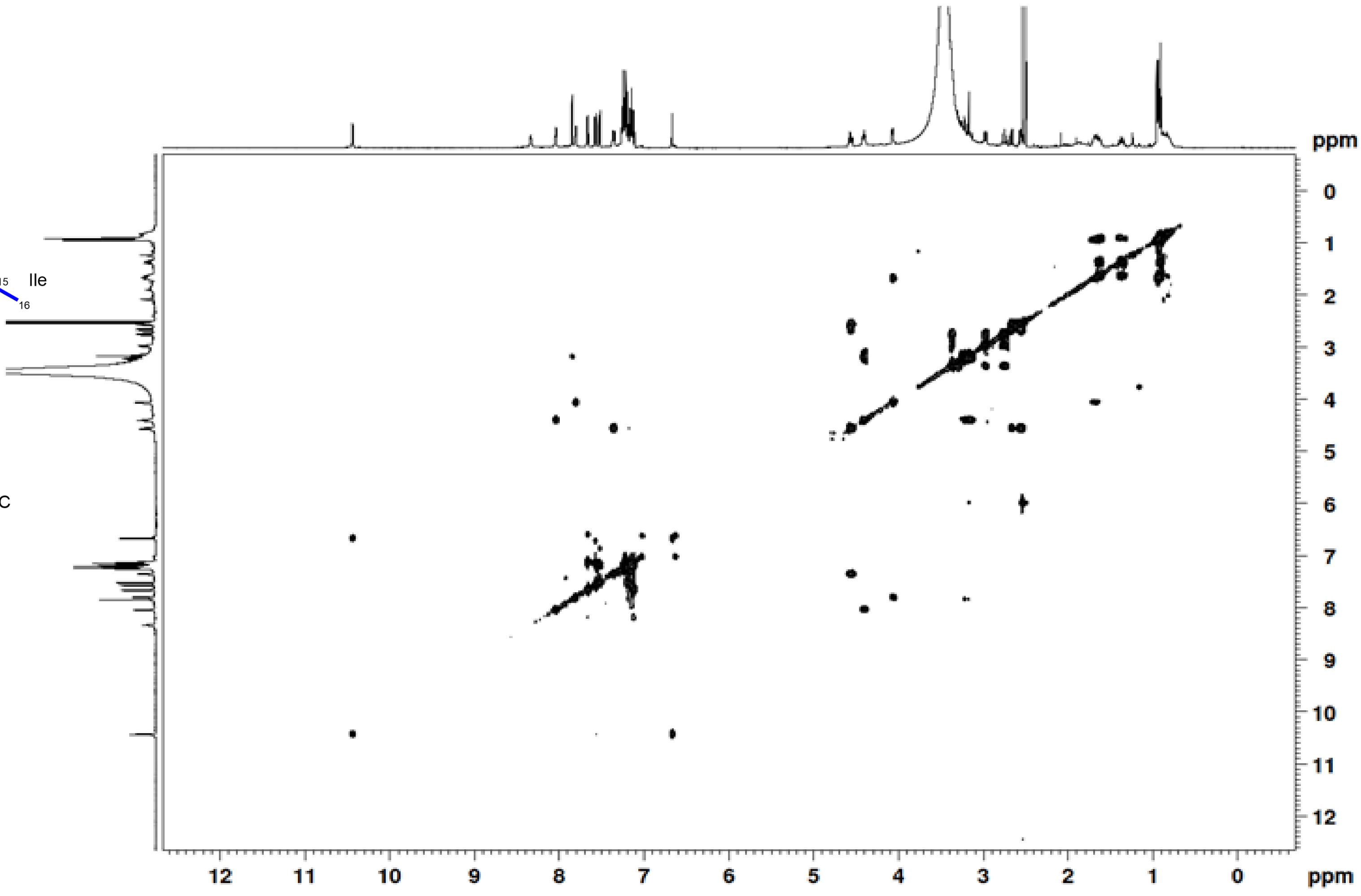


**Figure S41.** COSY spectrum (500 MHz) of embyscamide (**5**) in DMSO-$d_6$.

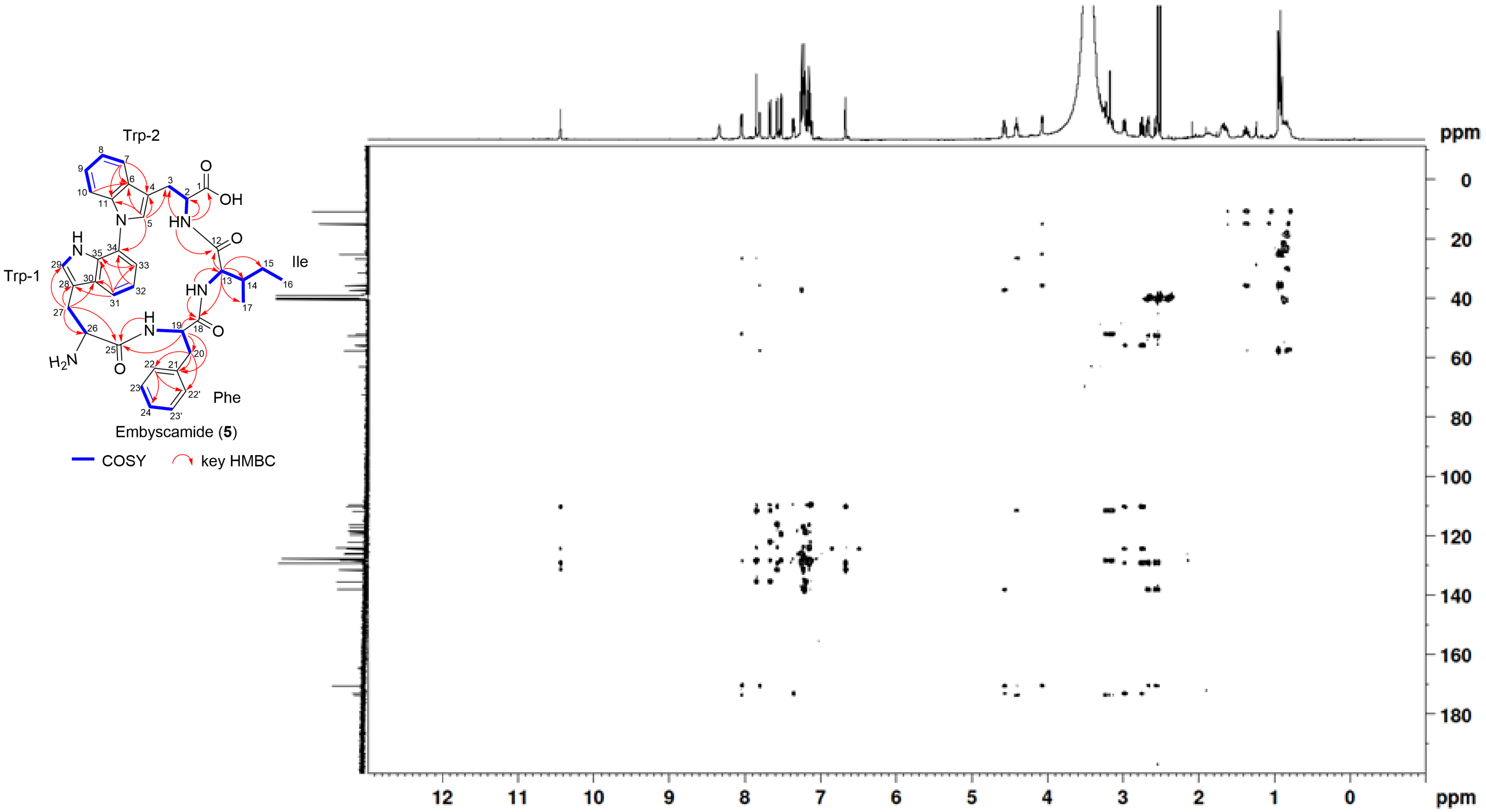


**Figure S42.** HMBC spectrum (500 MHz) of embyscamide (**5**) in DMSO-$d_6$.

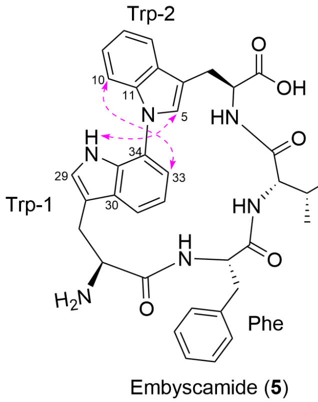


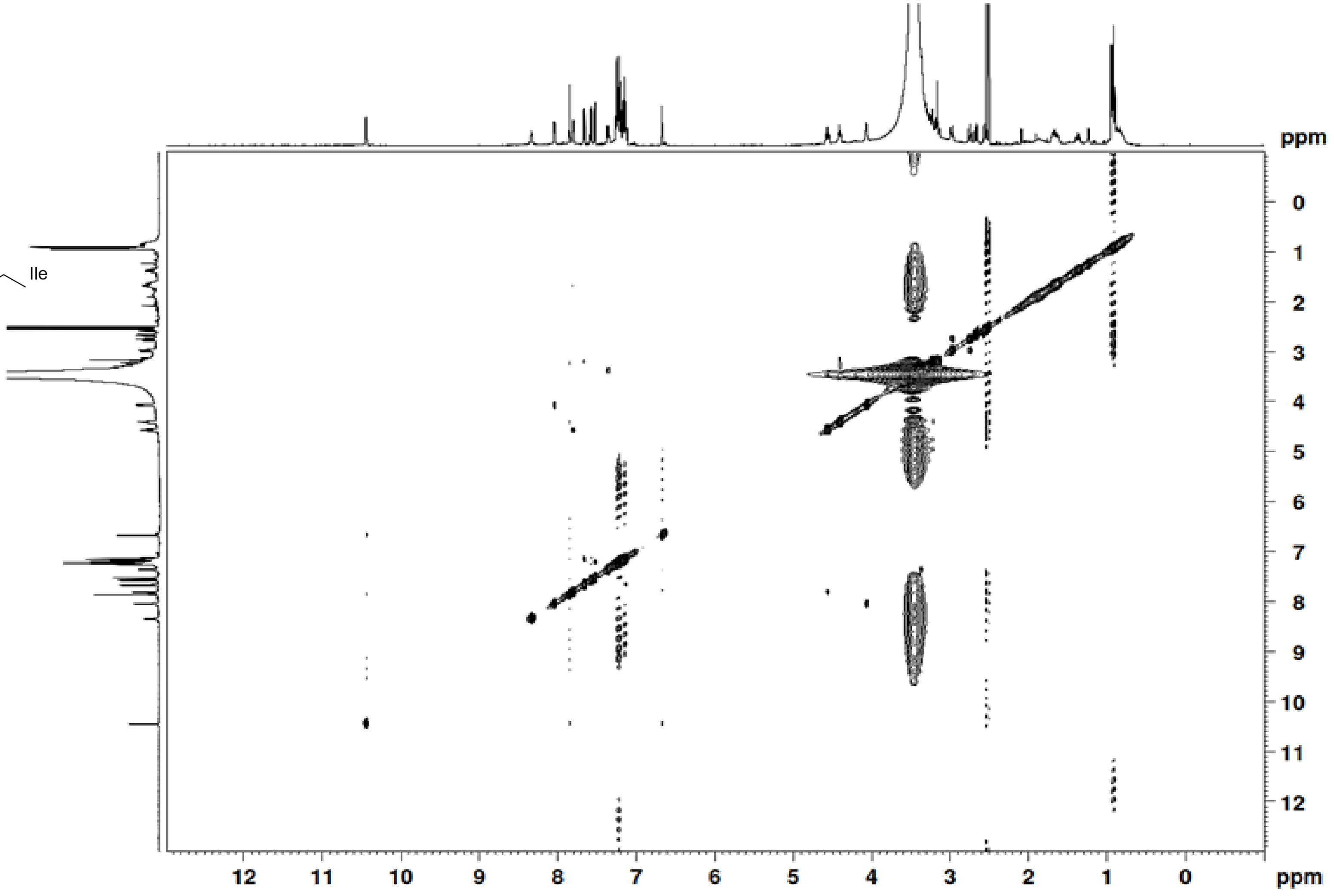


**Figure S43.** NOESY spectrum (500 MHz) of embyscamide (**5**) in DMSO-$d_6$.

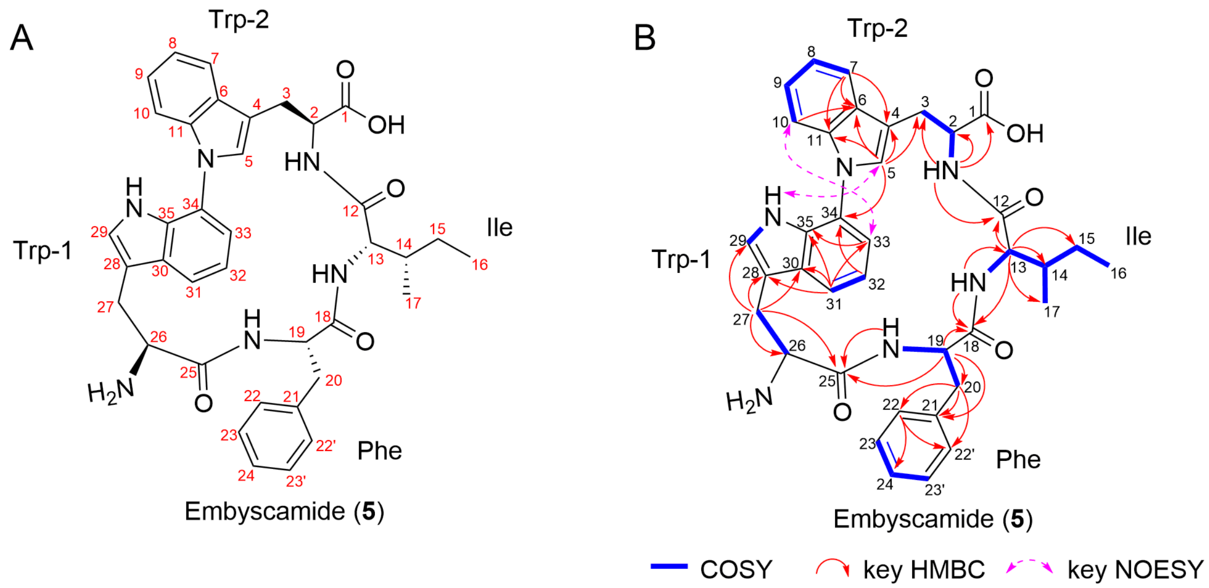


**Figure S44.** Structure elucidation of embyscamide (**5**). (A) Structure of **5** and atom numbering used throughout this study. (B) Key $^{1}H$-$^{1}H$ COSY, $^{1}H$-$^{13}C$ HMBC and $^{1}H$-$^{1}H$ NOESY correlation used for determining the structure of **5**.

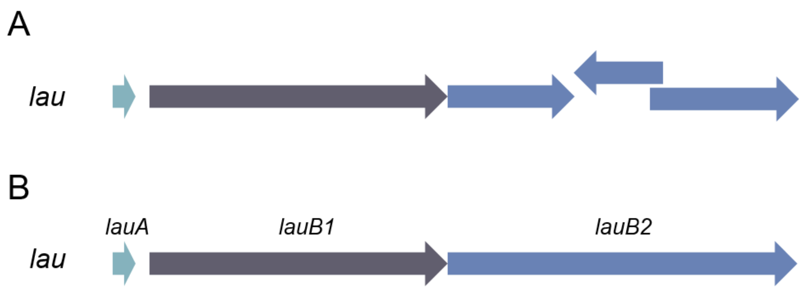


**Figure S45**. Resequencing of *lau* BGC. (A) the *lau* BGC downloaded from NCBI; (B) the resequenced *lau* BGC.

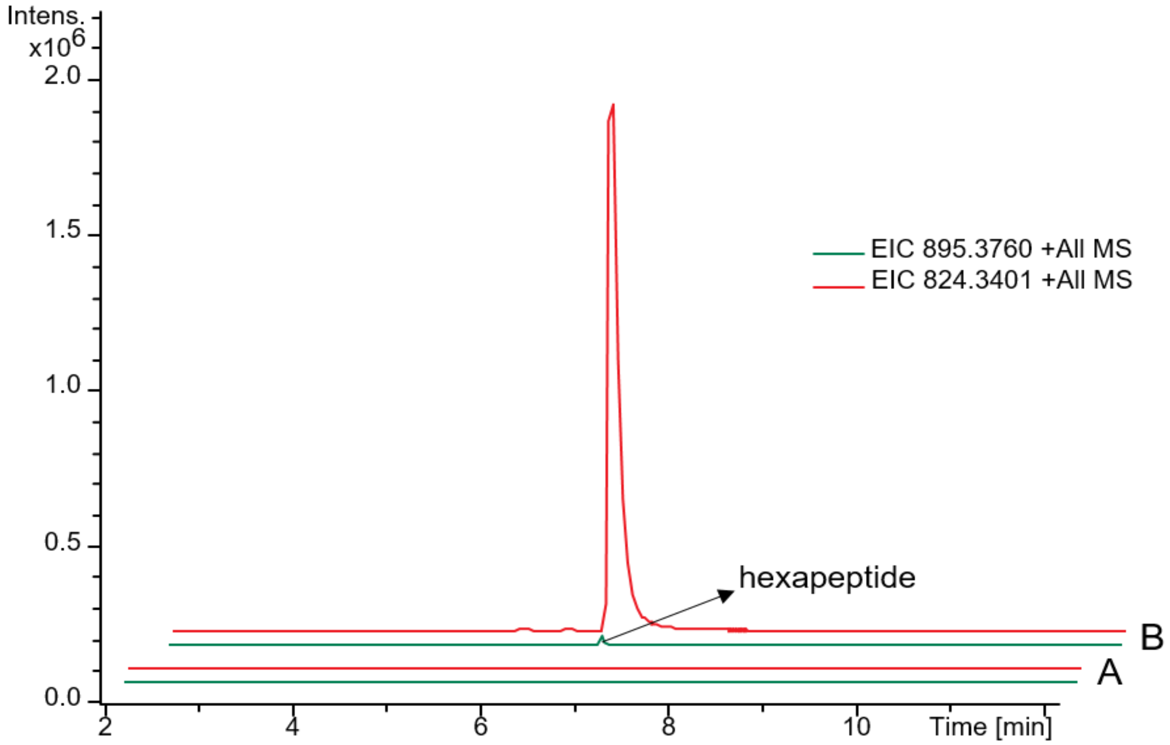


**Figure S46**. Extracted ion chromatogram of pentapeptide laurentirubin B (**6**) and hexapeptide from *S. albus* harboring pUWL201-oriT (A) and *S.albus* harboring pUWL201-OriT-*lau* (B), respectively.

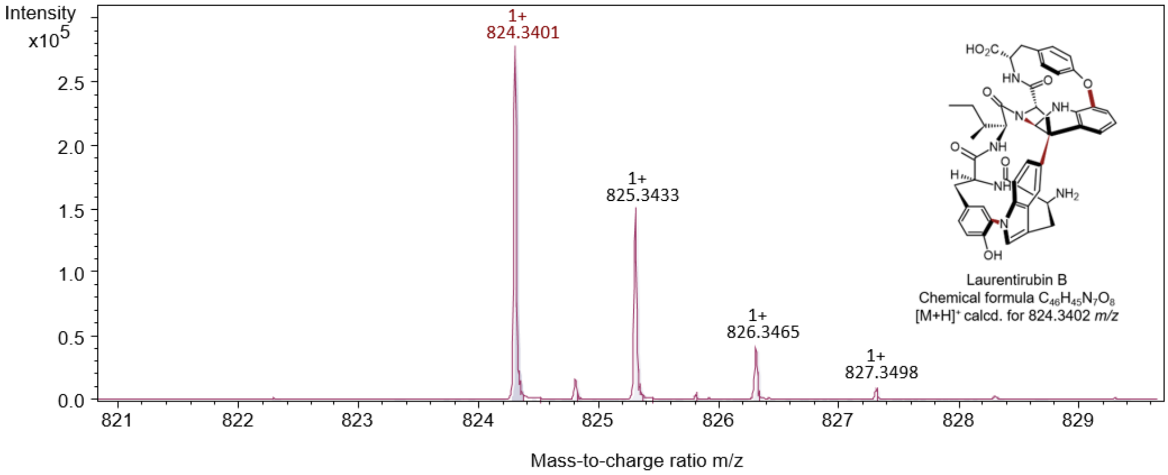


**Figure S47.** HPLC-ESI-QTOF-HRMS analysis of laurentirubin B (**6**).

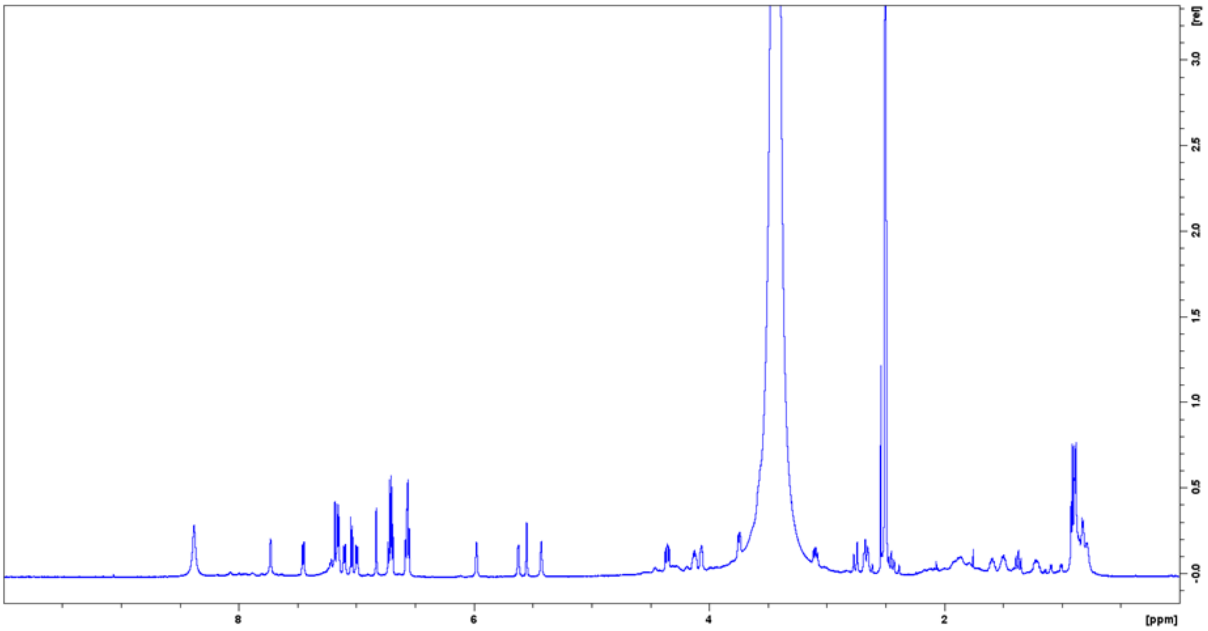


**Figure S48.** $^{1}$H NMR spectrum(600 MHz) of laurentirubin B (**6**) in DMSO-$d_6$.

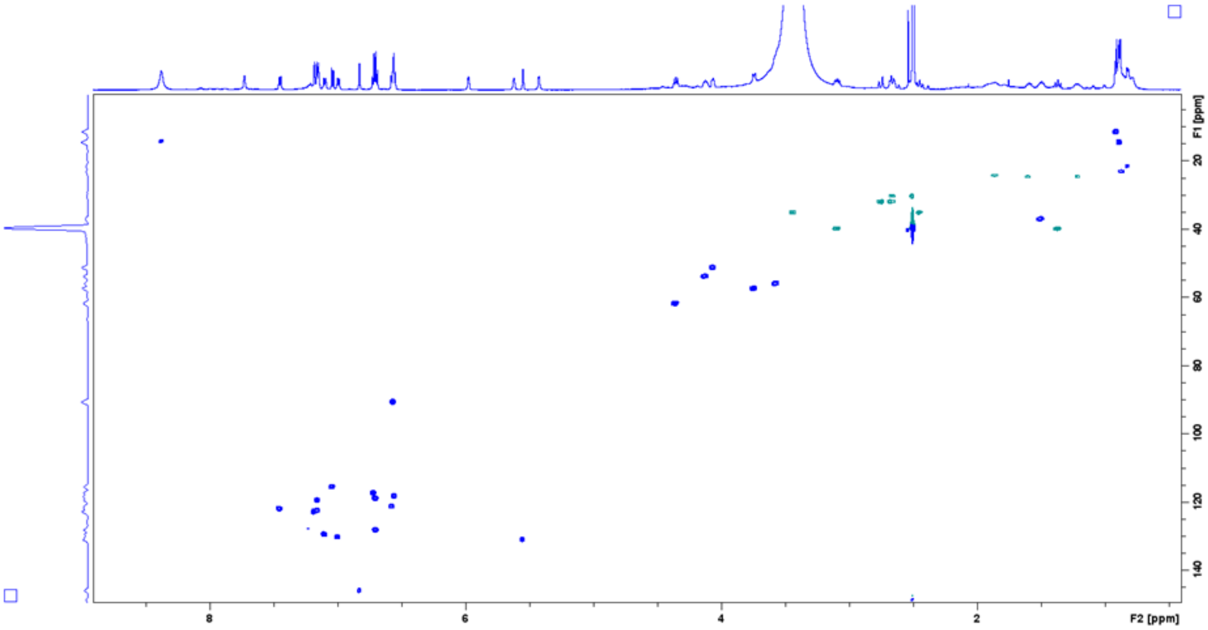


**Figure S49.** HSQC spectrum (600 MHz) of laurentirubin B (**6**) in DMSO-$d_6$.

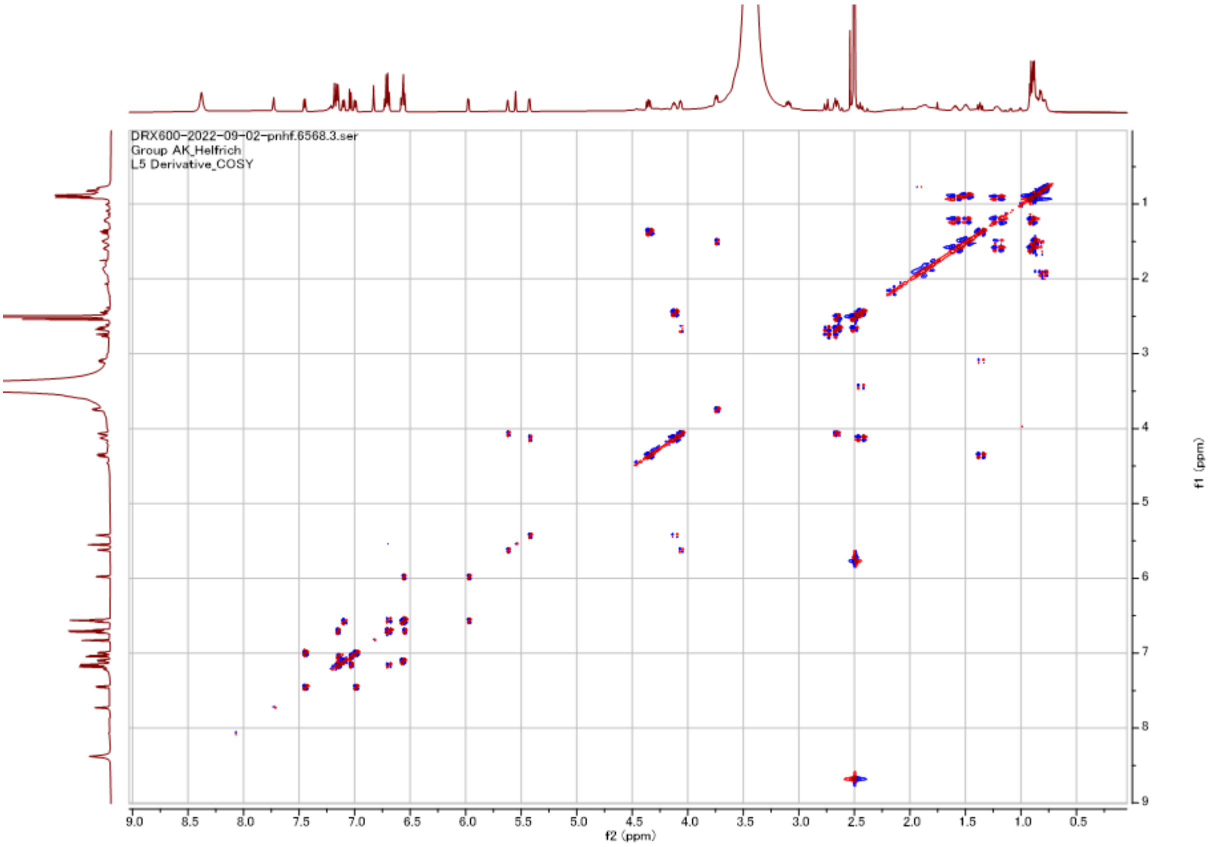


**Figure S50.** COSY spectrum (600 MHz) of laurentirubin B (**6**) in DMSO-$d_6$.

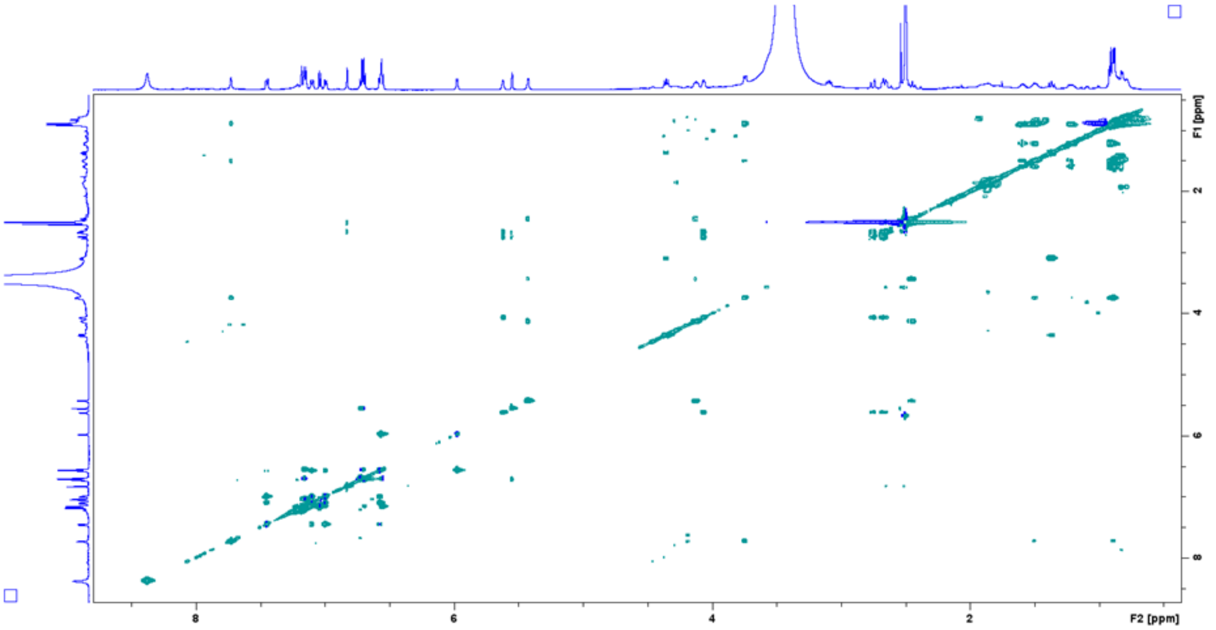


**Figure S51.** TOCSY spectrum (600 MHz) of laurentirubin B (**6**) in DMSO-$d_6$.

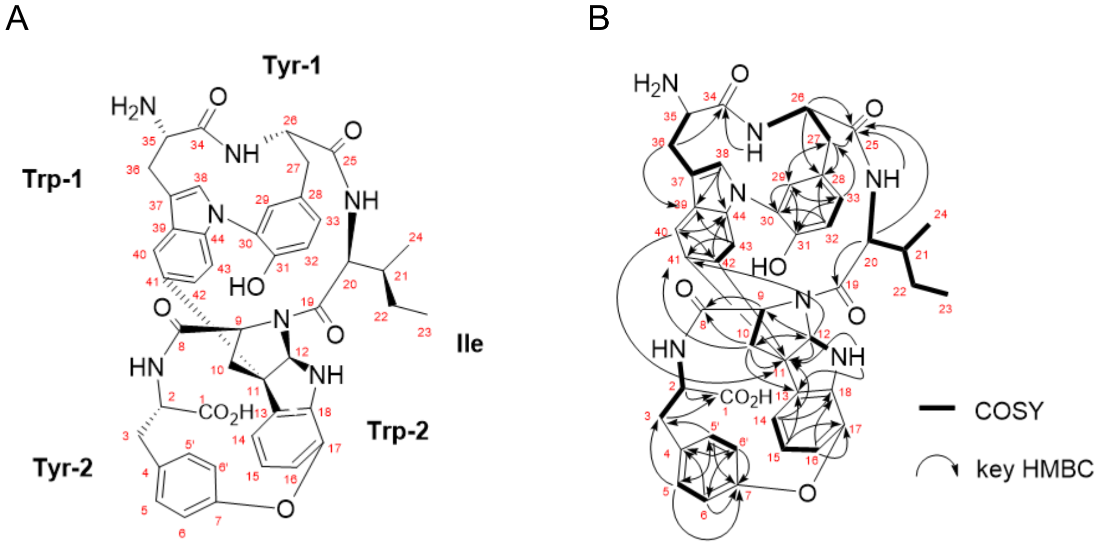


**Figure S52.** Structure of laurentirubin B (**6**). (A) Structure of laurentirubin B (**6**) and atom numbering used throughout this study. (B) COSY and key HMBC correlations of laurentirubin B (**6**) in DMSO-$d_6$.

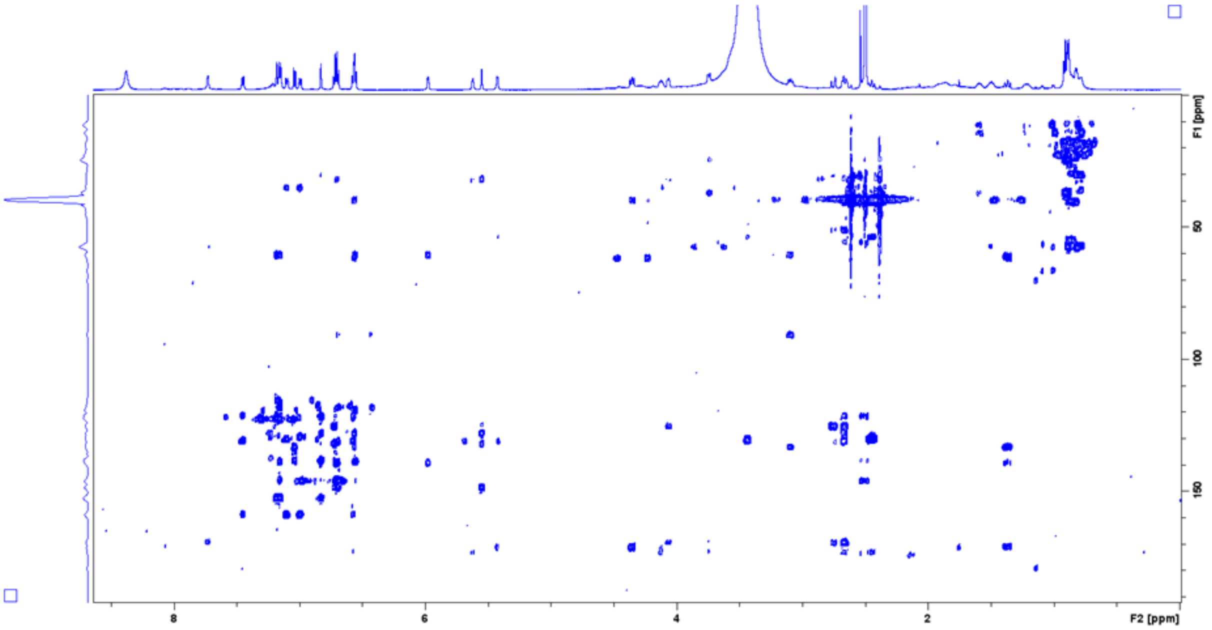


**Figure S53.** HMBC spectrum ($J$ = 8 Hz) of laurentirubin B (**6**) in DMSO-$d_6$.

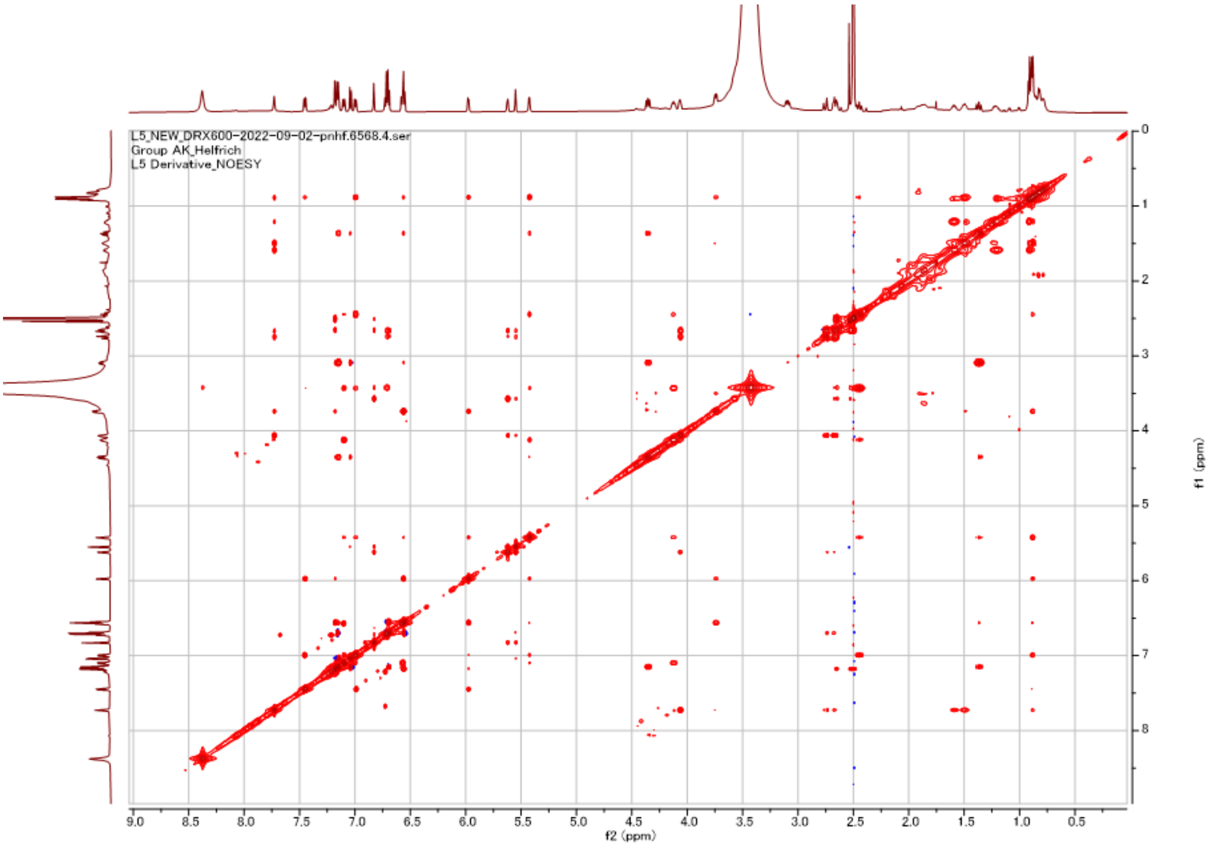


**Figure S54.** NOESY spectrum (600 MHz) of laurentirubin B (**6**) in DMSO-$d_6$.

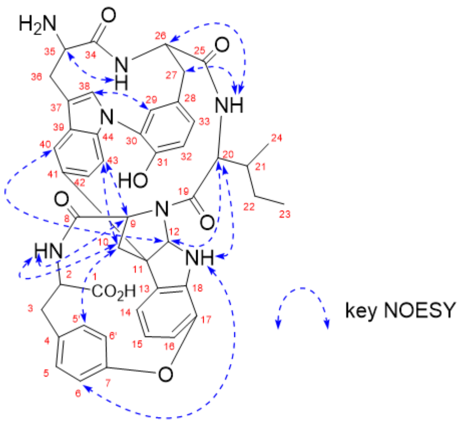


**Figure S55.** Key NOESY correlations of laurentirubin B (**6**).

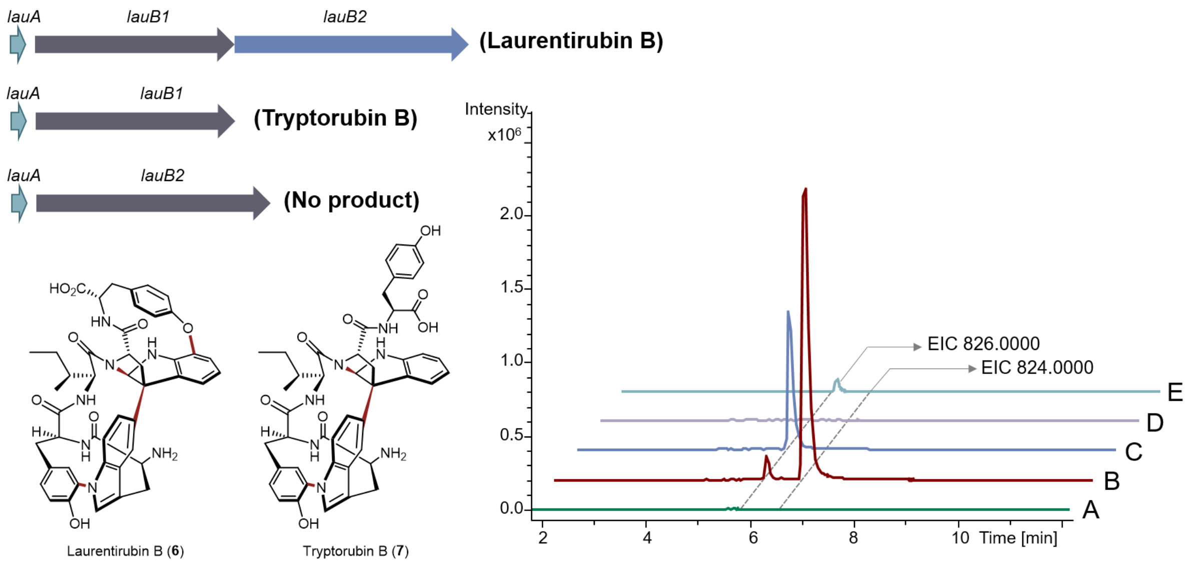


**Figure S56**. *In vivo* functional characterization of P450s in *lau* BGC. Extracted ion chromatogram of laurentirubin B (**6**) and tryptorubin B (**7**) together from different sources. (A) *S. albus* harboring pUWL201-oriT; (B) *S. albus*/pUWL201-OriT-*lau*; (C) *S. albus*/pUWL201-OriT-*lauA*+*lauB1*; (D) *S. albus*/pUWL201-OriT-*lauA*+*lauB2*; (E) authentic tryptorubin B standard.

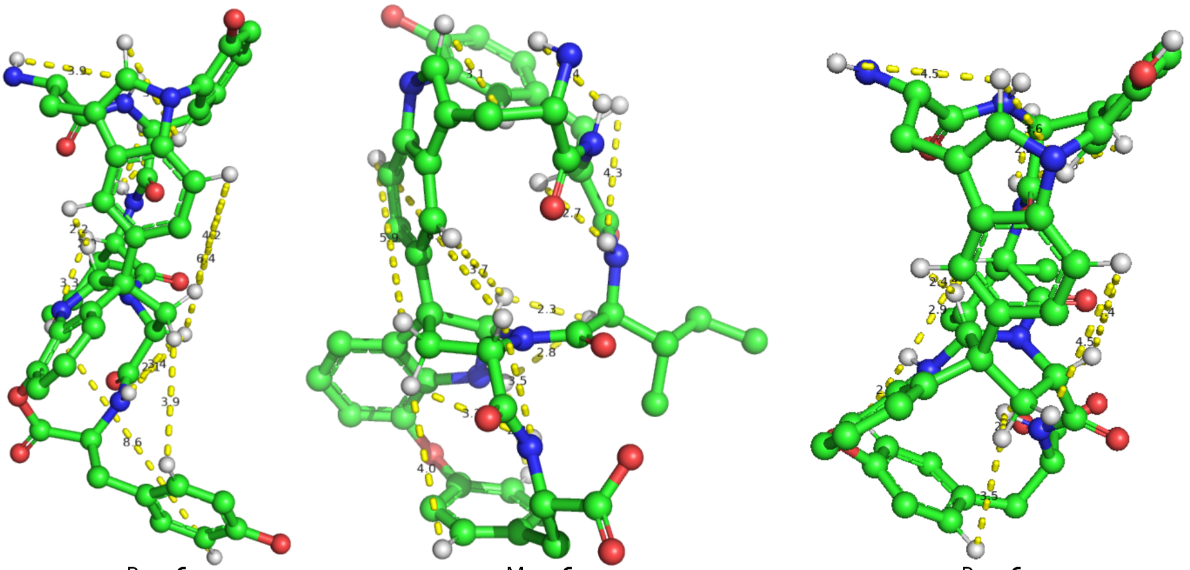

$P_{ansa}$-**6a** $M_{ansa}$-**6** $P_{ansa}$-**6**

**Figure S57.** The lowest energy conformer of $P_{ansa}$-**6a**, $M_{ansa}$-**6**, and $P_{ansa}$-**6** calculated at B3LYP/6-31G(d,p) level of theory. The values show distance (Å) of protons which showed NOESY correlations in **6**. Protons not involved in one of the NOESY correlations are hidden.

**Table S1.** Detailed metrics of final classifiers for differentiating atropopeptide-modifying P450s (class 1) from P450s modifying any other substrate (class 0)

| Classifier | Score | Balanced Accuracy Score | Cross Validation Scores | F1-Score Class 0 / 1 |
|---|---|---|---|---|
| **ExtraTreesClassifier** | 0.9997 | 0.9687 | [0.9665, 0.9614, 0.9614, 1.0000, 0.9164] | [1.00, 0.97] |
| **RandomForestClassifier** | 0.9997 | 0.9687 | [0.9665, 0.9614, 0.9164, 0.9665, 0.7994] | [1.00, 0.97] |
| **AdaBoostClassifier** | 0.9997 | 0.9687 | [0.9665, 0.9614, 0.9614, 1.0000, 0.9614] | [1.00, 0.97] |
| **BaggingClassifier** | 0.9978 | 0.9678 | [0.9114, 0.9283, 0.9614, 0.9372, 0.9164] | [1.00, 0.79] |
| **DecisionTreeClassifier** | 0.992 | 0.9649 | [0.9372, 0.9614, 0.8842, 0.9614, 0.8842] | [1.00, 0.51] |
| **MLPClassifier** | 0.9984 | 0.9681 | [0.9665, 0.9614, 0.9614, 0.9164, 0.9283] | [1.00, 0.83] |

**Table S2.** $^{1}H$ and $^{13}C$ NMR Data for varsitalin B1 (**2**) in DMSO-$d_6$.

| | $\delta_H$, mult. ($J$ in Hz) | $\delta_C$ | | $\delta_H$, mult. ($J$ in Hz) | $\delta_C$ |
|---|---|---|---|---|---|
| 1 | | 173.8 | 22[a] | | |
| 2 | 3.87, m | 52.5 | 22-$NH_2$[a] | | |
| 2-NH | 7.21, d(8.0) | | 23 | | 169.9 |
| 3 | 1.33, m | 42.3 | 24 | 3.93, m | 51.2 |
| | 1.39, m | | 24-NH | 5.89, br d | |
| 4 | 1.57, m | 24.0 | 25 | 2.61, ovlp.[iii] | 31.8 |
| 5 | 0.74, d (6.9) | 22.4 | | 2.68, dd (7.7, 17.4) | |
| 6 | 0.76, d (6.8) | 23.0 | 26 | | 125.0 |
| 7 | | 169.5 | 27 | 6.12, br s | 132.0 |
| 8 | 4.46, dd (7.3, 12.0) | 63.5 | 28 | | 131.6 |
| 9 | 2.09, m | 39.3 | 29 | | 148.2 |
| | 3.29, ovlp.[i] | | 30 | 6.69, ovlp.[ii] | 116.6 |
| 10 | | 60.0 | 31 | 6.67, ovlp.[ii] | 127.7 |
| 11 | 6.66, ovlp.[ii] | 91.6 | 32[a] | | |
| 11-NH[a] | | | 33 | 3.41, ovlp.[i] | 56.9 |
| 12 | | 135.8 | 33-$NH_2$[a] | | |
| 13 | 6.78, d (7.5) | 122.6 | 34 | 2.54, ovlp.[iv] | 30.4 |
| 14 | 6.64, t (7.5) | 117.9 | | 2.59, ovlp.[iii] | |
| 15 | 7.09, t (7.5) | 127.5 | 35 | | 121.2 |
| 16 | 6.72, d (7.5) | 108.9 | 36 | 6.72, s | 145.5 |
| 17 | | 148.6 | 37 | | 138.2 |
| 18 | | 172.4 | 38 | 7.12, s | 122.4 |
| 19 | 4.00, m | 52.4 | 39 | | 135.2 |
| 19-NH | 7.83, br s | | 40 | 7.34, d (8.4) | 122.4 |
| 20 | 1.64, m | 27.5 | 41 | 7.08, d (8.4) | 115.1 |
| | 1.90, m | | 42 | | 152.3 |
| 21 | 2.32, m | 29.7 | | | |
| | 2.36, m | | | | |

a: not observed
i: overlapped with water
ii: overlapped with H-11, H-30, and H-31
iii: overlapped with H-25 and H-34
iv: overlapped with solvent

**Table S3.** $^1H$ and $^{13}C$ NMR Data for varsitalin B2a (**4a**) in DMSO-$d_6$.

| | $\delta_H$, mult. (*J* in Hz) | $\delta_C$[a] | | $\delta_H$, mult. (*J* in Hz) | $\delta_C$[a] |
|---|---|---|---|---|---|
| 1 | | 173.6 | 22 | | 173.5 |
| 2 | 4.04, ddd (7.0, 7.0, 7.0) | 51.2 | 22-$NH_2$[b] | | |
| 2-NH | 7.26, d (7.0) | | 23 | | 170.7 |
| 3 | 1.46, m | 40.5 | 24 | 4.55, ddd (4.6, 9.1, 9.1) | 54.0 |
| 4 | 1.68, ovlp.[i] | 23.8 | 24-NH | 8.01, d (9.1) | |
| 5 | 0.88, d (6.5) | 21.9 | 25 | 2.55, dd (9.1, 13.7) | 36.3 |
| 6 | 0.90, d (6.7) | 22.7 | | 2.92, dd (4.6, 13.7) | |
| 7 | | 170.3 | 26 | | 127.7 |
| 8 | 4.32, ddd (4.2, 8.8, 8.8) | 53.4 | 27 | 6.95, d (8.3) | 130.0 |
| 8-NH | 6.95 br s | | 28 | 6.63, d (8.3) | 114.6 |
| 9 | 3.41, dd (4.2, 15.2) | 25.7 | 29 | | 155.5 |
| | 3.48, dd (8.8, 15.2) | | 30 | 6.63, d (8.3) | 114.6 |
| 10 | | 106.4 | 31 | 6.95, d (8.3) | 130.0 |
| 11 | | 136.9 | 32 | | 174.7 |
| 11-NH[a] | | | 33 | 3.30, ovlp.[ii] | 57.8 |
| 12 | | 128.8 | 33-$NH_2$[a] | | |
| 13 | 7.51, d (7.9) | 117.8 | 34 | 2.86, (dd, 3.0, 13.4) | 30.9 |
| 14 | 6.98, t (7.9) | 118.1 | | 3.03, (dd, 11.1, 13.4) | |
| 15 | 7.06, t (7.9) | 120.5 | 35 | | 110.6 |
| 16 | 7.31, d (7.9) | 110.5 | 36 | 7.24, d (1.8) | 124.9 |
| 17 | | 135.4 | 36-NH | 11.0, br s | |
| 18 | | 169.4 | 37 | | 135.9 |
| 19 | 3.68, ddd (6.8, 6.8, 6.8) | 52.1 | 38 | 7.43, s | 111 |
| 19-NH | 5.58, br s | | 39[a] | | |
| 20 | 1.31, m | 26.5 | 40 | 7.22, d (8.3) | 118.3 |
| | 1.59, m | | 41 | 7.48, d (8.3) | 119.1 |
| 21 | 1.70, ovlp.[i] | 30.6 | 42 | | 125.9 |
| | 1.83, ddd (5.9, 8.9, 14.8) | | | | |

a: assigned by cross peaks in the HSQC and HMBC spectra
b: not observed
i: overlapped with H-4 and H-21
ii: overlapped with water

**Table S4.** $^{1}$H and $^{13}$C NMR data of embyscamide (**5**) in DMSO-$d_6$

| position | $\delta_H$, mult. ($J$ in Hz) | $\delta_C$, type |
|---|---|---|
| 1 | | 173.7, C |
| 2 | 4.40, m | 52.3, CH |
| 2-NH | 8.04, d (5.7) | |
| 3 | 3.24, dd (16.0, 3.3) | 26.8, $CH_2$ |
| | 3.15, m | |
| 4 | | 111.7, C |
| 5 | 7.85, s | 128.6, CH |
| 6 | | 128.4, C |
| 7 | 7.66, d (7.9) | 118.9, CH |
| 8 | 7.12, m | 119.6, CH |
| 9 | 7.19, m | 122.1, CH |
| 10 | 7.52, d (8.0) | 109.6, CH |
| 11 | | 135.5, C |
| 12 | | 170.5, C |
| 13 | 4.06, dd (7.1, 3.8) | 57.8, CH |
| 13-NH | 7.81, d (3.8) | |
| 14 | 1.68, m | 35.8, CH |
| 15 | 1.66, m | 25.3, $CH_2$ |
| | 1.36, m | |
| 16 | 0.91, t (7.4) | 10.9, $CH_3$ |
| 17 | 0.94, d (6.8) | 15.1, $CH_3$ |
| 18 | | 170.5, C |
| 19 | 4.56, ddd (10.4, 9.3, 3.7) | 52.8, CH |
| 19-NH | 7.35, d (9.3) | |
| 20 | 2.67, dd (14.1, 3.7) | 37.4, $CH_2$ |
| | 2.57, dd (14.1, 10.4) | |
| 21 | | 138.1, C |
| 22 | 7.25, m | 129.2, CH |
| 23 | 7.21, m | 127.9, CH |
| 24 | 7.14, m | 126.1, CH |
| 23′ | 7.21, m | 127.9, CH |
| 22′ | 7.25, m | 129.2, CH |
| 25 | | 173.7, C |
| 26 | 3.35, m | 56.1, CH |
| 27 | 2.98, dd (13.2, 3.8) | 31.6, $CH_2$ |
| | 2.75, dd (12.5, 12.5) | |
| 28 | | 110.3, C |
| 29 | 6.66, d (2.2) | 124.4, CH |
| 29-NH | 10.45, s | |
| 30 | | 129.3, C |
| 31 | 7.57, d (7.85) | 117.1, CH |
| 32 | 7.15, m | 118.3, CH |
| 33 | 7.23, m | 116.2, CH |
| 34 | | 123.9, C |
| 35 | | 131.6, C |

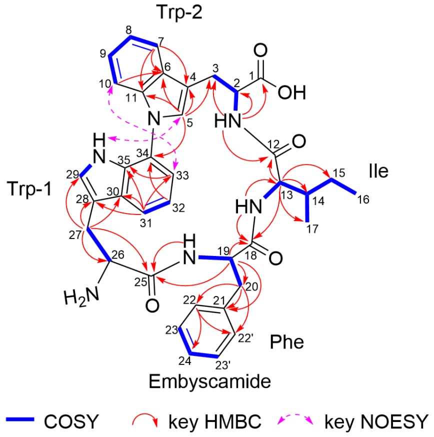

**Table S5.** $^{1}H$ and $^{13}C$ NMR data of laurentirubin B (**6**) in DMSO-$d_6$

| | $\delta_H$, mult. | $\delta_C$[a] | | $\delta_H$, mult. | $\delta_C$[a] |
|---|---|---|---|---|---|
| 1 | | 173.0 | 23 | 0.91, t (7.3) | 11.3 |
| 2 | 4.12, m | 53.8 | 24 | 0.89, d (6.6) | 14.4 |
| 2-NH | 5.43, d (3.9) | | 25 | | 169.3 |
| 3 | 2.45, t (13.1) | 35.1 | 26 | 4.07, m | 51.1 |
| | 3.44, ovlp.[i] | | 26-NH | 5.62, d (3.6) | |
| 4 | | 130.9 | 27 | 2.66, ovlp.[v] | 31.9 |
| 5 | 7.00, br d | 130.2 | | 2.76, d (17.3) | |
| 5' | 7.10, br d | 129.3 | 28 | | 125.4 |
| 6 | 7.45, dd (2.4, 8.2) | 121.9 | 29 | 5.55, s | 131.0 |
| 6' | 6.58, ovlp.[ii] | 121.2 | 30 | | 131.9 |
| 7 | | 158.9 | 31 | | 148.5 |
| 8 | | 171.2 | 31-OH | | |
| 9 | 4.36, dd (7.8, 10.9) | 61.7 | 32 | 6.72, ovlp.[iii] | 117.3 |
| 10 | 1.37, br dd | 39.8 | 33 | 6.70, ovlp.[iii] | 128.2 |
| | 3.09, dd (7.8, 13.5) | | 34 | | 173.2 |
| 11 | | 60.5 | 35 | 3.58, m | 55.8 |
| 12 | 6.57, ovlp.[ii] | 90.6 | 35-$NH_2$ | | |
| 12-NH | 5.98, d (4.5) | | 36 | 2.51, ovlp.[vi] | 30.2 |
| 13 | | 139.2 | | 2.66, ovlp.[v] | |
| 14 | 6.55, ovlp.[ii] | 118.2 | 37 | | 121.5 |
| 15 | 6.70, ovlp.[iii] | 118.9 | 38 | 6.83, s | 145.9 |
| 16 | 7.16, ovlp.[iv] | 119.4 | 39 | | 138.4 |
| 17 | | 145.7 | 40 | 7.18, s | 122.7 |
| 18 | | 138.7 | 41 | | 133.3 |
| 19 | | 172.8 | 42 | 7.16, ovlp.[iv] | 122.4 |
| 20 | 3.75, dd (2.8, 9.3) | 57.3 | 43 | 7.04, d (8.3) | 115.4 |
| 20-NH | 7.73, br s | | 44 | | 152.5 |
| 21 | 1.50, m | 36.9 | | | |
| 22 | 1.22, m | 24.6 | | | |
| | 1.59, m | | | | |

a: assigned by cross peaks in the HSQC and HMBC spectra
i: overlapped with water signal
ii: overlapped with H-6', H-12, and H-14
iii: overlapped with H-15, H-32, and H-33
iv: overlapped with H-16 and H-42
v: overlapped with H-27 and H-36
vi: overlapped with solvent

**Table S6.** Comparison of experimental and calculated $^{13}$C NMR chemical shifts for **6**

| no. | exp. $\delta_C$ | $P_{ansa}$-**6** | $P_{ansa}$-**6a**[a] | $M_{ansa}$-**6** |
|---|---|---|---|---|
| 1 | 173.0 | 175.8 | 167.6 | 169.3 |
| 2 | 53.8 | 49.6 | 67.7 | 55.5 |
| 3 | 35.1 | 35.2 | 45.4 | 37.1 |
| 4 | 130.9 | 132.3 | 129.3 | 133.0 |
| 5 | 130.2 | 133.8 | 130.4 | 130.8 |
| 6 | 129.3 | 128.9 | 129.4 | 130.5 |
| 7 | 121.9 | 120.5 | 115.6 | 122.0 |
| 8 | 121.2 | 120.5 | 113.7 | 120.5 |
| 9 | 158.9 | 163.4 | 154.7 | 161.7 |
| 10 | 171.2 | 168.4 | 168.9 | 168.2 |
| 11 | 61.7 | 67.0 | 64.6 | 67.9 |
| 12 | 39.8 | 39.8 | 49.9 | 45.2 |
| 13 | 60.5 | 67.4 | 76.6 | 68.7 |
| 14 | 90.6 | 95.9 | 91.3 | 91.8 |
| 15 | 139.2 | 140.4 | 138.3 | 139.4 |
| 16 | 118.2 | 119.4 | 120.7 | 119.6 |
| 17 | 118.9 | 119.1 | 117.9 | 119.7 |
| 18 | 119.4 | 121.9 | 121.5 | 122.5 |
| 19 | 145.7 | 148.1 | 135.8 | 147.5 |
| 20 | 138.7 | 140.4 | 146.0 | 139.3 |
| 21 | 172.8 | 169.0 | 166.7 | 174.2 |
| 22 | 57.3 | 60.7 | 64.6 | 62.2 |
| 23 | 36.9 | 42.6 | 49.2 | 40.5 |
| 24 | 24.6 | 29.6 | 33.0 | 26.8 |
| 25 | 11.3 | 12.0 | 19.2 | 11.7 |
| 26 | 14.4 | 11.1 | 19.3 | 12.4 |
| 27 | 169.3 | 174.1 | 174.0 | 169.8 |
| 28 | 51.1 | 55.9 | 61.6 | 65.5 |
| 29 | 31.9 | 34.1 | 40.8 | 29.4 |
| 30 | 125.4 | 126.0 | 127.0 | 129.7 |
| 31 | 131.0 | 136.1 | 135.2 | 125.5 |
| 32 | 131.9 | 135.4 | 136.1 | 130.3 |
| 33 | 148.5 | 149.2 | 148.6 | 150.2 |
| 34 | 117.3 | 114.2 | 115.5 | 116.7 |
| 35 | 128.2 | 128.0 | 128.5 | 128.5 |
| 36 | 173.2 | 173.9 | 175.7 | 180.5 |
| 37 | 55.8 | 64.1 | 68.1 | 58.6 |
| 38 | 30.2 | 35.6 | 36.9 | 32.3 |
| 39 | 121.5 | 122.1 | 124.9 | 130.2 |
| 40 | 145.9 | 145.7 | 146.5 | 144.9 |
| 41 | 138.4 | 140.2 | 140.1 | 138.7 |
| 42 | 122.7 | 122.8 | 122.1 | 124.0 |
| 43 | 133.3 | 136.8 | 140.4 | 140.0 |
| 44 | 122.4 | 123.3 | 123.9 | 127.6 |
| 45 | 115.4 | 116.0 | 117.4 | 116.8 |
| 46 | 152.5 | 153.1 | 151.6 | 160.8 |
| MAE[b] | | 2.57 | 4.96 | 3.00 |
| MSE[c] | | 10.97 | 41.57 | 17.38 |

[a]Compound **6a** has aryl ester bridge between C-1 and C-17 instead of aryl ether bond of C-7 and C-17. [b]Mean absolute errors (ppm). [c]Mean squared error.

**Table S7**. Strains and plasmids used and constructed in this study

| Strains/plasmids | Characteristic(s) | Sources |
|---|---|---|
| ***E. coli*** | | |
| DH5a | Host strain for cloning | [1] |
| ET12567/pUZ8002 | Donor strain for conjugation, Chl[r], Kan[r] | [2] |
| **Actinobacteria** | | |
| *Streptomyces jumonjinensis* DSM 747 | Wild type strain containing *jum* BGC | DSMZ[a] |
| *Streptomyces varsoviensis* DSM 40346 | Wildtype strain containing *sva* BGC | DSMZ[a] |
| *Embleya scabrispora* DSM 41855 | Wild type strain containing *emb* BGC | BCCM[b] |
| *Streptomyces laurentii* DSM 41684 | Wildtype strain containing *lau* BGC | DSMZ[a] |
| *S. albus* J1074 | Heterologous host | [3] |
| *S. albus* J1074/pUWL201-OriT | The introduction of pUWL201-OriT into *S. albus* J1074 | This study |
| *S. albus* J1074/pUWL201-OriT-*jum* | The introduction of pUWL201-OriT-*jum* into *S. albus* J1074 | This study |
| *S. albus* J1074/pUWL201-OriT-Svarcore1 | The introduction of pUWL201-OriT-Svarcore1 into *S. albus* J1074 | This study |
| *S. albus* J1074/pUWL201-OriT-Svarcore2 | The introduction of pUWL201-OriT-Svarcore2 into *S. albus* J1074 | This study |
| *S. albus* J1074/pUWL201-OriT-Svarcore1R | The introduction of pUWL201-OriT-Svarcore1R into *S. albus* J1074 | This study |
| *S. albus* J1074/pUWL201-OriT-Svarcore2R | The introduction of pUWL201-OriT-Svarcore2R into *S. albus* J1074 | This study |
| *S. albus* J1074/pUWL201-OriT-*emb* | The introduction of pUWL201-OriT-*erm* into *S. albus* J1074 | This study |
| *S. albus* J1074/ pUWL201-OriT-*lau* | The introduction of pUWL201-OriT-*lau* into *S. albus* J1074 | This study |
| *S.albus* J1074/ pUWL201-OriT-*lauA*+*lauB1* | The introduction of pUWL201-OriT-*lauA*+*lauB1* into *S.albus* J1074 | This study |
| *S.albus* J1074/ pUWL201-OriT-*lauA*+*lauB2* | The introduction of pUWL201-OriT-*lauA*+*lauB2* into *S.albus* J1074 | This study |
| **Plasmids** | | |
| pUWL201-OriT | Apr[r], *ermE**p, replicative expression vector in *Streptomyces* | [4] |
| pUWL201-OriT-*jum* | Apr[r], *jum* BGC was constructed into pUWL201-OriT | This study |
| pUWL201-OriT-Svarcore1 | Apr[r], *sva* BGC containing *svaA2* and *svaB* was constructed into pUWL201-OriT | This study |
| pUWL201-OriT-Svarcore2 | Apr[r], *sva* BGC containing *svaA1, svaA2* and *svaB* was constructed into pUWL201-OriT | This study |
| pUWL201-OriT-Svarcore1R | Apr[r], a RBS was inserted in front of *svaB* based on plasmid pUWL201-OriT-Svarcore1 | This study |
| pUWL201-OriT-Svarcore2R | Apr[r], a RBS was inserted in front of *svaB* based on plasmid pUWL201-OriT-Svarcore2 | This study |
| pUWL201-OriT-*emb* | Apr[r], *emb* BGC was constructed into pUWL201-OriT | This study |
| pIJ10257 | Hyg[r], *ermE**p, integrative expression vector in *Streptomyces* | [5] |

| | | |
|---|---|---|
| pIJ10257-*lau* | Hyg$^r$, *lau* BGC was constructed into pIJ10257 | This study |
| pUWL201-OriT-*lau* | Apr$^r$, *lau* BGC was constructed into pUWL201-OriT | This study |
| pUWL201-OriT-*lauA*+*lauB1* | Apr$^r$, *lauA and lauB1* was constructed into pUWL201-OriT | This study |
| pUWL201-OriT-*lauA*+*lauB2* | Apr$^r$, *lauA and lauB1* was constructed into pUWL201-OriT | This study |

DSMZ[a]: Deutsche Sammlung von Mikroorganismen und Zellkulturen.

BCCM[b]: Belgian Coordinated Collections of Microorganisms.

**Table S8**. Primers used in this study

| **Primers** | **Sequences (5' to 3')** |
|---|---|
| **For amplifying *jum* BGC** | |
| jumo_fwd | gaggcttgatATGAAGGTTCTCTTTGCCATTCGGCACAAGGTCAC |
| jumo_rev | ggaattcgatTCAGCGGACCCGGGCCGC |
| **For linearizing pUWL201-oriT to insert *jum* BGC** | |
| jumo_bb_fwd | ggtccgctgaATCGAATTCCTGCAGCCC |
| jumo_bb_rev | gaaccttcatATCAAGCCTCCTGTTCTAG |
| **For amplyifing *sva* BGC** | |
| Svarcore1_fwd | TCTAGAACAGGAGGCCCCATATGGAGGAATTTATGAAGCTGGTTCACCTG |
| Svarcore1_2_rev | TCGATATCAAGCTTATCGATTCACGGGCGCACCATCCG |
| Svarcore2_fwd | TCTAGAACAGGAGGCCCCATATGTATTCGGAAGGGGTTCCGGTTC |
| **For linearizing pUWL201-oriT to insert *sva* BGC** | |
| pUWL_OriT_fwd | ATCGATAAGCTTGATATCGAATTCCTGCAGC |
| pUWL_RBS_rev | ATGGGGCCTCCTGTTCTAGACGATC |
| **For inserting RBS into pUWL201-OriT (*sva* cloning)** | |
| Svar_RBS_CYP450_Fwd | AGGAGGTCACCCATGCCAATGCATCGC |
| Svar_CYP450_Rev | ACTCCGTTGCTGGGCTGCCC |
| **For amplifying *emb* BGC** | |
| emb-pp-F | GTCTAGAACAGGAGGCCCCATGTGATCAAGATCGTCAACTC |
| emb-pp-R | CAGGAATTCGATATCAAGCTTTCACCTCCCCGAGGCGAGG |
| **For linearizing the pUWL201-oriT to insert *emb* BGC** | |
| pUWL201-OriT-F | aagcttgatatcgaattcc |
| pUWL201-OriT-R | atggggcctcctgttctag |
| **For amplifying *lau* BGC** | |
| TrypLaurentii_Fwd_NdeI | CGCCATATGATGAAGCTTCTCTTCGCCATTCGC |
| TrypLaurentii_RV_XhoI | CCGCTCGAGTCAGTGTAGGCGGACCGGGAGG |
| **For reversely amplifying plasmid containing *lauA* and *lauB1*** | |
| lauAB1-F | ATCGATaagcttgatatcgaattcctgcag |
| lauAB1-R | TCAGACAGTGGCGGCACGCC |
| **For amplifying lauA fragment** | |
| lauA-F | gtctagaacaggaggccccatATGATGAAGCTTCTCTTCGC |
| lauA-R | GACAGTGGCGGCACGGCAAGTCTCCGGGAGGCCGAG |
| **For amplifying lauB2 fragment** | |
| lauB2-F | GCCTCCCGGAGACTTGCCGTGCCGCCACTGTCTGAAC |

| | |
|---|---|
| lauB2-R | ttcgatatcaagcttATCGATTCAGTGTAGGCGGACCGGGAG |
| **For linearizing pUWL201-OriT to coexpress *lauA* and *lauB2*** | |
| pUWL-LF | ATCGATaagcttgatatcg |
| pUWL-LR | atggggcctcctgttctag |

### Methods

#### Generation of the Training data set

For the development of the machine learning (ML) classifier, a negative dataset was assembled from the antiSMASH database[6], comprising 14,665 proteins annotated as 'cytochrome p450.' Concurrently, a positive dataset was meticulously curated by deploying the tryptorubin A cytochrome P450 WP_007820080.1 as a query for Blastp analysis against the NCBI non-redundant protein sequences (nr) database. From this analysis, 51 putative biosynthetic gene cluster (BGC) sequences were identified through a manual search for precursors situated in the genetic neighborhood of each Blast hit. To dereplicate both datasets, cdhit V4.8.1[7] was employed. Parameters were configured to cluster sequences with a minimum of 95% sequence similarity and to utilize a word size of 5. As a result of this dereplication process, the negative and positive datasets were refined to 9,065 and 37 sequences, respectively.

#### Assembly of classification dataset

For the classification, a dataset comprising 154,364 protein sequences was derived from the NCBI Identical Protein Groups database (25.01.2023).[8] These sequences, which encompassed the term "p450" in their descriptions and which had lengths ranging from 300 to 450 amino acids, were procured using the following query parameters: p450[All Fields] AND (refseq[filter] AND prokaryotes[filter] AND ("300"[SLEN] : "450"[SLEN])).[8] The method employed to assemble the dataset mirrored that of the training dataset."

#### Data preprocessing

Sequences were aligned using Clustal W version 1.2.3[9] with the default settings (clustalo-1.2.3-Ubuntu-x86_64 -i input.fasta -o clustal.aln -v --outfmt=clustal --output-order=tree-order --auto -t Protein) with a reference cytochrome P450 with known annotation of functional regions from *Mycobacterium tuberculosis* H2102 (GenBank: KBE51585.1) in a multiple sequence alignment. The sequences were fragmented at positions 92, 192, 275, and 395 of the reference to resemble the different functional regions of the cytochrome obtained from the annotations in the Cytochrome P450 Engineering Database record 10800 (N-terminus, substrate binding region 1, substrate binding region 2, core region and C-terminus).[10]
To prevent overfitting, the sequences were translated into a simplified amino acid code (Figure S3). The dataset was balanced using the Random Over Sampler from the Python package Imbalanced-learn 0.9.0.[11]

#### Training of the classifier and hyperparameter optimization

The machine learning model was implemented using scikit-learn version 1.0.2[12] for python. The number of overlapping *k*-mers of motifs with the length of four that occurred in at least half of the 37 putative atropopeptide sequences in each segment were used as features. Different classifiers were compared and the Random Forest classifier was chosen because of its high f1 score for atropopeptide P450s (0.97). Hyperparameter tuning was performed to optimize the maximum amount of samples per leaf and the maximum tree depth assessed on the obtained maximum balanced accuracy (balanced accuracy = (recall+ specificity)/2) (optimal parameters: 1 and 10, respectively). Every hit with a score > 0.15 was considered "positive". After the first run on the classification dataset, 202 putative atropopeptide cytochrome p450s were curated from the results using CoreFinder and sequence alignments. These additional putative atropopeptide cytochrome P450s were chosen to supplement the training dataset, leading to a size of 113 for the positive training data set after deduplication with cd-hit V4.8.1[7] (cutoff 0.95, word size = 5). The classifiers underwent training on the newly acquired dataset, according to the previous methodology (f1 score = 0.97). Parameter optimization yielded optimal values of a singular leaf (maximum leaves = 1) and a tree depth constrained to 14 (maximum tree depth = 14).
To obtain the variances of the classification for each point in the dataset, forestci version 0.6[13] was used.

#### Corefinder

Potential genes encoding precursor and core peptides were investigated in the 3 kb genetic neighborhood, both upstream and downstream of the putative atropopeptide cytochrome P450 genes in all relevant nucleotide records. Open reading frames (ORFs) ranging between 10 to 40 amino acids that contained either two consecutive aromatic amino acids within their six amino acid core peptide (first round) or one of the motifs 'KSLK', 'RSLK', 'ESLK', 'KSRK', 'KPLK', or 'PSLK' in the precursor peptide (second round) were selected. ORFs that overlapped with annotated coding sequences (CDS), with the exception of those labeled as "tryptorubin family RiPP precursor CDS" were ignored. Overlapping cores were sorted out to remove duplicates with slightly longer putative leader peptide encoding genes. It is noteworthy, that the exact start and thus length of the precursor peptides cannot be determined by Corefinder if multiple putative start codons are present. Corefinder uses fasta files containing protein identifiers and outputs a genbank file containing the region 3kb upstream and downstream with annotated precursor peptides.

#### Comparison with other tools

To compare AtropoFinder to state-of-the-art genome mining tools, all Genbank files of putative atropopeptide regions from the AtropoFinder output were converted to fasta files and merged to create a record that could then be analyzed by the respective tool. We used GECCO version 0.9.6[14], DeepBGC version 0.1.23[15], the antiSMASH version 7 webserver[16] (strictness: loose), the PRISM 4 webserver[17] and RODEO 2[18] all on default settings. The plot in Figure S5 was created using BGCViz[19].

#### Sequence Logos

The sequence logos were created using WebLogo 3[20] from sequence alignments of core peptides and precursor peptides using Clustal W version 1.2.3[9] with the default settings.

#### Sequence Similarity Networks

A sequence similarity network was generated utilizing the EFI-EST[21] platform from all putative atropopeptide precursors that contain the “KSLK” or a related motif. For both the precursors and P450s, an e-value of 5 and a threshold of 8 were set.

The generated sequence similarity network was then visualized using Cytoscape 3.10.1[22], and annotations were added using the AutoAnnotate 1.4 plugin[23].

#### Phylogenetic analysis of atropopeptide-modifying P450s

To build a maximum likelihood tree of all atropopeptide cytochrome p450s, all P450s detected by Atropofinder that had a Corefinder hit were aligned using ClustalW version 1.2.3[9] with the default settings. The alignment was trimmed using ClipKIT version 1.4.0[24] using the default settings. The tree was assembled using IQ-TREE multicore version 1.6.12[25] with default settings and visualized using iTOL version 5[26]. To perform the co-occurrence analysis, the RODEO2 web tool[18] was used with all putative P450s determined by AtropoFinder as input. To compare the GC content, the GC content of the P450 was compared to the average GC content of the species queried from the NCBI taxonomy browser[27].

#### BiG - SCAPE

The gene cluster family analysis was conducted using BiG-SCAPE[28]. The tool was run from the command line using the following parameters: ~/bin/run_bigscape . Output_bigscape/ --include_gbk_str * --include_singletons --mix --cutoffs 0.1 0.25 0.5 0.7 0.8 1.0 --mibig -v.

#### Protein structure modeling

Protein structure modeling was performed using AlphaFold 2[29] in multimer mode with standard parameters. The structures were visualized in pymol[30].

#### Materials

All chemicals, reagents and solvents were purchased from Sigma-Aldrich or Carl Roth (Germany), unless stated otherwise. Antibiotics and medium components were purchased from Carl Roth (Germany). Oligonucleotide primers were synthesized by Microsynth AG (Germany). All restriction enzymes, Q5 High Fidelity DNA polymerase, HiFi DNA Assembly master mix, T4 DNA ligase, deoxynucleotides (dNTPs) and DNA Ladder were purchased from New England Biolabs (UK). Monarch™ DNA gel extraction kit, Monarch™ Genomic DNA Purification Kit and Monarch™ Plasmid Miniprep kit from New England Biolabs (UK) were used to purify DNA.

#### Strains, plasmids and culture conditions

Bacterial strains and plasmids used in this study are summarized in Table S7. *Escherichia coli* strains were cultured in Luria-Bertani (LB) broth or on LB agar (10 g/L tryptone, 5 g/L yeast extract, 10 g/L NaCl, 15 g/L agar)) overnight at 37 °C. Tryptic soy broth medium (TSB) (BD Difco, UK) was used to grow *Streptomyces* strains for routine applications and Mannitol-soy flour agar (MS agar) (20 g/L mannitol, 20 g/L soya flour, 20 g/L agar, and 10 mM $MgCl_2$) or ISP2 (4.0 g/L glucose, 4.0 g/L yeast extract and 10.0 g/L malt extract and 20 g/L agar) was used for sporulation. *Streptomyces* strains were grown at 30 °C for 4-8 days. For the cultivation of plasmid containing clones, the medium was supplemented with the appropriate antibiotics at the following final concentrations: 50 µg/mL kanamycin and 50 µg/mL apramycin.

#### Molecular cloning

All polymerase chain reactions (PCRs) were conducted on a C1000 Touch Thermal Cycler (Bio-Rad) using Q5 High Fidelity DNA polymerase according to the manufacturer's instructions. PCR reactions were performed using the following conditions: initial denaturation (10 min, 98 °C) followed by 30-35 cycles of denaturation (20 s, 98 °C), annealing (30 s, 53–72 °C, depending on the melting temperature of primers) and elongation (based on PCR product length 30 s/1 kb, 72 °C); and final extension (10 min, 72 °C). DNA fragments from restriction digestions or PCR reactions were purified by agarose gel electrophoresis and isolated using the kits mentioned above. All plasmids constructed were verified by Sanger sequencing.

*jum* BGC

A 1252 bp DNA fragment containing the jumorubin BGC from *Streptomyces jumonjinensis* DSM 747 was PCR amplified with the primer pair (jumo_fwd and jumo_rev)(Table S8) using extracted genomic DNA as a template. The plasmid backbone was PCR amplified with the primer pair (bb_jumo_fwd and bb_jumo_rev)(Table S8) using pUWL201-oriT as a template.

The two resulting DNA fragments were assembled using HiFi assembly to generate pUWL201-OriT-*jum*.

*sva* BGC

The *sva* BGC from *Streptomyces varsoviensis* DSM 40346 contains two putative precursor genes *svaA1* and *svaA2*.

A 1219 bp DNA fragment SvarCore1 containing the *sva* BGC with the second precursor gene *svaA2* and P450 gene *svaB* from *Streptomyces varsoviensis* DSM 40346 was amplified from *S. varsoviensis* DSM 40346 genomic DNA using primer pairs Svarcore1_fwd and Svarcore1_2_rev (Table S8). The plasmid backbone (pUWL201-OriT) was linearized by PCR using primer pairs pUWL-OriT_fwd and pUWL_RBS_Rev (Table S8). The two resulting DNA fragments were assembled using HiFi assembly to generate pUWL201-OriT-SvarCore1.

A 1324 bp DNA fragment SvarCore2 containing the *sva* BGC with both precursor genes *svaA1*, *svaA2* and P450 gene *svaB* from *S. varsoviensis* DSM 40346 was amplified from the genomic DNA of *S. varsoviensis* DSM 40346 using the primer pairs Svarcore2_fwd and Svarcore1_2_rev (Table S8). The plasmid backbone (pUWL201-OriT) was linearized by PCR using the primer pairs pUWL-OriT_fwd and pUWL_RBS_Rev (Table S8). The two resulting DNA fragments were assembled using HiFi assembly to generate pUWL201-OriT-SvarCore2.

To insert a ribosomal binding site in front of *svaB*, the plasmids pUWL201-OriT-SvarCore1 and pUWL201-OriT-SvarCore2 were used as template for reverse amplification by PCR using primer pairs Svar_RBS_CYP450_Fw and Svar-CYP450_Rev (Table S8) to afford the linearized plasmid, which was then deal with KLD mix. 1 µL of the KLD mix reaction was transformed into *E. coli* DH5α. The generated plasmids were named as pUWL201-OriT-SvarCore1R and pUWL201-OriT-SvarCore2R, respectively.

*emb* BGC

A 1364 bp DNA fragment containing the *emb* BGC from the *Embleya scabrispora* DSM 41855 was PCR amplified using primer pairs emb-pp-F/R (Table S8) with the genomic DNA as template. The pUWL201-OriT backbone was linearized by PCR using primer pairs pUWL201-OriT-F/R (Table S8). The two resulting DNA fragments were assembled using HiFi assembly to generate pUWL201-OriT-*emb* after sequencing confirmation.

*lau* BGC

A 2436 bp DNA fragment containing the *lau* BGC from *S. laurentii* ATCC 31255 was first cloned into pIJ10257. The *lau* BGC fragment was amplified from genomic DNA with primer pairs TrypLaurentii_Fwd_NdeI and TrypLaurentii_RV_XhoI (Table S8). The plasmid pIJ10257 was linearized with *Nde*I and *Xho*I. The linearized pIJ10257 and the amplified *lau* BGC fragment were ligated using T4 DNA ligase to generate pIJ10257-*lau* after sequencing confirmation.
To clone *lau* BGC into pUWL201-OriT, the plasmid pUWL201-OriT was linearized with *Kpn*I and *Bam*HI. The fragment containing *lau* BGC was excised from pIJ10257-*lau* using *Kpn*I

and *Bam*HI. The linearized pUWL201-OriT and *lau* BGC fragments were ligated using T4 DNA ligase to generate pUWL201-OriT-*lau* after sequencing confirmation.

Coexpression of *lauA* and *lauB1*

To coexpress the *lauA* and *lauB1* genes, the plasmid pUWL201-OriT-*lau* was amplified using prime pairs lauAB1-F/R (Table S8) to afford the linearized plasmid, which was then treated with KLD mix. 1 µL of the KLD mix reaction was transformed into *E. coli* DH5α. The generated plasmid was named as pUWL201-OriT-lauA+lauB1.

Coexpression of *lauA* and *lauB2*

To coexpress the *lauA* and *lauB2* genes, the fragment containing *lauA* and fragment containing *lauB2* were PCR amplified by primer pairs lauA-F/R and lauB-F/R (Table S8), respectively, using plasmid pUWL201-OriT-*lau* as template. The pUWL201-OriT backbone was linearized by PCR using primer pairs pUWL-LF/LR (Table S8). The three resulting DNA fragments were assembled using HiFi assembly to generate plasmid pUWL201-OriT-lauA+lauB2.

#### Heterologous expression of selected BGCs

For heterologous expression of *jum* BGC, the plasmid pUWL201-OriT-*jum* were transformed into the heterologous host *S. albus* J1074 by conjugation. Briefly, the plasmid pUWL201-OriT-*jum* was transformed into *E. coli* ET12567/pUZ8002 to afford ET12567/pUZ8002/pUWL201-OriT-*jum* as the donor strain. 20 mL of LB medium was prepared to culture the donor strain at 37 °C, 180 rpm with appropriate antibiotics to an $OD_{600}$ of 0.6-0.8. The cells were harvested, washed twice with 20 mL antibiotic-free LB broth and resuspended in 400 uL of LB medium. The *S.albus* J1074 was streaked on MS agar plate and cultivated at 30 ºC for 5-7 days for sporulation. Then the spores of *S. albus* J1074 (recipient strain) were harvested, resuspended in 600 µL TSB medium and heated at 50 ºC for 10 min. After heating, the spore suspensions were incubated at 30 ºC for 0.5-1 h. The donor strain and the recipient strain were mixed and then diluted 1000 times. After dilution, 200 µL of the mixed strains were spread onto MS agar plates containing 10 mM $MgCl_2$. After incubation at 30 ºC for 15-18 h, the plates were overlayed with apramycin (50 µg/mL) and trimethoprim (100 µg/mL) or apramycin (50 µg/mL) and nalidixic acid (50 µg/mL) solutions and incubated at 30ºC for 3-4 days or until exconjugants were visible. The exconjugants were individually picked and re-streaked onto MS agar plate or ISP2 agar plates containing apramycin (50 µg/mL) and trimethoprim (100 µg/mL) or apramycin (50 µg/mL) and nalidixic acid (50 µg/mL) and incubated at 30 ºC for growth and sporulation. Three positive clones were randomly selected for small scale fermentation and subjected to metabolite analysis by LC-MS.

For heterologous expression of *sva* (SvarCore1, SvarCore2, SvarCore1R and SvarCore2R), *emb*, *lau*, *lauA*+*lauB1* and *lauA*+*lauB2*, the corresponding plasmids pUWL201-OriT-SvarCore1, pUWL201-OriT-SvarCore2*,* pUWL201-OriT-SvarCore1R, pUWL201-OriT-SvarCore2R, pUWL201-OriT-*emb,* pUWL201-OriT-*lau,* *pUWL201-OriT-lauA+lauB1,* pUWL201-OriT-*lauA+lauB2* were transformed into heterologous host *S. albus* J1074 by conjugation as described above.

#### Small-scale fermentation and LC-MS analysis

For the small scale fermentation, the heterologous recombinant strains were streaked on MS agar plate with apramycin (50 µg/mL) and cultivated at 30 ºC for 5-7 days for sporulation. Then an aggregate of spores of the heterologous recombinant strain was collected and transferred into a 250 mL Erlenmeyer flask containing 50 mL of TSB medium with apramycin (50 µg/mL) and incubated at 30 ºC, 180 rpm for 6 days. On the fifth day, 5% (w/v) Diaion HP-20 resin (Sigma-Aldrich) was added to the culture. The resin was harvested by centrifugation (3900 rpm, 10 min) and extracted with 20 mL of acetone under sonication for 15 min. The extracts were dried under reduced pressure and dissolved in methanol for LC-MS analysis.

LC-MS measurements were carried out on an Ultimate 3000 LC system (Thermo Fisher) coupled to an AmaZonX (Bruker) electro-spray ionization (ESI) mass spectrometer. Separation was achieved on a C18 column (ACQUITY UPLC BEH, 130Å, 1.7 µm particle size, 2.1 × 100 mm, Waters) at a flow rate of 0.4 mL/min at 40 °C, using acetonitrile and Milli-Q water supplemented with 0.1% (v/v) formic acid in a gradient ranging from 5 to 95% acetonitrile over 16 mins. HPLC-ESI-QTOF-MS analyses were conducted on an Ultimate 3000 LC system (Thermo Fisher) coupled to an Impact II QTOF mass spectrometer (Bruker). Separation was achieved on a C18 column (ACQUITY UPLC BEH, 130 Å, 1.7 µm particle size, 2.1 mm × 100 mm, Waters) at a flow rate of 0.4 mL/min at 40 °C, using acetonitrile and Milli-Q water supplemented with 0.1% (v/v) formic acid in a gradient ranging from 5 to 95% acetonitrile over 16 mins. Data were acquired in positive mode at a scan range between 100 to 1200 *m*/*z* to detect atropopeptides. The software DataAnalysis 4.3 (Bruker) was used to evaluate the measurements.

#### Large-scale production and purification of atropopeptides

For large-scale production of jumorubin, 6 seed cultures of Streptomyces strains carrying pUWL201-OriT-*jum* were prepared by fermentation in a culture tube containing 5 mL TSB medium with 50 µg/mL apramycin at 30 °C, 200 rpm, for 4 days. After incubation, 1 mL of the seed cultures was used to inoculate (6 × 100 mL) TSB medium containing 50 µg/mL apramycin in 500 mL Erlenmeyer flasks. After 3 days, 100 mL was used to inoculate (6 × 1000 mL) TSB medium containing 50 µg/mL apramycin in a 5000 mL Erlenmeyer flask. The cultures were incubated at 30 °C, 180 rpm, for 4 days. 5% (w/v) Diaion HP-20 resin (Sigma-Aldrich) was added to the culture and incubation was continued for one day. The Diaion HP-20 resin in the cultures was recovered by filtration using a metal sieve (40 mesh). The Diaion HP-20 resin was subsequently washed with $H_2O$ and then extracted twice with one culture volume of acetone. The crude extract was dried under reduced pressure.

The obtained crude extracts were fractionated by preparative chromatography (Büchi Pure C-850 Flash/Prep) using water and acetonitrile supplemented with 0.1% (v/v) formic acid as mobile phases A and B, respectively. The crude extract samples were purified using a Xbridge Prep C18 column (5 µm particle size, 250x19 mm, Waters) and eluted with a flow rate of 20 mL/min and a gradient from 10 to 45 % solvent B over 40 minutes. Eluting compounds were detected with a UV-detector (254–400 nm). Fractions were screened for atropopeptides by LC-MS analysis, and the fractions containing the desired compounds were dried under reduced pressure, re-dissolved in methanol and processed further by semi-preparative HPLC on an Agilent 1260 Infinity II UV– vis system, equipped with a Phenyl-Hexyl column (100 Å, 5 µm particle size, 250x10 mm, Phenomenex). Solvents used were Milli-Q water and acetonitrile as mobile phases A and B, respectively. Elution was achieved with a program containing three isocratic steps, 25% solvent B for 10 minutes, 28% solvent B for 20 minutes and 35% solvent B for 2 minutes using a flow rate of 5 mL/min. After LC-MS analysis of the fractions, fractions containing the compound of interest were collected, dried under reduced pressure and re-purified on the same semi-preparative HPLC system. A gradient was set from 30 to 39% solvent B over 30 min and then 100% solvent B for 5 min, giving a total run time of 35 min, with a flow rate of 3 mL/min. The purified compound was then subjected to NMR analysis.

For large-scale production of varsitalin, seed cultures of *Streptomyces* strains carrying pUWL201-OriT-SvarCore1R were prepared by fermentation in a culture tube containing 5 mL TSB medium with 50 µg/mL apramycin at 30 °C, 200 rpm, for 2-3 days. After incubation, 1 mL of the seed culture was used to inoculate (6 × 100 mL) TSB medium containing 50 µg/mL apramycin and 5% (w/v) Diaion HP-20 resin (Sigma-Aldrich) in 500-mL Erlenmeyer flasks. After 3 days, 100 mL was used to inoculate (6 × 1000 mL) mL TSB medium containing 50 µg/mL apramycin and 5% (w/v) Diaion HP-20 resin (Sigma-Aldrich) in a 5000-mL Erlenmeyer flask. The cultures were incubated at 30 °C, 180 rpm, for 4-6 days. The Diaion HP-20 resin in the TSB cultures was recovered by filtration with Miracloth (Milipore, MA, USA). The Diaion HP-20 resin was subsequently washed with $H_2O$ and then extracted twice with one culture volume of acetone. The crude extract was dried under reduced pressure. The obtained crude extracts were dissolved in DMSO and purified by preparative HPLC on an Agilent 1260 Infinity II UV–vis system, equipped with a XBridge BEH C18 OBD Prep Column (130Å, 10 µm particle size, 30 mm x 250 mm, Waters). Solvents used were Milli-Q water and acetonitrile supplemented with 0.1% (v/v) formic acid as mobile phases A and B, respectively. A gradient was set from 10 to 45% solvent B over 35 min and then 100% solvent B for 5 min, giving a total run time of 40 min, with a flow rate of 20 mL/min. The fractions containing the desired compounds were dried under reduced pressure, redissolved in DMSO and processed further by semi-preparative HPLC on an Agilent 1260 Infinity II UV–vis system, equipped with a Luna Phenyl-Hexyl column (100 Å, 5 µm particle size, 250 x 4.6 mm, Phenomenex). Solvents used were Milli-Q water and acetonitrile supplemented with 0.1% (v/v) formic acid as mobile phases A and B, respectively. A gradient was set from 20 to 60% solvent B over 30 min and then 100% solvent B for 5 min, giving a total run time of 35 min, with a flow rate of 3 mL/min. Fractions containing the desired compounds were subjected to LC-MS to check for purity. The purified compounds were weighed and redissolved in DMSO-$d_6$. After that, the purified compounds were subjected to NMR analysis (1D and 2D NMR).

For the large-scale fermentation of *S. albus*/pUWL201-OriT-*erm*, 6 L cultures was prepared as follows: the spores collected from one MS agar plate was incubated into a 5 L Erlenmeyer flask containing 1 L of the TSB medium with apramycin (50 µg/mL) and incubated on a rotary shaker (180 rpm) at 30 ºC for 6 days. 5% (w/v) of sterilized resin (Diaion HP-20) was added into each flask after 4 days of incubation, and the flasks were incubated for another 2 days. 4 L cultures was prepared as follows: the spores collected from one MS agar plate were inoculated into five 1 L Erlenmeyer flasks containing 200 mL of the TSB medium with apramycin (50 µg/mL). The incubation conditions were the same as described above. The resins from the 10 L cultures were harvested by filtration through a metal sieve (40 mesh). The harvested resins were washed with water, transferred to a separatory funnel and eluted with 5 L of acetone. After evaporation of the organic solvents under reduced pressure, the crude extracts were subjected to normal phase silica gel column chromatography (230–400 mesh) and eluted with $CHCl_3$/$CH_3OH$ (1:0, 97:3, 95:5, 90:10, 8:1, 4:1, 2:1, 0:1, v/v, 600 mL) to yield 8 fractions (Frs.1-8). Fr.6 and Fr.7 were combined and separated by Sephadex LH-20, eluted with $CHCl_3$/$CH_3OH$ (1:1, v/v) to obtain 4 sub-fractions (Fr.1.1 to Fr.1.4). Fr.1.2 was further purified by semi-preparative HPLC using a reverse-phase column (Luna phenyl-hexyl, 250×4.6 mm, 5 µm particle size, Phenomenex) with UV detection at 300 nm to afford compound **2** (3.2 mg) using the following gradient: solvent system (solvent A, water supplementing with 0.1% formic acid; solvent B, acetonitrile); 30 % B−39 % B (0-25 min), 39% B−100% B (25-26 min), 100% B (26-30 min), 100% B−30% B (30-31 min), 30% B (31-35 min); flow rate at 2.5 mL/min.

For large-scale production of laurentirubin B, seed cultures of *Streptomyces albus* strains carrying pUWL201-OriT-Precursor-laurentii were prepared by fermentation in a culture tube containing 5 mL TSB medium with appropriate antibiotic(s) at 30 °C, 200 rpm, for 2-3 days. After incubation, 1 mL of the seed culture was used to inoculate (6 × 100 mL) TSB medium containing appropriate antibiotic(s) and 5% (w/v) Diaion HP-20 resin in 500 mL Erlenmeyer flasks. After 3 days, 100 mL was used to inoculate (6 × 1000 mL) mL TSB medium containing appropriate antibiotic(s) and 5% (w/v) Diaion HP-20 resin in a 5000 mL Erlenmeyer flask. The cultures were incubated at 30 °C, 180 rpm, for 4-6 days. The Diaion HP-20 resin in the cultures was recovered by filtration with Miracloth. The Diaion HP-20 resin was subsequently washed with $H_2O$ and then extracted twice with one culture volume of acetone. The crude extract was dried under reduced pressure. The obtained crude extracts were dissolved in DMSO and then purified by preparative HPLC on an Agilent 1260 Infinity II UV–Vis system, equipped with a XBridge BEH C18 OBD Prep Column (130Å, 10 µm particle size, 30 mm x 250 mm, Waters). Solvents used were Milli-Q water and acetonitrile supplemented with 0.1% (v/v) formic acid as mobile phases A and B, respectively. A gradient was set from 10 to 45% solvent B over 35 min and then 100% solvent B for 5 min, giving a total run time of 40 min, with a flow rate of 20 mL/min. The fractions containing the desired compounds were dried under reduced pressure, redissolved in DMSO and processed further by semi-preparative HPLC on an Agilent 1260 Infinity II UV–vis system, equipped with a Luna Phenyl-Hexyl column (100 Å, 5 µm particle size, 250 x 4.6 mm, Phenomenex). Solvents used were Milli-Q water and acetonitrile supplemented with 0.1% (v/v) formic acid as mobile phases A and B, respectively. A gradient was set from 20 to 60% solvent B over 30 min and then 100% solvent B for 5 min, giving a total run time of 35 min, with a flow rate of 3 mL/min. Fractions containing the desired compounds were dried under reduced pressure. The purified compounds were weighed, redissolved in DMSO-$d_6$ and subjected to NMR analysis.

### Structure elucidation

Varsitalin B1 (**3**) was isolated as a yellow powder. Its molecular formula was determined to be $C_{42}H_{46}N_8O_8$ based on a protonated ion at *m/z* 791.3540 (calcd. for $C_{42}H_{47}N_8O_8^+$, 791.3517, Δ +2.9 ppm) in HR-ESI-QTOF-MS data (Figure S16). Analysis of the $^1$H and $^{13}$C NMR spectra (Figure S17 and S18), with the aid of the HSQC spectrum (Figure S19), revealed the presence of 40 carbons including four carbonyl carbons, 11 sp$^2$ protonated carbons, nine sp$^2$ unprotonated carbons, a quaternary carbon, seven methines, six methylenes, and two doublet methyls. These signals and signals for exchangeable NH protons ($\delta_H$ 7.83, 7.21, 5.89) are characteristic for peptides (Table S2). The $^1$H–$^1$H COSY spectrum (Figure S20) showed a constituted spin system of signals at $\delta_H$ 7.21, 3.87, 1.57, 1.39, 1.33, 0.76, and 0.74, indicating the presence of a Leu residue. The typical AMPX spin system ($\delta_H$ 7.09, 6.78, 6.72, 6.64) (Table S2) and the HMBC correlations from H-9 to C-11 and C-12 and from H-11 to C-8 (Figure S21) indicated the presence of a Trp residue (Trp2) that forms pyrroloindoline by a C–N bond between C-11 and N-8 (Figure S23). Another constituted spin system ($\delta_H$ 7.83, 4.00, 2.36, 2.32, 1.90. 1.64) and the core peptide sequence obtained from genomic data suggested the presence of a Gln residue. The presence of C-28-substituted Tyr was determined by HMBC correlations from H-25 to C-23, C-26, C-27, and C-31, from H-27 to C-29 and C-31, from H-30 to C-26 and C-28, and from H-31 to C-29 (Figure S23). Furthermore, C-39-substituted Trp (Trp1) was determined based on COSY correlations of H-33/H-34 and H-40/H-41 as well as HMBC correlations from H-36 to C-37 and C-42, from H-41 to H-37 and H-39, and from H-38 to C-40 and C-42. The amino acid sequence of **3** was determined to be $NH_2$-Trp1-Tyr-Gln-Trp2-Leu-$CO_2H$ based on the COSY and HMBC correlations (Figure S23), which were consistent with the amino acid sequence of the core peptide from *svaA2*. The C–C bond between C-39 of Trp-1 and C-10 of Trp-2 was determined by HMBC correlations from H-38 and H40 of Trp-1 to the quaternary carbon C-10 of Trp-2. In addition, the NOESY correlations of H-27/H-36 and H-27/H-33 (Figure S22, S23) suggested the presence of a C–N bond between N-36 of Trp-1 and the quaternary carbon C-28 of Tyr. The stereochemistry of *α*-carbons for each amino acid was deduced as L-configured based on the genomic data. The NOESY correlations of H-8/H-40, H-9/H40, H-9/H-41, and H-11/H-38 (Figure S23) suggest the relative configuration of the pyrroloindoline moiety to be (10*S**,11*R**) and the ansameric configuration[31] of the bicyclic macrolactam ring in **3** to be $P_{ansa}$. The ansameric configuration $P_{ansa}$ of **3** is the same as that of “bridge-above”-type tryptorubin A.[32]

Varsitalin B2a (**4a**) was isolated as an orange powder. Its molecular formula was determined to be $C_{42}H_{48}N_8O_8$ based on a protonated ion at *m/z* 793.3655 (calcd. for $C_{42}H_{49}N_8O_8^+$, 793.3668, Δ −1.6 ppm) in HR-ESI-QTOF-MS data (Figure S26). The $^1$H and 2D NMR spectra (Figures S27–S29, S31, S32) showed similarity to those of **3** except for the presence of the AA′XX′ spin system ($\delta_H$ 6.95, 6.63) for a Tyr residue, a secondary amine proton signal ($\delta_H$ 7.24) for Trp-1, and an amide proton signal ($\delta_H$ 6.95) for Trp-2, and the absence of a signal for H-11 of Trp-2. These differences suggested that **4a** is a peptide analog of **3** without the pyrroloindoline moiety at Trp-2 and a C–N bond between N-36 of Trp-2 and C-28 of Tyr. The HMBC correlations from H-38 and H-40 to C-11 (Figure S30) indicated the presence of a C–C linkage between C-39 of Trp-1 and C-11 of Trp-2. The presence of this C–C bond was further confirmed by NOESY correlations of H-8/H-38, H-8/H-40, H-9/H-38, H-9/H-40, and H-9/H-41 (Figure S33, S34). The stereochemistry of *α*-carbons for each amino acid was deduced as L-configuration based on the genomic data. The axial chirality between C-39 of Trp-1 and C-11 of Trp-2 was not determined in this study.

Embyscamide (**5**) was obtained as a white powder. Its molecular formula was determined to be $C_{37}H_{40}N_6O_5$ (*m/z* 649.3132 [M + H]$^+$, calcd. for $C_{37}H_{41}N_6O_5^+$, 649.3133, Δ −0.15 ppm) by HR-ESI-QTOF-MS data (Figure S36), suggesting the index of hydrogen deficiency to be 21. Analysis of the $^1$H and $^{13}$C NMR spectra, with the aid of the HSQC spectrum (Table S4, Figure S37–S40), revealed the presence of 37 carbons assignable to four carbonyl carbons, eight sp$^2$ nonprotonated carbons, 14 sp$^2$ methines, five sp$^3$ methines, four sp$^3$ methylenes, and two methyls. These signals and signals for exchangeable NH protons ($\delta_H$ 8.04, 7.81, 7.35) are typical for peptides. The presence of a Phe residue was determined based on the $^1$H–$^1$H COSY correlations between H-23/23′ and H-24 and the HMBC correlations of H-22/H-24, H-20/H-22, and H-20/H-21 (Figure S44). A continuous spin system in the aromatic region ($\delta_H$ 7.66, 7.52, 7.19, 7.12), a singlet signal for an aromatic proton ($\delta_H$ 7.85), and unequivalent methylene signals ($\delta_H$ 3.24, 3.15) suggested the presence of a Trp residue (Trp-2). The $^1$H–$^1$H COSY spectrum (Figure S41) showed another constituted spin system region including resonances for two terminal methyls ($\delta_H$ 7.81, 4.06, 1.68, 1.66, 1.36, 0.94, 0.91), indicating the presence of an Ile residue. Furthermore, the C-34-substituted Trp (Trp-1) was assigned based on the $^1$H–$^1$H COSY correlations of NH-29/H-29 and H-31/H-32 along with HMBC correlations from H-27 to C-29 and C-30, and from H-31 to C-30, C-33, and C-35, from H-32 to C-34, and from H-33 to C-35 (Figure S44). The amino acid sequence of **5** was determined to be $NH_2$-Trp-Phe-Ile-Trp-$CO_2H$ based on the HMBC correlations from amino protons to carbonyl carbons (Figure S42). The stereochemistry of *α*-carbons for each amino acid was deduced as L-configured based on the genomic data. The HMBC correlation from H-5 of Trp-2 to a quaternary carbon C-34 of Trp-1 suggested the presence of a C–N bond between C-34 of Trp-1 and N-5 of Trp-2. This bond was further confirmed by the NOESY correlations between the H-10/H-33, H-5/NH-29 (Figure S43).

Laurentirubin B (**6**) was isolated as a yellow powder. Its molecular formula of $C_{46}H_{45}N_7O_8$ was determined by a protonated ion at *m/z* 824.3401 (calcd. for $C_{46}H_{46}N_7O_8^+$, 824.3402, Δ −0.1 ppm) in HR-ESI-QTOF-MS data (Figure S47). The $^1$H and 2D NMR spectra were akin to those of tryptorubin B[33] except for the absence of an aromatic proton H-17 at a Trp-2 and signals of a Tyr-2 residue detected at $\delta_H$ 7.45, 7.10, 7.00, and 6.58 (Table S5, Figures S48–S51 and S53). These NMR data combined with the genomic data suggested that the amino acid sequence of **6** is identical to those of tryptorubin B ($NH_2$-Trp-Tyr-Ile-Trp-Tyr-$CO_2H$), but the modification of the amino acid sequence is different. The formation of the pyrroloindoline moiety at a Trp-2 residue was assigned from the COSY correlation between NH-12 and a sp$^3$ methine H-12 as well as the HMBC correlations from H-12 to an *α*-carbon C-9, a methylene carbon C-10, and a quaternary carbon C-11. The HMBC correlation of H-10/C-41, H-12/C-41, H-40/C-11, and H-42/C-11 indicated the presence of a C–C bond between a quaternary carbon C-11 of Trp-1 and a nonprotonated sp$^2$ carbon C-41 of Trp-1 (Figure S53). The NOESY cross peak between H-38 at Trp-1 and H-29 at Tyr-1 suggested the presence of a C–N bond between N-38 of Trp-1 and C-30 of Tyr-1. The aryl ether bond between C-7 of Tyr-2 and C-17 of Trp-2 was deduced from the NOESY correlations of H-5′/H-10*α* and H-6/NH-12 (FIgure S54 and S55), as well as asymmetric $^1$H and $^{13}$C signals for Tyr-2 residue (C-5, C-5′, C-6, C-6′) and a presence of a

1,2,3-trisubstituted aromatic ring in the pyrroloindoline moiety. The stereochemistry of α-carbon for each amino acid residue was suggested as L-configuration based on the genomic data. Besides, the NOESY correlations of H-9/H-43, H-10/H-43, and H-12/H-40 (FIgure S54 and S55) indicated the relative configuration of a quaternary carbon C-11 and an aminal carbon C-12 as (11*S**,12*R**) and the ansameric configuration of a bicyclic macrolactam ring as $P_{ansa}$. To confirm the location of the aryl ether bridge and stereochemical configuration of the ansameric bismacrocycle, , theoretical $^{13}$C NMR chemical shifts were calculated for possible conformers $P_{ansa}$-**6**, $M_{ansa}$-**6**, along with a possible isomer $P_{ansa}$-**6a** that features an aryl ester bond between C-1 and C-17 instead of aryl ether bond between C-7 and C-17 using the GIAO method. The calculated $^{13}$C NMR chemical shifts of $P_{ansa}$-**6a** with the lowest values of the mean absolute error and mean squared error (Table S6) were more favorable than those of $M_{ansa}$-**6** and $P_{ansa}$-**6a**. Furthermore, the lowest energy conformer of $P_{ansa}$-**6** at the B3LYP/6-31G(d,p) level of theory (Figure S57) showed a rational geometry that was in agreement with the observed NOESY correlations in **6**, except for the NOESY correlation of H-9/H-43. These DFT calculations supported the NMR analysis-based assignment of the ansameric configuration as $P_{ansa}$, as well as the location of the aryl ether bond at C-7–C-17.

### DFT Calculations

3-dimensional structures for $P_{ansa}$-**6,** $P_{ansa}$-**6a,** and $M_{ansa}$-**6** were modeled in the Spartan '18 V1.4.5 modeling software. The models were minimized using molecular mechanics with the Spartan MMFF force field and subsequently subjected to conformational search using molecular mechanics again with the MMFF force field and using a Monte Carlo search method (specified by the option SEARCHMETHOD=MC). For each structure, the ten conformers with the lowest energy, as determined by MMFF, were selected for DFT calculations. All DFT calculations were performed with Gaussian 16 using the B3LYP function with a 6-31G(d,p) basis set and in the gas phase. Each conformer was energy minimized in Gaussian and checked for convergence with a frequency calculation. If the minimization had not converged, we repeated the minimization until convergence. We then performed an NMR GIAO calculation on the minimized structure. We used regression coefficients described by Konstantinov and Broadbelt[34] to calculate the predicted chemical shifts for all carbons. We then averaged over the ten predicted spectra for each structure using a Boltzmann-weighted average. We determined and visualized the distances in the lowest energy structures corresponding to key NOESYs using Pymol.
